# Grey and white matter metrics demonstrate distinct and complementary prediction of differences in cognitive performance in children: Findings from ABCD (N= 11 876)

**DOI:** 10.1101/2023.03.06.529634

**Authors:** Lea C. Michel, Ethan M. McCormick, Rogier A. Kievit

**Affiliations:** Cognitive Neuroscience Department, Radboud University Medical Center, 6525 GA Nijmegen, The Netherlands; Methodology and Statistics, Institute of Psychology, Leiden University, Leiden, Netherlands; Department of Psychology and Neuroscience, University of North Carolina, Chapel Hill, United States

## Abstract

Individual differences in cognitive performance in childhood are a key predictor of significant life outcomes such as educational attainment and mental health. Differences in cognitive ability are governed in part by variations in brain structure. However, studies commonly focus on either grey or white matter metrics in humans, leaving open the key question as to whether grey or white matter microstructure play distinct or complementary roles supporting cognitive performance.

To compare the role of grey and white matter in supporting cognitive performance, we used regularized structural equation models to predict cognitive performance with grey and white matter measures. Specifically, we compared how grey matter (volume, cortical thickness and surface area) and white matter measures (volume, fractional anisotropy and mean diffusivity) predicted individual differences in cognitive performance. The models were tested in 11,876 children (ABCD Study, 5680 female; 6196 male) at 10 years old.

We found that grey and white matter metrics bring partly non-overlapping information to predict cognitive performance. The models with only grey or white matter explained respectively 15.4% and 12.4% of the variance in cognitive performance, while the combined model explained 19.0%. Zooming in we additionally found that different metrics within grey and white matter had different predictive power, and that the tracts/regions that were most predictive of cognitive performance differed across metric.

These results show that studies focusing on a single metric in either grey or white matter to study the link between brain structure and cognitive performance are missing a key part of the equation.

**Significance Statement:** This paper enriches the recent debates on the challenges of linking variation in brain structure to phenotypic differences (Marek et al., 2022). We demonstrate that using latent variables (to improve power), structural equation modelling (to allow greater flexibility in linking brain to behaviour), and by simultaneously incorporating multiple measures of grey and white matter in a large sample, we demonstrate relatively strong and robust brain-behaviour associations, which highlight the complementarity of grey and white matter metrics in predicting cognitive performance as well as the importance of incorporating the full complexity of these associations over 1-to-1 linkages. This finding should lead researchers to consider integrating both grey and white matter measures when demonstrating a more comprehensive picture of brain-cognition relationships.

## Introduction

The field of cognitive neuroscience is premised on the hypothesis that differences in cognitive performance can be understood in part by studying differences in brain structure and function (Basten et al., 2015). Although much is known about the relation between grey and white matter structure and cognitive performance (Tamnes, Østby, Walhovd, et al., 2010; Tamnes, Østby, Fjell, et al., 2010; Magistro et al., 2015; Muetzel et al., 2015; Schnack et al., 2015; Deoni et al., 2016), less is known about how the structure of the two tissues *together* explain differences in cognitive performance.

One way to look at this challenge is to consider grey and white matter as two sides of the same coin. At the cellular level, they are both composed of neurons, although different parts (cell bodies, dendrites, synapses and axons for grey matter and (un)myelinated axons for white matter), glial cells and vasculature (Wandell, 2016; Purves et al., 2019). Moreover, recent findings observe even more overlap, namely that differences in one tissue (white matter myelination) may affect the other tissue (cortical thickness) (Natu et al., 2019), suggesting they may capture the same neurobiological properties using different techniques.

A counter argument would be that grey and white matter are two distinct brain properties with distinct mechanistic roles. Twin study reports that grey and white matter volume shared 68% heritability, suggesting both overlapping and distinct genetic mechanisms (Baaré et al., 2001) and recent attention has focused on their different transcriptome patterns, highlighting the differentiation of their cells and their specific functional roles (Mills et al., 2013). Studies have shown the importance of both dendritic network and myelinated axons to provide faster and more efficient cognitive performance (Kail, 1997; Tamnes et al., 2012; Rolls & Deco, 2015), thus illustrating the potentially complementary roles of grey and white matter to explain differences in cognitive performance.

Beyond the conventional classification of grey and white matter, it is important to consider the heterogeneity within these tissues. Each specific region/tract *within* grey and white matter may offer unique or shared information in predicting cognitive performance, both within a single metric but also across diverse metrics. Understanding the intricate interplay between these regions/tracts and metrics can provide a more comprehensive understanding of brain-behavior associations.

In summary, there is compelling evidence to suggest that grey and white matter play complementary roles explaining differences in cognitive performance. To date, few studies investigated the combining effects of grey and white matter differences on cognitive performance and so far the literature point towards a complementary roles of the two tissues (Østby et al., 2011; Kievit et al., 2014; Ritchie et al., 2015).

The causes of this paucity are manifold. First, study designs tend to focus on either classical MRI sequences (T1,T2,T2*) or diffusion weighted imaging data. These are sequences with distinct scanner demands and require dedicated analysis expertise, which often leads to papers focusing on one tissue. Moreover, the standard implementation in neuroimaging software, where a brain region/voxel is the *outcome* of a regression equation, makes it considerably more challenging to implement models where multiple brain metrics predict a single phenotypic outcome (e.g. cognitive performance). Finally, the high number of predictors of such models might hinder their feasibility, decrease the statistical power and generate false positives.

To overcome these challenges, we investigated the unique predictive roles of grey and white matter metrics in predicting cognitive performance in childhood. By conducting a multimodal analysis in a regularized structural equation model in a large sample, we overcome many existent weaknesses in previous work, and it allowed us to maximize the likelihood of distinguishing the competing hypotheses of interest. In addition, we will explore the regional distribution of these associations within and across metrics, as well as identify which specific metrics in grey and white matter are more influential in predicting cognitive performance.

This study provides new insights into the complementary information provided by grey and white matter in supporting cognitive performance.

## Methods

### Participants

The ABCD study (https://abcdstudy.org/) is an ongoing longitudinal study across 21 data acquisition sites enrolling 11 876 children from 9 years old to 16 years old. For more information on ABCD protocols and inclusion/exclusion criteria see Volkow and colleagues (Volkow et al., 2018).

This paper analysed the baseline sample (9-11 years old) from release 4.0 (https://abcdstudy.org/; http://doi.org/10.15154/1523041) that include a sample of 11,876 children. Data entry outliers for 10 participants in the Little Man task were replaced with NA and included in the full information maximum likelihood (FIML) estimation. We included participants with partial data across cognitive tasks and the neuroimaging data in our models.

Given the complexity of the analysis and the a priori challenges associated with high dimensional regularized structural equation modeling approaches, a fully preregistered analysis was not possible. However, to ensure robustness of our findings, we divided the sample into two subsets, allowing us to balance exploratory model optimisation and validation in a non-overlapping sample. In our study, a random sample of 15% of the data were used to optimize the model-building and estimation steps, and the other 85% of the sample was used as a validation sample (Srivastava, 2018) (Figure 1). The set of regions used in the validation sample was derived from the best predictive regions identified in regularized models estimated using the model-building sample (Schwarz, 1978) (the regularisation techniques will be explained more in the Experimental Design and Statistical Analysis section). This strategy allows us to jointly optimise robustness and flexibility in cases where model estimation, adaptation and convergence are non-trivial, whilst maintaining sufficient power in the validation set to be sufficiently well powered.

**Figure 1:**
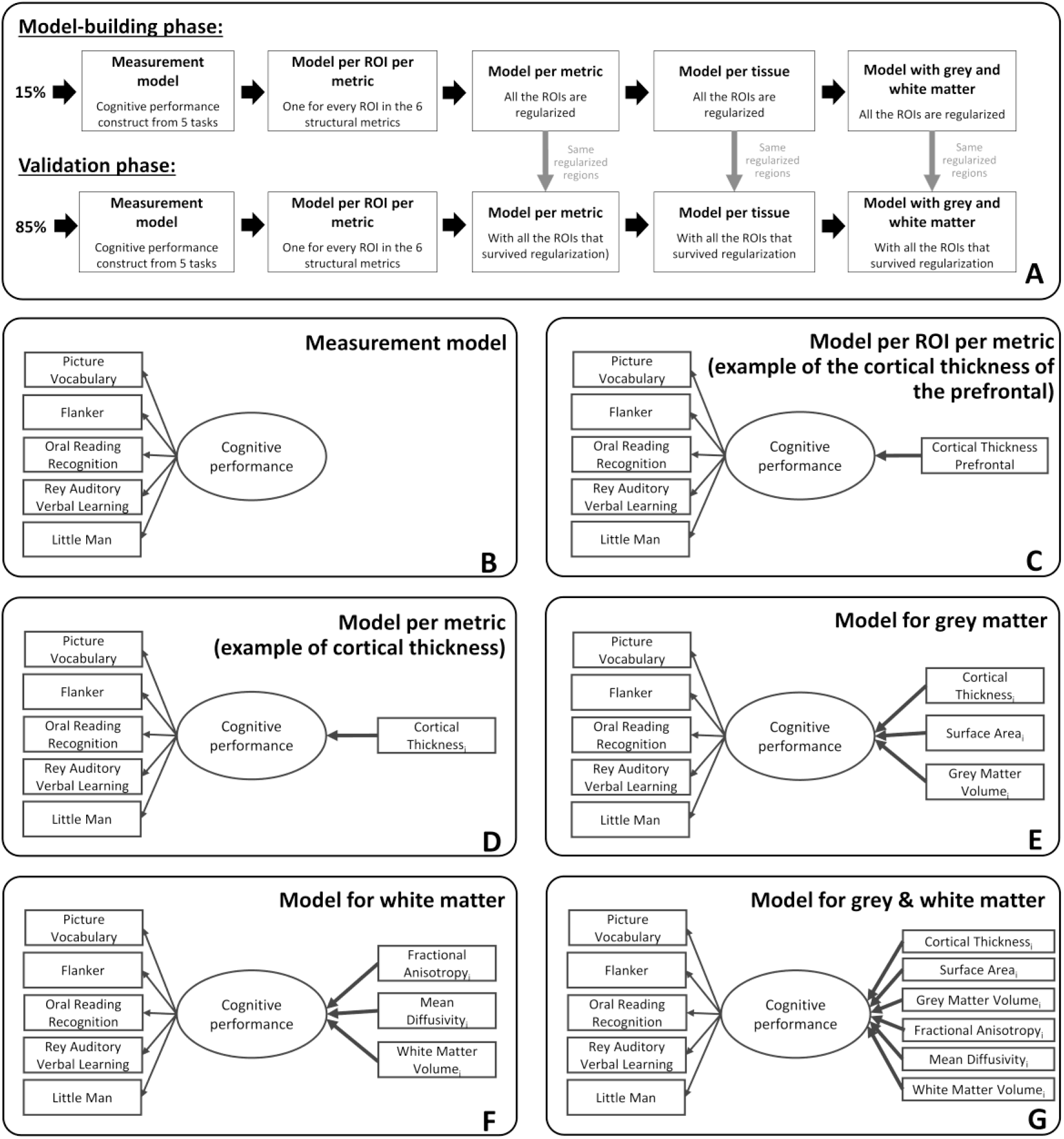
(A) Steps followed during the model-building and the validation phases of the study. The regions of interest fitted into the models of the validation phases resulted from an initial regularisation made in the model-building phase. (B) Measurement model of cognitive performance based on five cognitive tasks. (C) Example of a MIMIC model path diagram of a model per ROI per metric where the one metric of one region of interest (cortical thickness of the prefrontal) predict cognitive performance. (D) Example of a MIMIC model path diagram of a model per metric where all regions of interest of one metric (here cortical thickness) that survived regularization predict cognitive performance. (E) MIMIC model path diagram for the grey matter model with cortical thickness, surface area and grey matter volume predicting cognitive performance. i represents the regions of interest from each metric in grey matter that survived regularization in the model and can go from 1 to 34 (as we use the Desikan-Killiany atlas and average bilaterally). (F) MIMIC model path diagram for the white matter model with fractional anisotropy, mean diffusivity and white matter volume predicting cognitive performance. j represents the regions of interest from each metric in white matter that survived regularization in the model and can go from 1 to 20 (as we use the AtlasTrack atlas and average bilaterally). (G) Path diagram for the grey and white matter model with cortical thickness, surface area, grey matter, fractional anisotropy, mean diffusivity and white matter volume of the regions that survived regularization predicting cognitive performance.

The model-building sample consisted of 1,781 children (49.1% female, mean age=9.9, SD=0.6, range=8.9-11.1) and the validation sample included 10,095 children (47.6% female, mean age=9.9, SD=0.6, range=8.9-11).

### Cognitive performance

In our modelling approaches, we want to focus on construct capturing a broad sampling of cognitive ability. To this end, we use the same procedure as identified in Sauce et al. (2022), by selecting five cognitive tasks from the NIH Toolbox Cognition Battery were selected: Picture Vocabulary, Flanker, Oral Reading Recognition, Rey Auditory Verbal Learning and Little Man (Sauce et al., 2022). They measure respectively language vocabulary knowledge, attention and inhibitory control, reading decoding skill, verbal learning and memory, visuospatial processing flexibility and attention. All of the tasks were administered using an iPad with support or scoring from a research assistant where needed. For more information on each task, see Luciana et al. (2018).

Next we specified a confirmatory factor model which posits a single latent factor for cognitive performance, reducing measurement error and increasing precision. It enables the model to compute the common variance between these five tasks and thus to have a more accurate representation of cognitive performance (Figure 1B). The measurement model has already been validated in another study, for more information on the rationale for the choice of cognitive tasks and the measurement model see Sauce and colleagues (Sauce et al., 2022). Note, the model chosen here should not be interpreted as a commitment to a strong ‘causal g’ (see Kievit et al., 2017), but rather as a way to specify a broad cognitive factor which will maximize our statistical power to capture the patterns of interest. A fruitful avenue of future work will be to examine the degree (or lack of) overlap in the neural mechanisms predicting each individual cognitive domain – a goal beyond the present paper.

### Brain structure measures

MRI and DTI scans were collected across the 21 research sites using Siemens Prisma, GE 750 and Philips 3T scanners. Scanning protocols were harmonized across sites and scanners. Full details of all the imaging acquisition protocols and the processing methods used in ABCD are outlined elsewhere (Casey et al., 2018; Hagler et al., 2019).

To determine the importance of grey and white matter, we chose six structural metrics available in the ABCD study: cortical thickness (CT), surface area (SA) and volume (GMV) for grey matter and fractional anisotropy (FA), mean diffusivity (MD) and volume (WMV) for white matter.

## Grey Matter

Grey matter measures were estimated from MRI scans with FreeSurfer (version 7.1.1).

The processing steps involved cortical surface reconstruction, subcortical segmentation, removal of non-brain tissue, Talairach transformation, segmentation of white matter and deep gray matter structures, intensity normalization, and surface deformation. The images were registered to a spherical atlas based on individual cortical folding patterns, and the cerebral cortex was parcellated into 34 regions per hemisphere with the Desikan-Killiany atlas (Desikan et al., 2006). The full pipeline can be found elsewhere (Hagler et al., 2019).

The FreeSurfer output includes volume, cortical thickness and surface area (Casey et al., 2018). Considering the challenges of dimensionality in our SEM, we decided to average every region of interest bilaterally (i.e. the brain was parcellated in 34 regions across the brain for each metric).

## White Matter

White matter structure can be measured through diffusion-weighted imaging (DWI) and structural MRI (Wandell, 2016). DWI is a technique that allows researchers to record the diffusion of water in the brain using diffusion tensor imaging (DTI) to model the diffusion within the axons and myelin sheath, thus to analyse the directionality of the white matter tracts (Le Bihan, 2003; Assaf & Pasternak, 2008). Structural MRI provides a measure of volume while DTI computes fractional anisotropy, mean diffusivity, radial diffusivity and axial diffusivity (AD).

In the ABCD study, diffusion measures were obtained using tabulated diffusion MRI data from a high angular resolution diffusion imaging (HARDI) sequence with multiple b-values. The images were corrected for head movement, eddy current distortions, B0 distortion, and gradient nonlinearity distortion. The resulting diffusion measures were used for subsequent analyses. The processing steps are described elsewhere (Hagler et al., 2019).

White matter measures include white matter volume, fractional anisotropy and mean diffusivity. The tracts were divided with the AtlasTrack atlas which created 37 regions of interest (17 regions per hemisphere and 3 regions aside) (Hagler et al., 2019). We followed a similar procedure as with the grey matter metrics and averaged every tract of interest bilaterally.

### Experimental Design and Statistical Analysis

Next, we specified the core questions investigated by our analyses.

- *Do the metrics of grey matter and white matter demonstrate complementary roles predicting cognitive performance?*
- *Do the different metrics in each tissue have unique predictive roles? If so, which metric is most important to predict cognitive performance?*
- *For each metric, do the different regions of interest have unique predictive roles?*
- *Do the same regions of interest have the strongest predictive power across different metrics in each tissue?*

Our focus will be on both the tissue scale (grey and white matter) and the metrics scale (cortical thickness, surface area, grey matter volume, fractional anisotropy, mean diffusivity and white matter volume). For each scale, we will study if the predictors of cognitive performance each have a unique role (i.e. there is no overlap in the information contributed by the different variables to predict cognitive performance) or if they have complementary roles (i.e. there is a partial or total overlap in the information contributed by the different variables to predict cognitive performance).

The study uses structural equation model approach to evaluate the hypothesis that brain structural metrics predict the latent variable, cognitive performance. Structural equation models, and more particularly MIMIC models (Multiple Indicator Multiple Cause) offer an effective way to model cognition as a latent variable and to estimate the contribution of multiple simultaneous hypothesized causes to explain individual differences in cognitive performance (Jöreskog & Goldberger, 1975; Kievit et al., 2014).

First, we estimated a series of structural equation models to test how each metric in each individual region/tract predicted cognitive performance (i.e., every region in all the six metrics have been computed independently). The key question of interest is the strength of the key parameter highlighted in bold (Figure 1C).

Next, to assess how each metric predicted cognitive performance we used a regularized structural equation modelling (SEM) approach (Jacobucci et al., 2019) which incorporates a penalty on key parameters of interest. Specifically, it allows us to have many regions/metrics simultaneously predict the outcome, with a penalty on the path estimates from brain metrics to the cognitive latent variable, that induces sparsity. Regularization is a method that imposes a penalty in order to decrease the complexity of the model while keeping the variables that are the most important in predicting the outcome. For instance, it allows us to include the cortical thickness measures from all 34 regions of interest as simultaneously predicting cognitive performance with a lasso penalty which pushes parameter estimates of small or absent effects to 0, and retains only those regions which contribute meaningfully in predicting the outcome for the validation sample. This accounts for the partial overlap in variance arising from the inherent way the metrics are measuring the same underlying structure. For instance, grey matter volume is a product of surface and cortical thickness. If grey matter volume functions as a deterministic transformation with no additional information beyond cortical thickness or surface area for cognitive performance, our regularization approach should ensure all redundant parameter estimates are pushed to 0. For example, if the cortical thickness, surface area and grey matter volume of the frontal pole each have large effects on cognitive performance and these effects are redundant (they include the same information to predict cognitive performance), then the measures with the smaller effects of the three will be pushed to 0.

We developed a procedure to select a subset of regions/tracts based on their predictive ability in the model-building sample (e.g., 15% of the total sample). The output of each regularized model estimation was a set of all the regions/tracts with a regularized beta different from zero, or considered ‘important’ in a regularized framework. These regions were entered into the models predicting cognitive performance in the validation sample (e.g., 85% of the total sample). We implemented this process across six models per metric (Figure 1D), two models per tissue (Figures 1E-F) and the model combining grey and white matter metrics (Figure 1G). The benefit of this approach is to have a parsimonious representation of the key regions which help predict cognitive performance in each metric. We also examined ‘full’ models combining every region of interest within a metric (e.g. the 34 regions for cortical thickness), within a tissue (e.g. the 102 regions for the three grey matter metrics and the 60 regions for the three white matter metrics) and within grey and white matter metric (e.g. the 162 regions across the six metrics).

To compare the predictive information of grey and white matter, three models were fitted: one model with the regions extracted from the regularization of the three grey matter metrics, one model with the regions extracted from the regularization of the three white matter metrics and a final model with the regions extracted from the regularization of the grey and white matter metrics (e.g., as displayed in Figure 1A, the regions were selected because they survived the regularisation in the model-building sample). For each model, we use a likelihood ratio test to examine whether the inclusion of a tissue (grey/white matter) or a metric within a tissue (e.g. cortical thickness) improves the model compared to a model where the paths corresponding to an additional metric/tissue are constrained to 0. For each final model, we extracted the (adjusted) R-square to assess the proportion of variance in cognitive performance explained by each model.

From the analyses, we can imagine our results being captured by one of three (simplified) scenarios:

- Grey and white matter metrics give the same, non-complementary information to predict cognitive performance. The models with only grey matter, only white matter or both will have a similar adjusted R-squared (Figure 2A), and model selection would favour a model with only one tissue.
- Grey and white matter metrics give fully distinct/complementary information to predict cognitive performance. Under this scenario, model selection would favour a model including both tissues and the joint R-squared would approximate the sum of the R-squared of each tissue in isolation (Figure 2B).
- Grey and white matter metrics give complementary, but partially overlapping, information to predict cognitive performance. In this case, model selection would favour a model with both tissues, and the joint R-squared will be higher than the one of the most predictive tissue, but lower than the sum of the R-squared of each tissue in isolation (i.e. the R-squareds are neither interchangeable nor fully additive) (Figure 2C).

**Figure 2:**
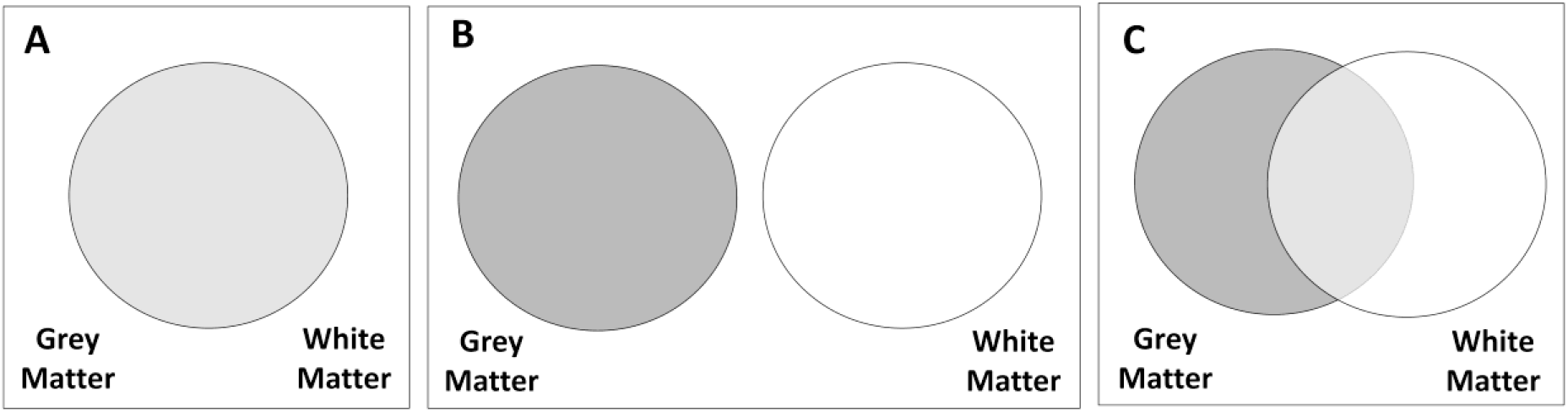
Hypotheses regarding the information provided by grey and white matter metrics. (A) Redundant, grey and white matter give the same information. (B) Independent, grey and white matter give completely different information. (C) Partially overlapping, grey and white matter give both similar and different information.

These analyses were replicated separately for male and female participants to examine the potential impacts of sex differences in brain structure for the findings. In an additional analysis, total intracranial volume (TIV) was added as a third predictor to explain the differences in cognitive performance. Accounting for TIV in the model is a contentious issue (Hyatt et al., 2020), as it modifies the question from whether it is the absolute size (or thickness, anisotropy,..) of a region that matters to predict cognitive performance, or the relative measure given a participant’s TIV.

Regularization has many strengths, as outlined before, but this comes at a specific cost, namely its attempt to build a simplified model. In other words, if two regions have almost the same predictive power, but are somewhat redundant in this regard, a regularized model is likely to push the parameter estimate of one of the two regions to 0 and ‘keep’ the other region instead. If we were to report the regional pattern as such, this may lead to an overinterpretation of the absence and presence of some regions. To examine the regional specificity of our findings, we conducted a repeated regularization procedure, iterated 1000 times using a different 15% sample on each occasion. Across these 1000 iterations, we computed the percentage of times each region survived regularization (i.e. when a given region had a non-zero estimate in the final model). This approach allowed us to assess the stability of each region/tract in providing unique information to explain variations in cognitive performance. Within each model (see Figure 1D-G), we calculated the percentage of instances in which a region/tract survived regularization. Regions/tracts with higher percentage values demonstrate a more consistent and distinct contribution to the prediction of cognitive performance. In the model employing a single metric (Figure 1D), this reflects the differential information provided by distinct regions/tracts within the same metric. In the models incorporating multiple metrics (Figure 1E-G), the likelihood of regions/tracts surviving regularization is lower if they provide redundant information across different metrics. Therefore, regions/tracts with a higher percentage not only demonstrate a unique predictive role across different regions/tracts but also across the various metrics.

We used the following guidelines to assess the good fit of the models: root mean square error of approximation (RMSEA) <0.05 (acceptable: 0.05–0.08), comparative fit index (CFI)>0.97 (acceptable: 0.95–0.97) and standardized root mean square residual (SRMR)<0.05 (acceptable: 0.05–0.10) (Schermelleh-Engel et al., 2003; Mueller & Hancock, 2008). We compared models fit using akaike information criterion (AIC) and bayesian information criterion (BIC). These parameters penalize models with a higher number of predictors, thus encouraging the selection of more parsimonious models that explain the data well while avoiding overfitting.

All analyses were carried out on the data of the first wave using R, version 4.1.0 (http://www.r-project.org/) and the lavaan package (Rosseel, 2012). All models were fit using Maximum Likelihood Estimation, with FIML to account for missing data and robust estimation with adjusted standard errors to deal with deviations from (multivariate) normality.

### Data and Code Accessibility

Data can be requested through [https://nda.nih.gov/], and the code to reproduce our analyses is available on [https://osf.io/ryskf/].

## Results

### Measurement model (validation, parameters, invariance)

To assess cognitive performance, we used the same measurement model built by Sauce and colleagues (Sauce et al., 2022) in a slightly different sample. Model estimates were highly similar.

The confirmatory factor model fitted the data well in the validation sample, x² = 156.404, degrees of freedom (df)= 5, p<0.001, RMSEA = 0.055 (0.00 - 0.070), CFI= 0.980, SRMR= 0.020. This result demonstrates that the common variance among all five cognitive tasks can be captured by one latent variable that we call here “cognitive performance”.

Oral Reading Recognition task and Picture Vocabulary task have the strongest standardized factor loadings (0.74 and 0.7 respectively). Rey Auditory Verbal Learning task, Little Man task and Flanker task are mildly predicted by the construct between 0.41 and 0.53. (Figure 3). The results show little change when we add age as a predictor of the latent variable.

**Figure 3:**
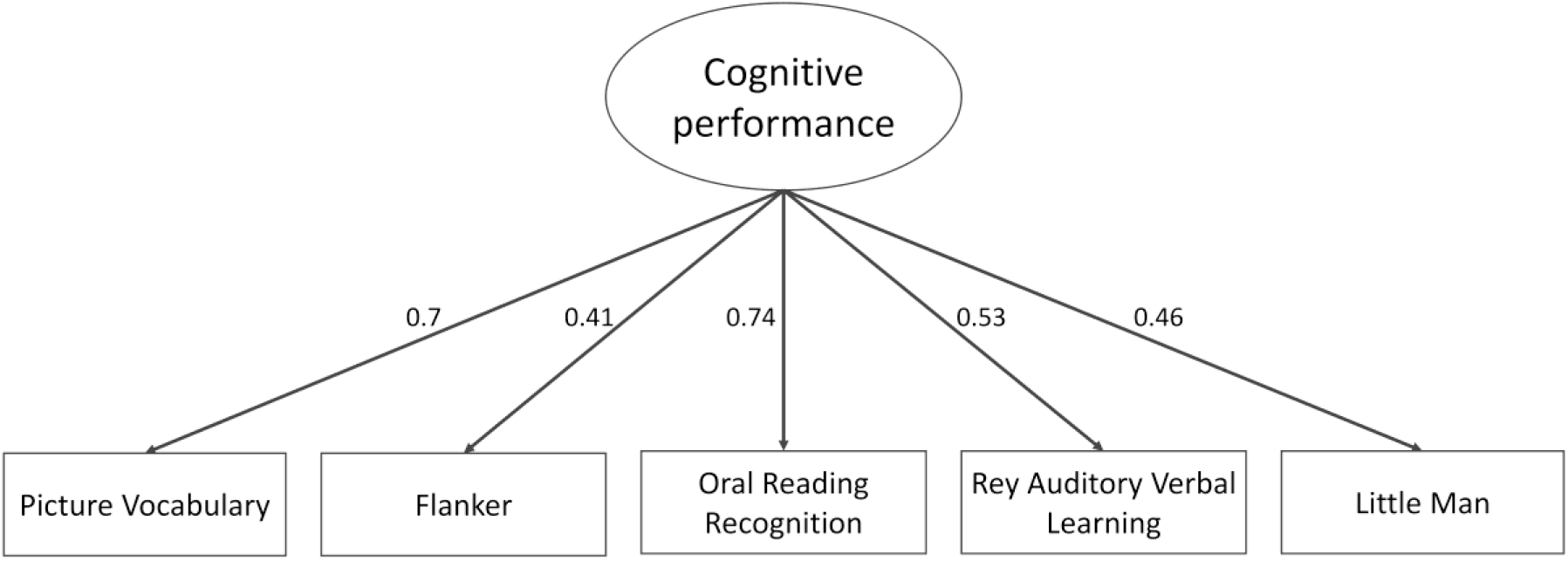
Path diagram of a measurement model of cognitive performance based on five cognitive tasks. All variables are standardized and significant.

### 1. Grey and white matter give both overlapping and unique information to predict cognitive performance

We fitted the full model including all regions/tracts that survived regularisation in grey & white matter metrics, and compared it to a model which includes the same predictors, but constrains either all grey or all white matter predictors to 0. This comparison allows us to test if the metrics in grey matter and white matter have complementary roles predicting cognitive performance.

The model with both grey and white matter fitted the data well (x² = 793.903, df= 205, p<0.001, RMSEA = 0.017 (0.016 - 0.019), CFI= 0.935, TLI=0.918 SRMR= 0.011) and showed the best performance among the three models (AICdiff > 265, BICdiff > 136 in favour of the model with both grey and white matter), and explained 19.0% of the variance in cognitive performance. The model with only grey matter explained 15.4% of the variance and the one with only white matter explained 12.4% of the variance (Table 1).

**Table 1:** Estimates of the models fit and the variance explained by the predictors within each model. The best model is shown in bold.

|  | Grey Matter model | White Matter model | Grey and White Matter<br>model |
| --- | --- | --- | --- |
| **R-square**<br>**(adjusted)** | 0.154 | 0.124 | **0.190** |
| **AIC** | 122941.1 | 123153.5 | **122675.3** |
| **BIC** | 123277.4 | 123389.6 | **123140.5** |

These findings demonstrate that grey and white matter bring both overlapping and unique information to predict the differences in cognitive performance in line with hypothesis C – ‘partially overlapping’ – in Figure 2.

These results were replicated in samples comprising exclusively of male (R-square adjusted for grey matter and white matter= 17.4%, for grey matter= 14.6% and for white matter= 10.8%) and also female participants (R-square adjusted for grey matter and white matter= 23.0%, for grey matter= 17.8% and for white matter= 15.9%).

In the model accounting for total intracranial volume, we also compared a model with only TIV as a predictor and a model with grey matter, white matter and TIV as a predictor instead of only grey and white matter. The model with grey matter, white matter and TIV explained the best the variance in cognitive performance (R-square adjusted for grey matter and white matter and TIV= 19.1%), while the model with only TIV explained 6.8%. The addition of TIV as a predictor did not improve the fit of the model, nor did its inclusion substantially altered the parameters estimates of the regions/tracts.

Figure 4 illustrates the overlap in information within grey and white matter metrics, as evident from the depicted contrast in the percentage of times one region/tract is consistently surviving regularization in a model with only the metrics on one tissue versus a model including the metrics of both grey and white matter. Despite this overlap, certain regions still reliably predict cognitive performance, highlighting the distinct information observed within both tissue types.

**Figure 4:**
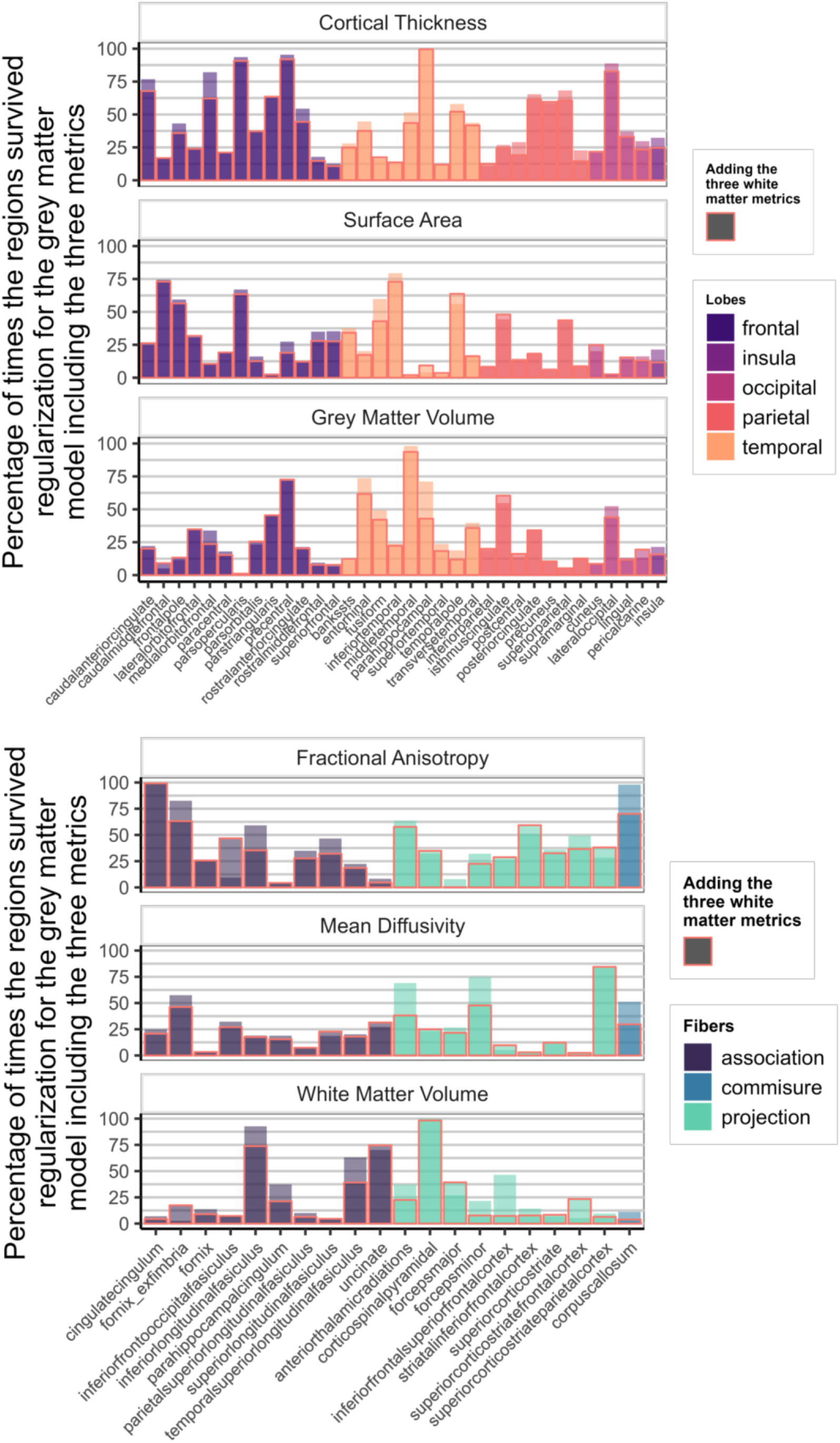
Percentage of times one region/tract in a metric survived regularization in the model with the three metrics of one tissue (top: grey matter, bottom: white matter) (Model E & F in Fig 1). The columns highlighted in red represent the percentage of survival for the region when the model includes the metrics of the other tissue (Model G in Fig 1)

### 2. Some metrics are more predictive of cognitive performance than others

Next, we investigated if the different metrics in each tissue have a unique predictive role, and thus if the choice of the metric was important when you want to predict cognitive performance. Comparing the predictive power of the different metrics, we plotted the standardized estimate model parameters of every region across metrics. Figure 5 shows the range of observed values in the path estimates for each metrics, within the grey matter metrics, surface area (range *β*^*std*^ = [0.092;0.265]) and volume (range *β*^*std*^ = [0.123;0.275]) were overall stronger predictors of cognitive performance compared with cortical thickness (range *β*^*std*^ = [-0.06;0.145]). This is somewhat surprising given the prominence of cortical thickness as the metric of choice in (developmental) cognitive neuroimaging studies of individual differences. Cortical thickness is commonly used both in healthy and case control studies, with 5 times more papers combining cortical thickness and cognition compared to the number of papers using grey matter volume and cognition since 2016 (N=2351 for cortical thickness and N=506 for grey matter volume in a PubMed search as of 30/10/2023)^1^.

**Figure 5:**
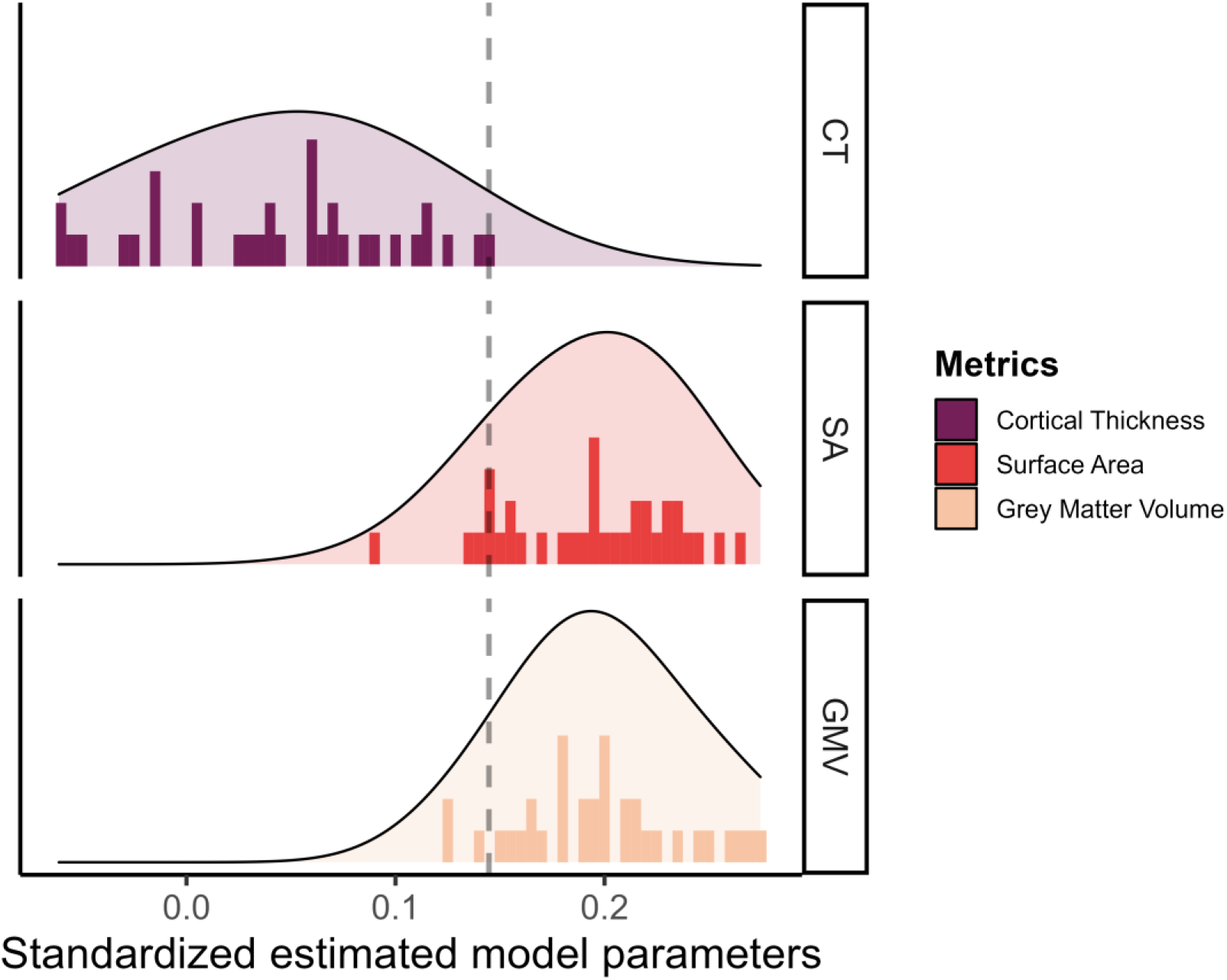
Standardized estimated model parameters on how one region of interest in one metric predicts cognitive performance in the models per ROI per metric for the three grey matter metrics.

The results for the white matter metrics is represented in Figure 6. Surprisingly, white matter volume is overall the strongest predictor of cognitive performance (range*β*^*std*^ = [0.169;0.303]), followed by fractional anisotropy (range *β*^*std*^ = [-0.010;0.142]) and mean diffusivity (range *β*^*std*^ = [-0.062;0.031]).

**Figure 6:**
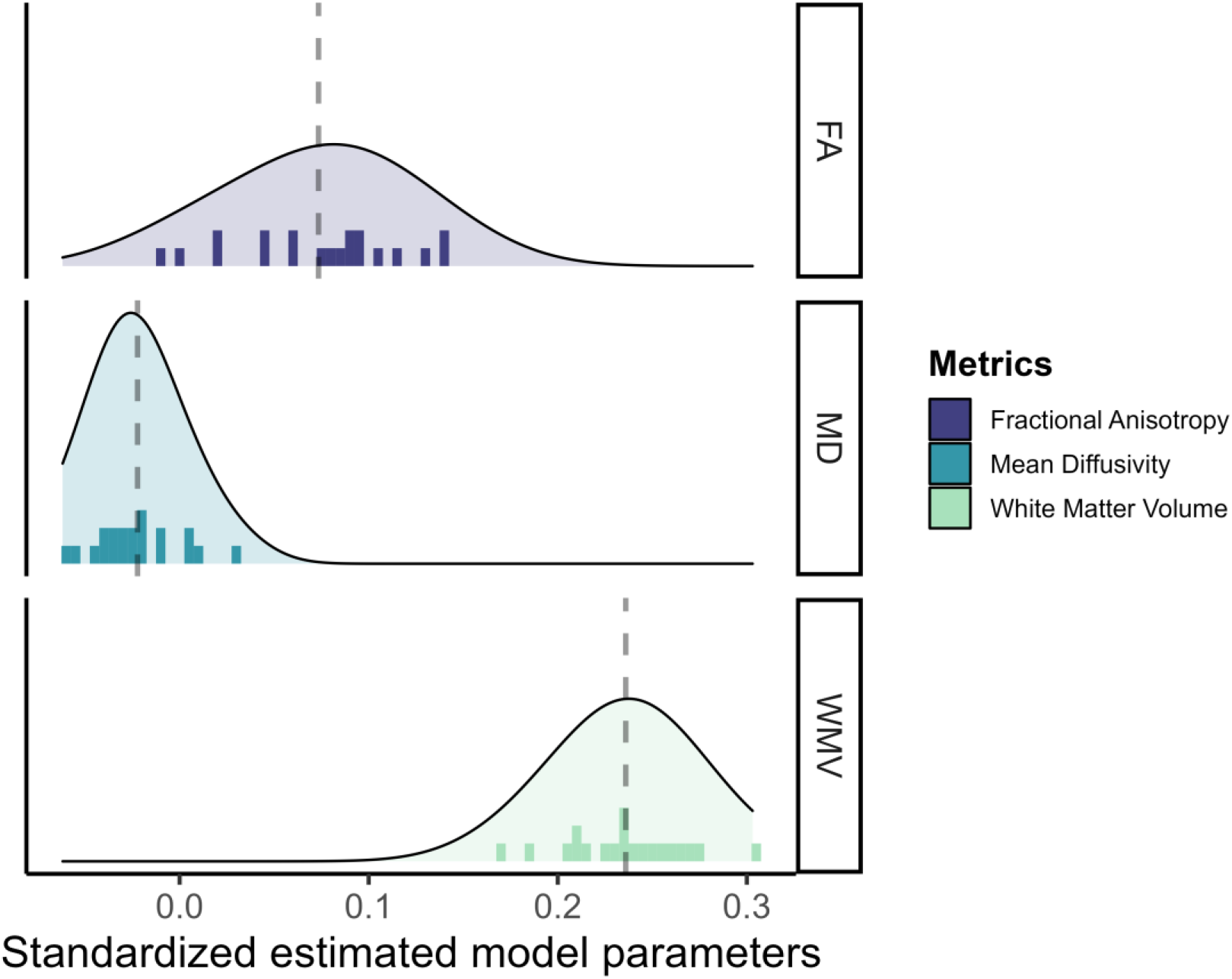
Standardized estimated model parameters on how one region of interest in one metric predicts cognitive performance in the models per ROI per metric for the three white matter metrics.

These findings were replicated in both male and female participants.

These results provide evidence that different metrics within grey and white matter better predict cognitive performance.

### 3. Within a metric, not every region gives the same information to the prediction of cognitive performance

In addition to different metrics showing distinct predictive strengths as observed above, it is also highly plausible (e.g. Basten et al., 2015) that there is regional specificity. In other words, that different regions of interest might have a unique predictive role, and that this role depends on the metrics being studied. To assess the effects of the different regions within a metric, we compared a model with freely estimated parameters to a model where all the parameters are equality constrained, which captures the hypothesis that each region contributes equally. With three exceptions, the free models showed the best fit to the data (AICdiff > 50, BICdiff > 15 in favour of the models with freely estimated parameters for 15/18 of the models with all the regions and the ones with the regularized regions). These exceptions pertained exclusively to BIC criteria in the models with all the regions, their occurrence can be attributed to the penalization of increased model complexity.

In addition, the regularized models, which favour sparse models with only few predictors, always retained multiple regions even within the same metric. If a single ‘key’ region contains all relevant predictive information, we would not expect to observe this pattern. Across the nine models that underwent regularization, each model estimation showed at least 30% of all regions to have a regularized beta different from zero, demonstrating the regional specificity and complementarity hypothesized above.

Figure 7 shows how often each region appeared in the final, regularized models across 1000 different subsets of the data. Regions or tracts with a high percentage of survival are those that most consistently provide unique information to predict differences in cognitive performance compared to all other different regions and tracts within the metric. For instance, a researcher interested in the parietal cortex will see in Figure 7 that regions within this lobe are more consistently important (compared to regions in other lobes) in a model including only cortical thickness, than in a model that analyses surface area or grey matter volume.

**Figure 7:**
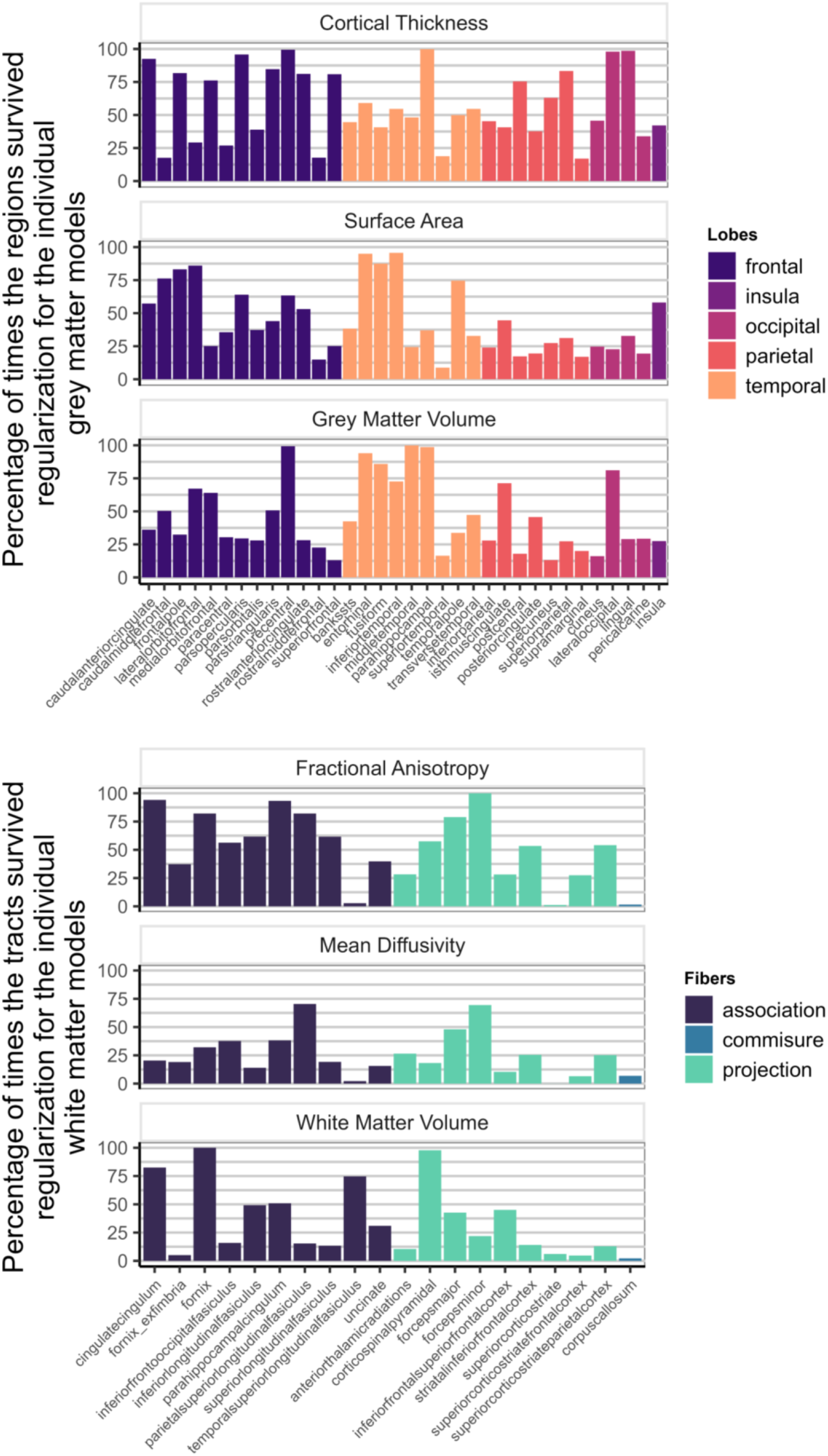
Percentage of times individual region/tract in a metric survived regularization in the individual models for each metric (Model D in Fig 1)

These converging findings demonstrate that to have a better picture of cognitive performance we would need to assess different regions in a metric simultaneously.

### 4. Across metrics, different regions give unique information to the prediction of cognitive performance

Finally, we investigated if the same regions of interest bring unique information to predict cognitive performance across different metrics in each tissue (e.g. if the fractional anisotropy and white matter in a specific tract contain similar information then the regularization will likely regularize one of the metric to 0 for that given tract). A lasso regression was estimated with every region of interest in each grey (or white) matter metrics, the remaining regions were entered in a model as predictors of cognitive performance. This regression allows us to check which regions were nonessential to the prediction of cognitive performance when taking into account all the regions (or tracts) in the three metrics.

We found that despite the strong penalty included in the regularisations, the model still retained regions from each of the grey matter metrics, as well as each of the three white matter metrics, to be significantly predictive of cognitive performance (Figure 8). This suggests that different regions/tracts give unique information to the prediction of cognitive performance.

**Figure 8:**
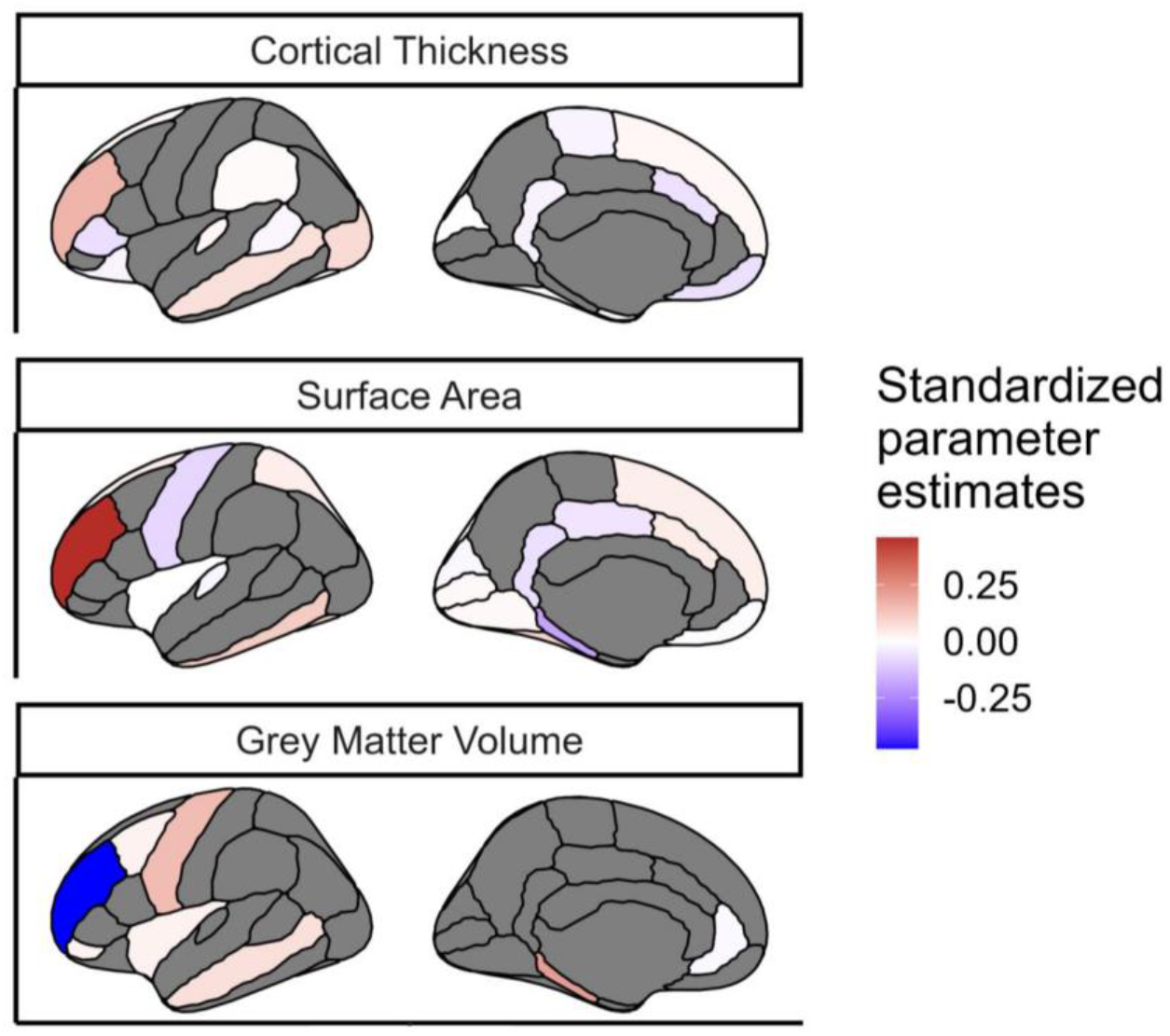
Regions of interest in each grey matter metric that survived the regularization in the model including the three metrics of grey matter. The metrics were averaged bilaterally across the hemispheres. The color indicated the standardized estimated model parameters. The regions in grey did not survived regularization for this model. The regions in white survived regularization but have a standardized estimated model parameter that is close to 0, these regions survived regularization likely because it improved the predictive power of other regions in the model.

Figure 9 illustrates the survival rates of each region/tract across 1000 regularizations in a model incorporating either the three grey matter metrics or the three white matter metrics as predictors of cognitive performance. Comparing the percentage values to those in Figure 7, it becomes evident that a portion of the unique information associated with each region in each metric becomes redundant when performing the regularization analysis with combined grey and white matter metrics.

**Figure 9:**
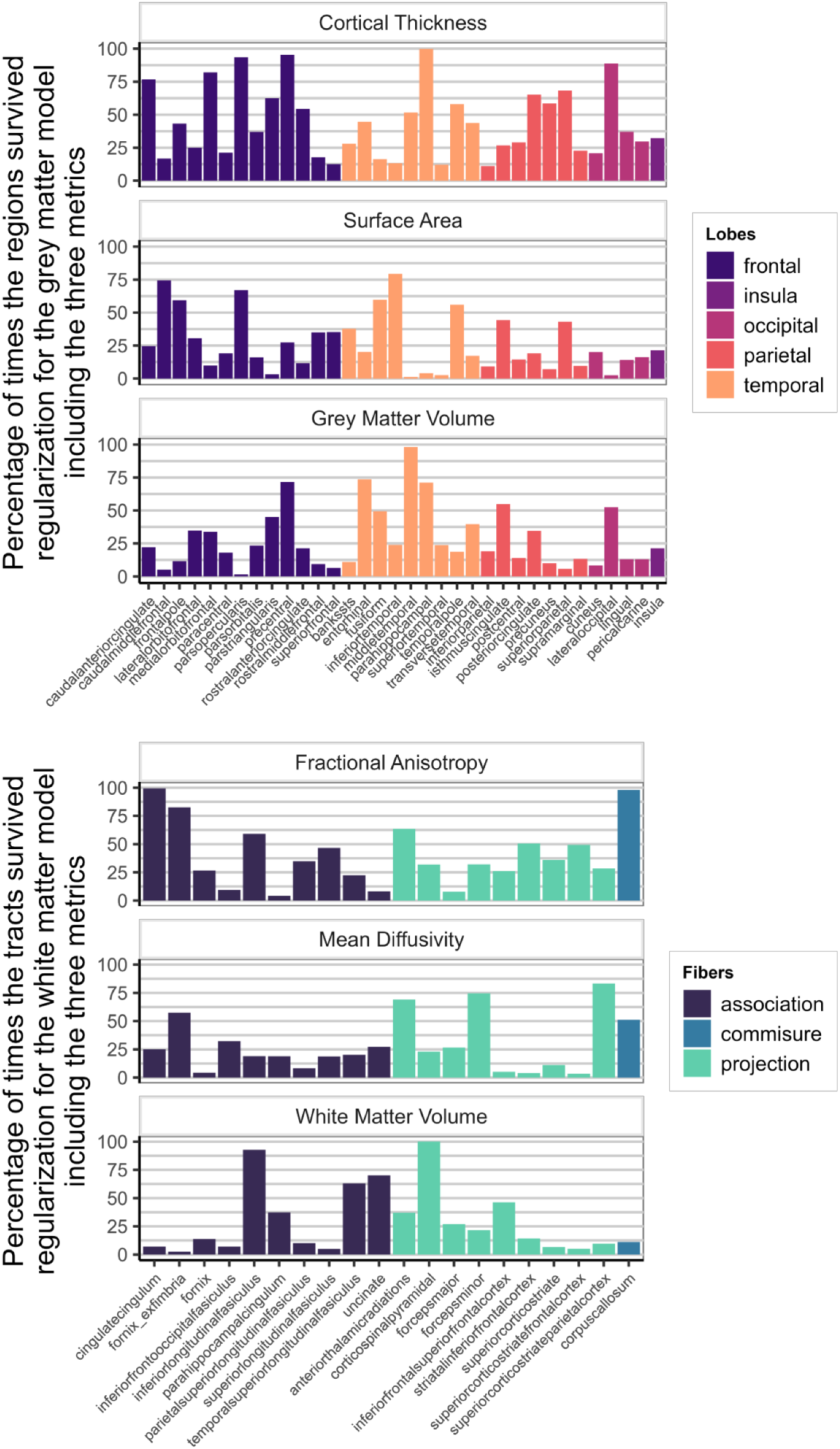
Percentage of times one region in a metric survived regularization in the model with every grey matter metrics included (top figure – Model E in Fig 1) and in the model with every white matter metrics (bottom figure – Model F in Fig 1).

The complete results of all the models studies in this paper are available on OSF [https://osf.io/eygwz].

## Discussion

In a large developmental sample, we examined the role of grey and white matter in supporting cognitive performance. Using regularized structural equation model, we observed that the variance in cognitive performance explained by both grey and white matter (19.0%) is considerably greater than either one in isolation (15.4/12.4%), but not fully additive. This demonstrates that grey and white matter structure bring both unique as well as shared information in predicting individual differences in cognitive abilities in children.

The observed shared information is unsurprising, given the interdependence of these tissues both during development and aging (Giorgio et al., 2010; Hoagey et al., 2019). Previous work suggests close associations between grey and white matter during development (Tamnes, Østby, Fjell, et al., 2010; Kochunov et al., 2011) and some similarity in the ability of grey and white matter measures to predict cognition (Hoagey et al., 2019). Zhao et al. (2021) proposed that differences in sample ages may explain differences in regional and metric specific findings (Zhao et al., 2021) in line with Østby et al. (2011) and Fuhrmann et al. (2020) who observed age-dependent differences in the nature and strength of the relationship between grey matter, white matter and cognitive performance (Østby et al., 2011; Fuhrmann et al., 2020).

Regarding the secondary goals, we found evidence that different metrics within grey and white matter have distinct predictive power in this sample. For grey matter, surface area and volume show the best predictive power compared to cortical thickness, a surprising finding given the prevalence of studies which (only) use cortical thickness as a structural predictor of cognitive performance. However, this result is consistent with several studies reporting that surface area is more related to cognitive performance than cortical thickness (Borgeest et al., 2021; Fürtjes et al., 2023; Pulli et al., 2023). Moreover, recent evidence within the same sample suggests that cortical thickness has lower reliability than surface area or volume, which may differentially attenuate the effects a given study observes (Parsons et al., 2023).

For white matter, we found that in this sample white matter volume is the strongest predictor compared to more ‘advanced’ diffusion-based metrics such as fractional anisotropy and mean diffusivity. This is also unexpected since tract integrity measures are seen as more closely associated with the underlying physiology of tracts and their lesions (Fjell et al., 2008; Xing et al., 2021).

The regional differentiation found at the level of the metrics and the tissues in our models illustrates the importance of selecting regions of interest. Some regions emerge as robust predictors of variance in cognitive performance, while others contribute minimally or lack unique informational value. However, it is apparent that the predictive efficacy of individual regions/tracts varies across the metrics within the models. For instance, cortical thickness of the superior frontal region consistently predicts cognitive performance (Figure 7), but surface area or volume of the same region is considerably less predictive. In general, the inclusion of additional metrics within a model is associated with lower frequency of a region/tract’s survival through regularization, in line with partially overlapping predictive information across different metrics (e.g. Figure 4). In contrast, and somewhat surprisingly, the inclusion of TIV as a covariate had negligible consequences for both the parameter estimates and the overall explanatory power of the model. Focusing on gender as a potential factor by fitting the same model separately yielded similar overall conclusions, apart from an overall higher predictive performance observed in female compared to male. This phenomenon could potentially be explained by variations in motion artefacts (Afacan et al., 2016) or an accelerated cortical maturation process in girls during development (Grabowska, 2017; Fuhrmann et al., 2020, fig 4c).

Our study has several strengths. The uniquely large childhood sample of ABCD allows us to both explore the optimal model in an exploratory sample of only 15% of the data, and validate it in a test set (Srivastava, 2018). As such the sample size enables us to optimize exploration and confirmation, as well as increase parameter precision and power in line with recent recommendations (Marek et al., 2022). Moreover, by using a previously validated measurement model for the latent cognitive variable we further decrease measurement error and increase power and precision (Sauce et al., 2022).

However, the main strength of our study is what is commonly absent in the empirical literature: a multimodal analysis including several measures of grey and white matter and the comparison of their explanatory performance. Although previous studies have simultaneously integrated multimodal imaging data (Richard et al., 2020; Rasero et al., 2021), those more commonly examine the individual covariances between multimodal components or factors rather than examine whether distinct metrics provide unique complementary predictive information. The challenge in models with many predictors is that additional predictors will always increase the strength of the joint prediction, at least within sample. In our modeling approach, we rely on three approaches to guard against needless complexity to differentiate the three models: (i) regularization penalizes the predictive estimates downwards to ensure a parsimonious (and often sparse) model; (ii) model comparison criteria which penalize unnecessary complexity, favoring a model with fewer predictors that does ‘almost’ as well as the more complex model (Jacobucci et al., 2019); and (iii) report effect sizes as adjusted r-squared to account for the number of parameters needed to achieve an overall amount of variance explained.

Despite these strengths, our study also has several limitations. First and foremost, the present analysis focuses on cross-sectional differences, in a specific age range (9-11 years old). This is a unique developmental period for the brain during which grey matter starts to decrease while white matter continues to increase (Bethlehem et al., 2022), so we expect the precise pattern of brain-behaviour relations to be contingent on this specific developmental period. Future studies will extend our results to a longitudinal approach to tease apart leading and lagging effects and provide additional support in line with causal hypotheses (see King et al., 2018; Vijayakumar et al., 2018). Moreover, although ABCD undertook considerable efforts to recruit a representative sample of the USA, our findings cannot be assumed to generalize to more diverse populations within and especially beyond the US (LeWinn et al., 2017; Garavan et al., 2018).

In terms of methodology, the ABCD sample has been collected across different sites and scanners, which may include site or scanner variance we did not incorporate (although Parsons et al., 2023 suggests these site effects in ABCD are considerably smaller than other sources of (un)reliability). Moreover, our findings will be affected by certain pragmatic methodological choices. For example, bilaterally averaging the regions of interest across the hemispheres allowed the possibility to do both regularization and maximum likelihood with a large number of regions. However, it precludes us observing any hemispheric specificity on the prediction of cognitive performance. Finally, the definition of grey and white matter as two different tissues measured by different metrics is inherently an oversimplification. MRI derived measures are proxies of the underlying brain structure and are not able (yet) to isolate specific biophysiological components of grey or white matter. For instance, (apparent) cortical thickness is known to be sensitive to myelinisation of adjacent white matter (Natu et al., 2019), suggesting a partial overlap in statistical contributions of grey and white matter may be as much methodological artefact as biological reality. Finally, segmenting the brain into defined regions reduced structural complexity, correspondingly diminishing explained cognitive performance variance (Fürtjes et al., 2023).

To develop a full picture of the complementary roles of grey and white matter to the prediction of cognitive performance, future studies will need to examine large, multimodal, longitudinal and crucially diverse samples across the lifespan. In addition, numerous new measures are needed to estimate cellular mechanisms more accurately. For instance, sulcal depth and curvature analysis provides detailed cortical structure mapping, while myelin water fraction or magnetisation transfer ratio offer more direct approaches to study axons microstructure and myelinisation (Timmers et al., 2016; Lipp et al., 2019). Future work will benefit from metrics that demonstrate a better representation of underlying cellular mechanisms (e.g. Goriounova & Mansvelder, 2019). Beyond the metrics, the emergence of new scanners with higher resolution like the 11.7 Tesla or the CONNECTOM scanners will help unveil the structure within the six layers of the cortex and regional cortico-cortical connectivity (Nowogrodzki, 2018; Huang et al., 2021; Raven et al., 2023).

The present study was designed to evaluate the extent of the overlap between grey and white matter metrics in the prediction of cognitive performance. Our findings suggest that studies focusing solely on one tissue or one metric when linking brain and cognition are likely missing out on complementary explanatory power. Studies limited by pragmatic concerns should carefully consider which metric to focus on, informed by the phenotype of interest and the population being studied, and make explicit that the findings are likely contingent on the metrics used in a study. Future work should incorporate a more holistic view of brain structure across modalities, metrics and measures to better elucidate the relationships between brain and cognitive performance.

## Acknowledgements

RAK and LM were supported by a Hypatia Fellowship at the RadboudUMC. We would like to thank Dr Sam Parsons and Dr Nick Judd for their valuable inputs on earlier drafts of the manuscript.

Data used in the preparation of this article were obtained from the Adolescent Brain Cognitive DevelopmentSM (ABCD) Study (https://abcdstudy.org), held in the NIMH Data Archive (NDA). This is a multisite, longitudinal study designed to recruit more than 10,000 children age 9-10 and follow them over 10 years into early adulthood. The ABCD Study® is supported by the National Institutes of Health and additional federal partners under award numbers U01DA041048, U01DA050989, U01DA051016, U01DA041022, U01DA051018, U01DA051037, U01DA050987, U01DA041174, U01DA041106, U01DA041117, U01DA041028, U01DA041134, U01DA050988, U01DA051039, U01DA041156, U01DA041025, U01DA041120, U01DA051038, U01DA041148, U01DA041093, U01DA041089, U24DA041123, U24DA041147. A full list of supporters is available at https://abcdstudy.org/federal-partners.html. A listing of participating sites and a complete listing of the study investigators can be found at https://abcdstudy.org/consortium_members/. ABCD consortium investigators designed and implemented the study and/or provided data but did not necessarily participate in the analysis or writing of this report. This manuscript reflects the views of the authors and may not reflect the opinions or views of the NIH or ABCD consortium investigators.

## Footnotes

1 Pubmed search: ((“cortical thickness”[tiab:∼0])) AND ((“cognit*”))

