## Supplementary Material for "Grey and white matter metrics demonstrate distinct and complementary prediction of differences in cognitive performance in children: Findings from ABCD (N= 11 876)"

### Contents

I. Summary of the variables
II. Correlations between the 5 cognitive tasks
III. Confirmatory Factor Analysis
IV. Individual models estimating cognitive factor from one region in one metric
   i. Cortical Thickness
   ii. Surface Area
   iii. Grey Matter Volume
   iv. Fractional Anisotropy
   v. Mean Diffusivity
   vi. White Matter Volume
V. Models per metric estimating cognitive factor from several regions in one metric
   i. Cortical Thickness (all regions and regularized regions)
   ii. Surface Area (all regions and regularized regions)
   iii. Grey Matter Volume (all regions and regularized regions)
   iv. Fractional Anisotropy (all regions and regularized regions)
   v. Mean Diffusivity (all regions and regularized regions)
   vi. White Matter Volume (all regions and regularized regions)
VI. Models per tissue estimating cognitive factor from several regions in three metrics
   i. Grey Matter Metrics (all regions and regularized regions)
   ii. White Matter Metrics (all regions and regularized regions)
VII. Models with grey and white matter metrics estimating cognitive factor from several regions in the six metrics
VIII. Comparison between models with grey & white matter metrics and with grey & white matter metrics & TIV

### Summary of the variables

*Supplementary Table 1. Characteristics of the variables used in the models. All the measures of grey and white matter share the same frequency and number of missing data per tissue.*

| Variable | Stats / Values | Freqs (% of Valid) | Graph | Missing |
|---|---|---|---|---|
| Subjectkey [character] | | 11 876 distinct values | | 0 |
| interview_age [numeric] | Mean (sd) : 119 (7.5)<br>min ≤ med ≤ max:<br>107 ≤ 119 ≤ 133<br>IQR (CV) : 14 (0.1) | 27 distinct values | 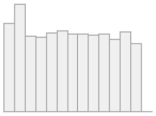 | 0<br>(0.0%) |
| sex [factor] | 1. F<br>2. M | 5680 ( 47.8% )<br>6196 ( 52.2% ) | 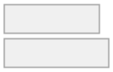 | 0<br>(0.0%) |
| Picture_Vocabulary [numeric] | Mean (sd) : 84.5 (8.1)<br>min ≤ med ≤ max:<br>29 ≤ 84 ≤ 119<br>IQR (CV) : 11 (0.1) | 70 distinct values | 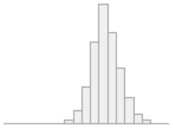 | 147<br>(1.2%) |
| Flanker [numeric] | Mean (sd) : 94 (9.1)<br>min ≤ med ≤ max:<br>51 ≤ 95 ≤ 116<br>IQR (CV) : 11 (0.1) | 68 distinct values | 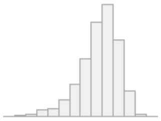 | 153<br>(1.3%) |
| Oral_Reading_Recognition [numeric] | Mean (sd) : 90.9 (6.9)<br>min ≤ med ≤ max:<br>59 ≤ 91 ≤ 119<br>IQR (CV) : 8 (0.1) | 63 distinct values | 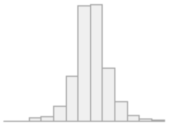 | 161<br>(1.4%) |
| Rey_Auditory_Verbal_Learning [numeric] | Mean (sd) : 44.2 (10)<br>min ≤ med ≤ max:<br>0 ≤ 45 ≤ 72<br>IQR (CV) : 13 (0.2) | 69 distinct values | 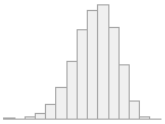 | 292<br>(2.5%) |
| Little_Man [numeric] | Mean (sd) : 58.9 (17)<br>min ≤ med ≤ max:<br>0 ≤ 56.2 ≤ 100<br>IQR (CV) : 25 (0.3) | 32 distinct values | 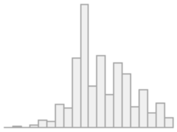 | 337<br>(2.8%) |
| bankssts_ct [numeric] | Mean (sd) : 0.1 (0.8)<br>min ≤ med ≤ max:<br>-6.8 ≤ 0.1 ≤ 4.3<br>IQR (CV) : 1.1 (7.9) | 11521 distinct values | 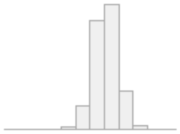 | 116<br>(1.0%) |
| forecepsmajor_fa [numeric] | Mean (sd) : -0.1 (1)<br>min ≤ med ≤ max:<br>-9.3 ≤ 0 ≤ 9.4<br>IQR (CV) : 1.2 (-8.2) | 11107 distinct values | 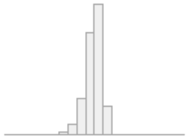 | 739<br>(6.2%) |

### Correlations between the 5 cognitive tasks

*Supplementary Figure 1. Correlation plot of the five measures of cognition: Picture Vocabulary, Flanker, Oral Reading Recognition, Rey Auditory Verbal Learning and Little Man in the whole sample.*

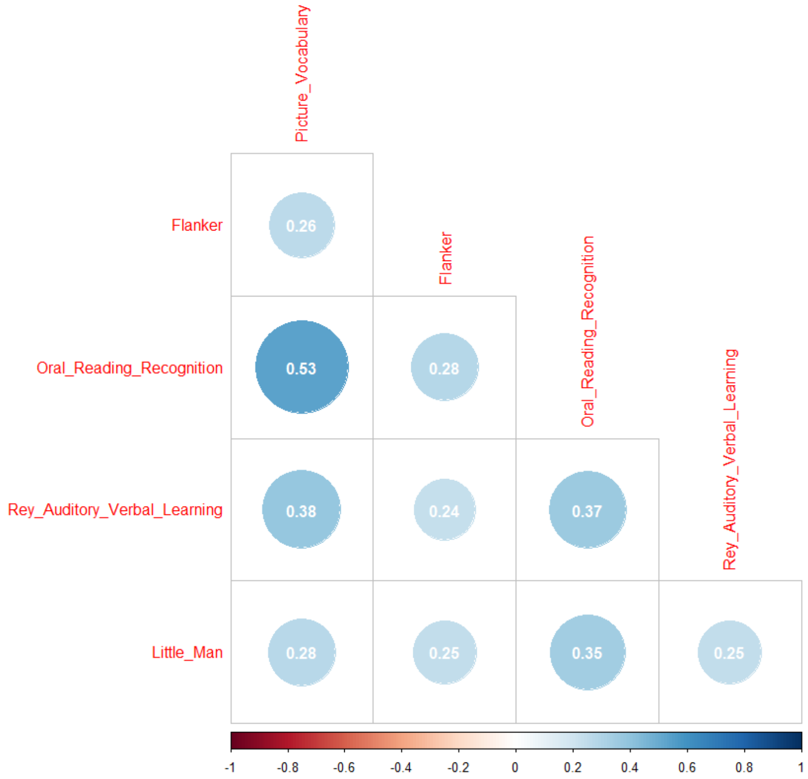


### Confirmatory Factor Analysis

*Supplementary Table 2. Latent variable loadings for the confirmatory factor analysis estimating the cognitive factor*

| Path | Estimate | SE | *p* | Standardized Estimate |
|---|---|---|---|---|
| Cognitive_factor =~ Picture_Vocabulary | 1.000 | 0.000 | NA | 0.699 |
| Cognitive_factor =~ Flanker | 0.582 | 0.020 | *p<0.001* | 0.406 |
| Cognitive_factor =~ Oral_Reading_Recognition | 1.056 | 0.022 | *p<0.001* | 0.738 |
| Cognitive_factor =~ Rey_Auditory_Verbal_Learning | 0.753 | 0.018 | *p<0.001* | 0.527 |
| Cognitive_factor =~ Little_Man | 0.651 | 0.019 | *p<0.001* | 0.455 |

### Individual models estimating cognitive factor from one region in one metric

Note: Every regions is bilaterally averaged. The results reported here are based on the validation sample (85%).

#### Cortical Thickness

*Supplementary Table 3. Regression estimates for the model estimating how cortical thickness of one region predict the cognitive factor*

| Path | | | Estimate | SE | *p* | Standardized Estimate |
|---|---|---|---|---|---|---|
| Cognitive_factor | ~ | bankssts_ct | 0.052 | 0.010 | *p<0.001* | 0.062 |
| Cognitive_factor | ~ | caudalanteriorcingulate_ct | -0.049 | 0.010 | *p<0.001* | -0.059 |
| Cognitive_factor | ~ | caudalmiddlefrontal_ct | 0.030 | 0.009 | *0.001* | 0.038 |
| Cognitive_factor | ~ | cuneus_ct | 0.088 | 0.009 | *p<0.001* | 0.114 |
| Cognitive_factor | ~ | entorhinal_ct | 0.049 | 0.009 | *p<0.001* | 0.061 |
| Cognitive_factor | ~ | fusiform_ct | 0.077 | 0.009 | *p<0.001* | 0.100 |
| Cognitive_factor | ~ | inferiorparietal_ct | 0.032 | 0.009 | *p<0.001* | 0.042 |
| Cognitive_factor | ~ | inferiortemporal_ct | 0.065 | 0.009 | *p<0.001* | 0.085 |
| Cognitive_factor | ~ | isthmuscingulate_ct | 0.003 | 0.010 | 0.778 | 0.003 |
| Cognitive_factor | ~ | lateraloccipital_ct | 0.103 | 0.009 | *p<0.001* | 0.139 |
| Cognitive_factor | ~ | lateralorbitofrontal_ct | 0.020 | 0.009 | *0.027* | 0.026 |
| Cognitive_factor | ~ | lingual_ct | 0.109 | 0.009 | *p<0.001* | 0.145 |
| Cognitive_factor | ~ | medialorbitofrontal_ct | -0.042 | 0.009 | *p<0.001* | -0.053 |
| Cognitive_factor | ~ | middletemporal_ct | 0.058 | 0.009 | *p<0.001* | 0.076 |
| Cognitive_factor | ~ | parahippocampal_ct | 0.098 | 0.009 | *p<0.001* | 0.127 |
| Cognitive_factor | ~ | paracentral_ct | 0.055 | 0.009 | *p<0.001* | 0.072 |
| Cognitive_factor | ~ | parsopercularis_ct | -0.026 | 0.009 | *0.006* | -0.032 |
| Cognitive_factor | ~ | parsorbitalis_ct | -0.012 | 0.010 | 0.238 | -0.014 |
| Cognitive_factor | ~ | parstriangularis_ct | -0.021 | 0.009 | *0.029* | -0.026 |
| Cognitive_factor | ~ | pericalcarine_ct | 0.086 | 0.009 | *p<0.001* | 0.113 |
| Cognitive_factor | ~ | postcentral_ct | 0.066 | 0.009 | *p<0.001* | 0.087 |
| Cognitive_factor | ~ | posteriorcingulate_ct | -0.011 | 0.010 | 0.268 | -0.014 |
| Cognitive_factor | ~ | precentral_ct | 0.081 | 0.009 | *p<0.001* | 0.108 |
| Cognitive_factor | ~ | precuneus_ct | 0.027 | 0.009 | *0.002* | 0.036 |
| Cognitive_factor | ~ | rostralanteriorcingulate_ct | -0.044 | 0.010 | *p<0.001* | -0.052 |
| Cognitive_factor | ~ | rostralmiddlefrontal_ct | 0.004 | 0.009 | 0.661 | 0.005 |
| Cognitive_factor | ~ | superiorfrontal_ct | -0.011 | 0.009 | 0.220 | -0.014 |
| Cognitive_factor | ~ | superiorparietal_ct | 0.032 | 0.009 | *p<0.001* | 0.044 |
| Cognitive_factor | ~ | superiortemporal_ct | 0.044 | 0.009 | *p<0.001* | 0.059 |
| Cognitive_factor | ~ | supramarginal_ct | 0.045 | 0.009 | *p<0.001* | 0.059 |
| Cognitive_factor | ~ | frontalpole_ct | -0.051 | 0.010 | *p<0.001* | -0.061 |
| Cognitive_factor | ~ | temporalpole_ct | 0.057 | 0.009 | *p<0.001* | 0.071 |
| Cognitive_factor | ~ | transversetemporal_ct | 0.053 | 0.009 | *p<0.001* | 0.066 |
| Cognitive_factor | ~ | insula_ct | 0.026 | 0.009 | *0.006* | 0.032 |

*Supplementary Figure 2. Standardized parameter estimates of how the cortical thickness of each region of interest predict the cognitive factor*

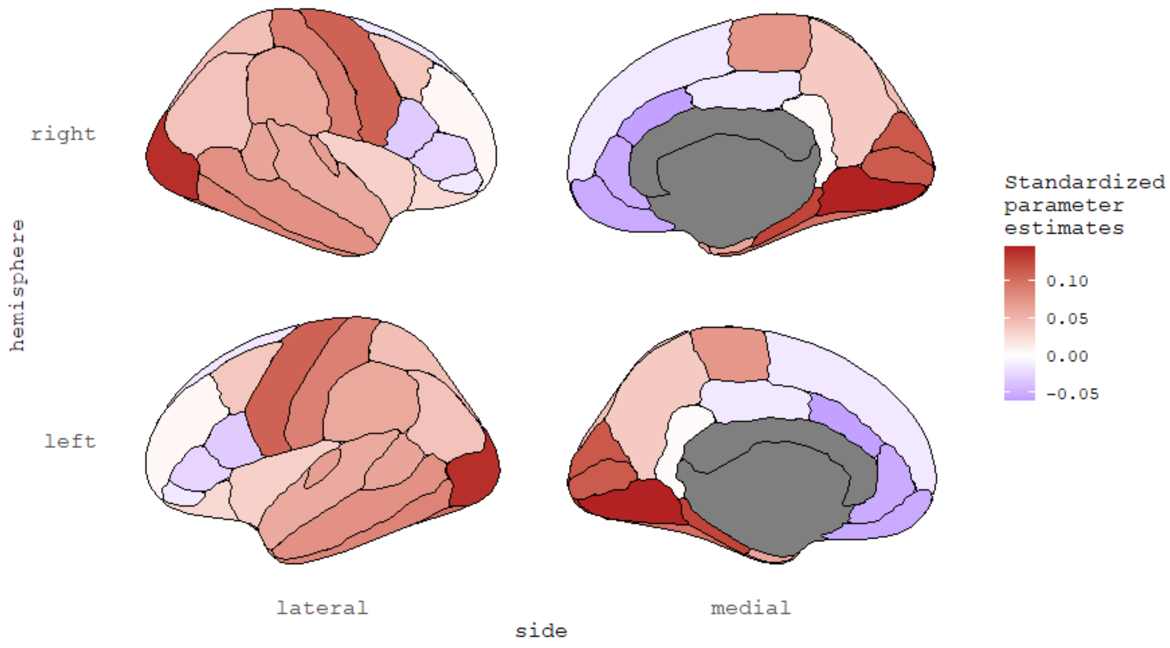


### Surface Area

*Supplementary Table 4. Regression estimates for the model estimating how surface area of one region predict the cognitive factor*

| Path | | | Estimate | SE | *p* | Standardized Estimate |
|---|---|---|---|---|---|---|
| Cognitive_factor | ~ | bankssts_sa | 0.123 | 0.010 | *p<0.001* | 0.152 |
| Cognitive_factor | ~ | caudalanteriorcingulate_sa | 0.172 | 0.011 | *p<0.001* | 0.196 |
| Cognitive_factor | ~ | caudalmiddlefrontal_sa | 0.170 | 0.010 | *p<0.001* | 0.218 |
| Cognitive_factor | ~ | cuneus_sa | 0.099 | 0.009 | *p<0.001* | 0.133 |
| Cognitive_factor | ~ | entorhinal_sa | 0.134 | 0.009 | *p<0.001* | 0.170 |
| Cognitive_factor | ~ | fusiform_sa | 0.193 | 0.009 | *p<0.001* | 0.255 |
| Cognitive_factor | ~ | inferiorparietal_sa | 0.141 | 0.009 | *p<0.001* | 0.185 |
| Cognitive_factor | ~ | inferiortemporal_sa | 0.201 | 0.009 | *p<0.001* | 0.265 |
| Cognitive_factor | ~ | isthmuscingulate_sa | 0.118 | 0.011 | *p<0.001* | 0.153 |
| Cognitive_factor | ~ | lateraloccipital_sa | 0.147 | 0.009 | *p<0.001* | 0.196 |
| Cognitive_factor | ~ | lateralorbitofrontal_sa | 0.181 | 0.009 | *p<0.001* | 0.244 |
| Cognitive_factor | ~ | lingual_sa | 0.107 | 0.009 | *p<0.001* | 0.145 |
| Cognitive_factor | ~ | medialorbitofrontal_sa | 0.176 | 0.009 | *p<0.001* | 0.230 |
| Cognitive_factor | ~ | middletemporal_sa | 0.174 | 0.009 | *p<0.001* | 0.236 |
| Cognitive_factor | ~ | parahippocampal_sa | 0.123 | 0.009 | *p<0.001* | 0.157 |
| Cognitive_factor | ~ | paracentral_sa | 0.127 | 0.009 | *p<0.001* | 0.162 |

| | | | | | | |
|---|---|---|---|---|---|---|
| Cognitive_factor | ~ | parsopercularis_sa | 0.154 | 0.010 | *p<0.001* | 0.193 |
| Cognitive_factor | ~ | parsorbitalis_sa | 0.168 | 0.010 | *p<0.001* | 0.219 |
| Cognitive_factor | ~ | parstriangularis_sa | 0.116 | 0.009 | *p<0.001* | 0.147 |
| Cognitive_factor | ~ | pericalcarine_sa | 0.067 | 0.009 | *p<0.001* | 0.092 |
| Cognitive_factor | ~ | postcentral_sa | 0.150 | 0.009 | *p<0.001* | 0.199 |
| Cognitive_factor | ~ | posteriorcingulate_sa | 0.157 | 0.011 | *p<0.001* | 0.195 |
| Cognitive_factor | ~ | precentral_sa | 0.173 | 0.009 | *p<0.001* | 0.230 |
| Cognitive_factor | ~ | precuneus_sa | 0.159 | 0.009 | *p<0.001* | 0.216 |
| Cognitive_factor | ~ | rostralanteriorcingulate_sa | 0.179 | 0.010 | *p<0.001* | 0.224 |
| Cognitive_factor | ~ | rostralmiddlefrontal_sa | 0.161 | 0.009 | *p<0.001* | 0.215 |
| Cognitive_factor | ~ | superiorfrontal_sa | 0.177 | 0.009 | *p<0.001* | 0.239 |
| Cognitive_factor | ~ | superiorparietal_sa | 0.142 | 0.009 | *p<0.001* | 0.190 |
| Cognitive_factor | ~ | superiortemporal_sa | 0.154 | 0.009 | *p<0.001* | 0.207 |
| Cognitive_factor | ~ | supramarginal_sa | 0.141 | 0.009 | *p<0.001* | 0.181 |
| Cognitive_factor | ~ | frontalpole_sa | 0.186 | 0.010 | *p<0.001* | 0.233 |
| Cognitive_factor | ~ | temporalpole_sa | 0.115 | 0.009 | *p<0.001* | 0.147 |
| Cognitive_factor | ~ | transversetemporal_sa | 0.111 | 0.010 | *p<0.001* | 0.142 |
| Cognitive_factor | ~ | insula_sa | 0.158 | 0.009 | *p<0.001* | 0.211 |

*Supplementary Figure 3. Standardized parameter estimates of how the surface area of each region of interest predict the cognitive factor*

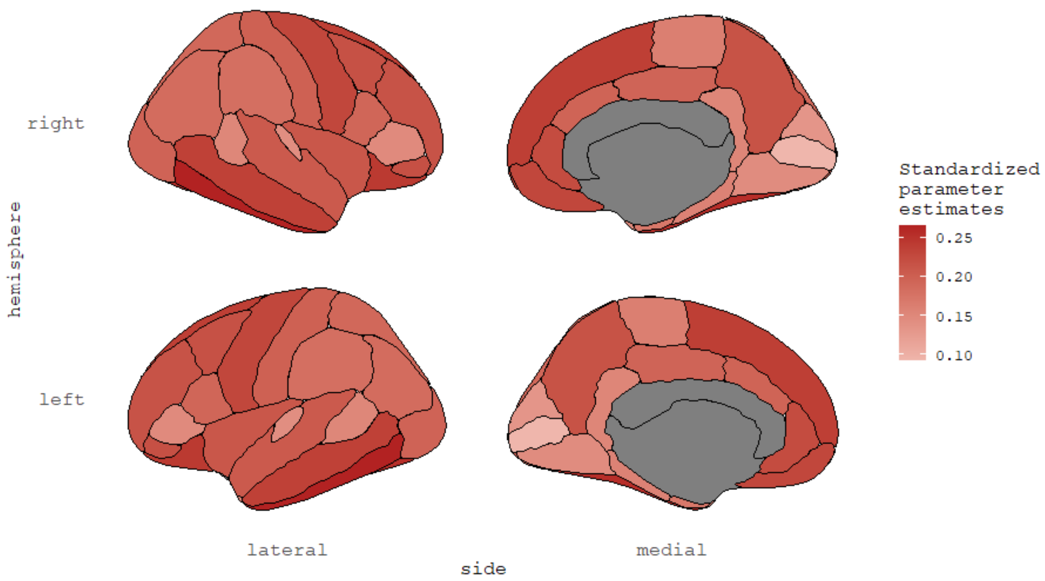

### Grey Matter Volume

*Supplementary Table 5. Regression estimates for the model estimating how grey matter volume of one region predict the cognitive factor*

| Path | | | Estimate | SE | *p* | Standardized Estimate |
|---|---|---|---|---|---|---|
| Cognitive_factor | ~ | bankssts_gmv | 0.136 | 0.010 | *p<0.001* | 0.165 |
| Cognitive_factor | ~ | caudalanteriorcingulate_gmv | 0.158 | 0.011 | *p<0.001* | 0.171 |
| Cognitive_factor | ~ | caudalmiddlefrontal_gmv | 0.165 | 0.009 | *p<0.001* | 0.209 |
| Cognitive_factor | ~ | cuneus_gmv | 0.118 | 0.009 | *p<0.001* | 0.157 |
| Cognitive_factor | ~ | entorhinal_gmv | 0.152 | 0.010 | *p<0.001* | 0.191 |
| Cognitive_factor | ~ | fusiform_gmv | 0.200 | 0.009 | *p<0.001* | 0.261 |
| Cognitive_factor | ~ | inferiorparietal_gmv | 0.151 | 0.009 | *p<0.001* | 0.196 |
| Cognitive_factor | ~ | inferiortemporal_gmv | 0.203 | 0.009 | *p<0.001* | 0.265 |
| Cognitive_factor | ~ | isthmuscingulate_gmv | 0.117 | 0.010 | *p<0.001* | 0.150 |
| Cognitive_factor | ~ | lateraloccipital_gmv | 0.184 | 0.009 | *p<0.001* | 0.245 |
| Cognitive_factor | ~ | lateralorbitofrontal_gmv | 0.185 | 0.009 | *p<0.001* | 0.251 |
| Cognitive_factor | ~ | lingual_gmv | 0.133 | 0.009 | *p<0.001* | 0.178 |
| Cognitive_factor | ~ | medialorbitofrontal_gmv | 0.138 | 0.009 | *p<0.001* | 0.178 |
| Cognitive_factor | ~ | middletemporal_gmv | 0.205 | 0.009 | *p<0.001* | 0.275 |
| Cognitive_factor | ~ | parahippocampal_gmv | 0.166 | 0.010 | *p<0.001* | 0.208 |
| Cognitive_factor | ~ | paracentral_gmv | 0.142 | 0.009 | *p<0.001* | 0.179 |
| Cognitive_factor | ~ | parsopercularis_gmv | 0.134 | 0.010 | *p<0.001* | 0.165 |
| Cognitive_factor | ~ | parsorbitalis_gmv | 0.159 | 0.010 | *p<0.001* | 0.202 |
| Cognitive_factor | ~ | parstriangularis_gmv | 0.099 | 0.009 | *p<0.001* | 0.123 |
| Cognitive_factor | ~ | pericalcarine_gmv | 0.090 | 0.009 | *p<0.001* | 0.123 |
| Cognitive_factor | ~ | postcentral_gmv | 0.168 | 0.009 | *p<0.001* | 0.221 |
| Cognitive_factor | ~ | posteriorcingulate_gmv | 0.153 | 0.010 | *p<0.001* | 0.189 |
| Cognitive_factor | ~ | precentral_gmv | 0.205 | 0.009 | *p<0.001* | 0.269 |
| Cognitive_factor | ~ | precuneus_gmv | 0.159 | 0.009 | *p<0.001* | 0.216 |
| Cognitive_factor | ~ | rostralanteriorcingulate_gmv | 0.166 | 0.010 | *p<0.001* | 0.200 |
| Cognitive_factor | ~ | rostralmiddlefrontal_gmv | 0.152 | 0.009 | *p<0.001* | 0.201 |
| Cognitive_factor | ~ | superiorfrontal_gmv | 0.174 | 0.009 | *p<0.001* | 0.233 |
| Cognitive_factor | ~ | superiorparietal_gmv | 0.148 | 0.009 | *p<0.001* | 0.197 |
| Cognitive_factor | ~ | superiortemporal_gmv | 0.170 | 0.009 | *p<0.001* | 0.225 |
| Cognitive_factor | ~ | supramarginal_gmv | 0.158 | 0.009 | *p<0.001* | 0.201 |
| Cognitive_factor | ~ | frontalpole_gmv | 0.114 | 0.010 | *p<0.001* | 0.140 |
| Cognitive_factor | ~ | temporalpole_gmv | 0.129 | 0.010 | *p<0.001* | 0.160 |
| Cognitive_factor | ~ | transversetemporal_gmv | 0.142 | 0.010 | *p<0.001* | 0.179 |
| Cognitive_factor | ~ | insula_gmv | 0.156 | 0.009 | *p<0.001* | 0.213 |

*Supplementary Figure 4. Standardized parameter estimates of how the grey matter volume of each region of interest predict the cognitive factor*

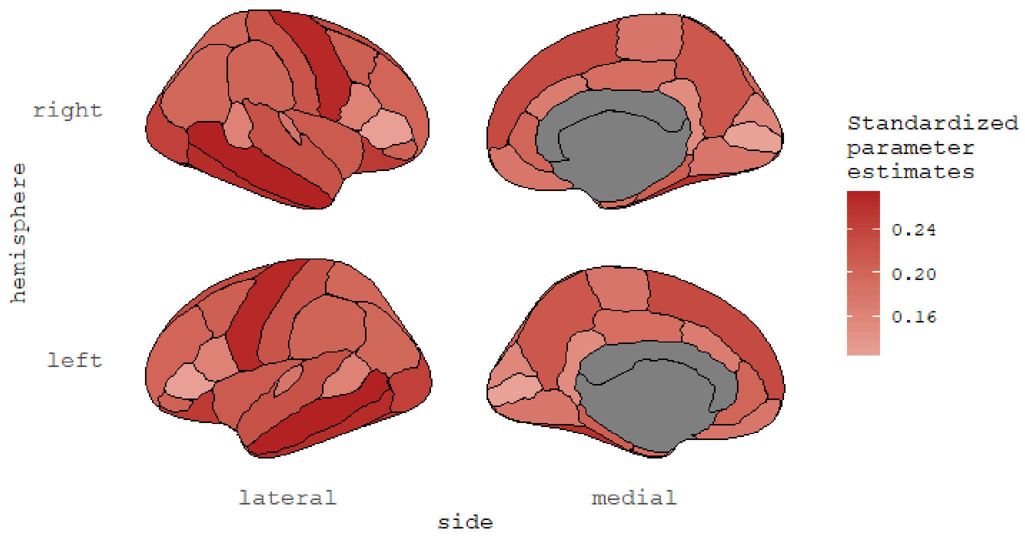


### Fractional Anisotropy

*Supplementary Table 6. Regression estimates for the model estimating how fractional anisotropy of one region predict the cognitive factor*

| Path | | | Estimate | SE | *p* | Standardized Estimate |
|---|---|---|---|---|---|---|
| Cognitive_factor | ~ | fornix_fa | 0.063 | 0.009 | *p<0.001* | 0.085 |
| Cognitive_factor | ~ | cingulatecingulum_fa | 0.001 | 0.009 | 0.889 | 0.002 |
| Cognitive_factor | ~ | parahippocampalcingulum_fa | 0.077 | 0.009 | *p<0.001* | 0.103 |
| Cognitive_factor | ~ | corticospinalpyramidal_fa | 0.068 | 0.009 | *p<0.001* | 0.093 |
| Cognitive_factor | ~ | anteriorthalamicradiations_fa | 0.032 | 0.009 | *p<0.001* | 0.043 |
| Cognitive_factor | ~ | uncinate_fa | 0.033 | 0.009 | *p<0.001* | 0.044 |
| Cognitive_factor | ~ | inferiorlongitudinalfasiculus_fa | 0.085 | 0.010 | *p<0.001* | 0.114 |
| Cognitive_factor | ~ | inferiorfrontooccipitalfasiculus_fa | 0.067 | 0.009 | *p<0.001* | 0.092 |
| Cognitive_factor | ~ | forcepsmajor_fa | 0.064 | 0.009 | *p<0.001* | 0.093 |
| Cognitive_factor | ~ | forcepsminor_fa | -0.007 | 0.008 | 0.400 | -0.010 |
| Cognitive_factor | ~ | corpuscallosum_fa | 0.040 | 0.009 | *p<0.001* | 0.059 |
| Cognitive_factor | ~ | superiorlongitudinalfasiculus_fa | 0.102 | 0.009 | *p<0.001* | 0.141 |
| Cognitive_factor | ~ | temporalsuperiorlongitudinalfasiculus_fa | 0.094 | 0.009 | *p<0.001* | 0.128 |
| Cognitive_factor | ~ | parietalsuperiorlongitudinalfasiculus_fa | 0.104 | 0.009 | *p<0.001* | 0.142 |
| Cognitive_factor | ~ | superiorcorticostriate_fa | 0.054 | 0.009 | *p<0.001* | 0.073 |
| Cognitive_factor | ~ | superiorcorticostriatefrontalcortex_fa | 0.057 | 0.009 | *p<0.001* | 0.078 |
| Cognitive_factor | ~ | superiorcorticostriateparietalcortex_fa | 0.044 | 0.009 | *p<0.001* | 0.059 |

| | | | | | | |
|---|---|---|---|---|---|---|
| Cognitive_factor | ~ | striatalinferiorfrontalcortex_fa | 0.014 | 0.009 | 0.121 | 0.019 |
| Cognitive_factor | ~ | inferiorfrontalsuperiorfrontalcortex_fa | 0.066 | 0.009 | *p<0.001* | 0.090 |
| Cognitive_factor | ~ | fornix_exfimbria_fa | 0.015 | 0.009 | 0.102 | 0.020 |

### Mean Diffusivity

*Supplementary Table 7. Regression estimates for the model estimating how mean diffusivity of one region predict the cognitive factor*

| Path | | | Estimate | SE | *p* | Standardized Estimate |
|---|---|---|---|---|---|---|
| Cognitive_factor | ~ | fornix_md | -0.021 | 0.010 | 0.038 | -0.029 |
| Cognitive_factor | ~ | cingulatecingulum_md | -0.020 | 0.010 | *0.033* | -0.025 |
| Cognitive_factor | ~ | parahippocampalcingulum_md | -0.024 | 0.009 | *0.005* | -0.033 |
| Cognitive_factor | ~ | corticospinalpyramidal_md | -0.014 | 0.008 | 0.068 | -0.019 |
| Cognitive_factor | ~ | anteriorthalamicradiations_md | 0.007 | 0.008 | 0.386 | 0.010 |
| Cognitive_factor | ~ | uncinate_md | -0.006 | 0.007 | 0.404 | -0.008 |
| Cognitive_factor | ~ | inferiorlongitudinalfasiculus_md | -0.021 | 0.009 | *0.015* | -0.028 |
| Cognitive_factor | ~ | inferiorfrontooccipitalfasiculus_md | 0.003 | 0.009 | 0.711 | 0.004 |
| Cognitive_factor | ~ | forcepsmajor_md | -0.027 | 0.010 | *0.007* | -0.039 |
| Cognitive_factor | ~ | forcepsminor_md | 0.021 | 0.008 | *0.012* | 0.031 |
| Cognitive_factor | ~ | corpuscallosum_md | -0.013 | 0.009 | 0.131 | -0.019 |
| Cognitive_factor | ~ | superiorlongitudinalfasiculus_md | -0.040 | 0.009 | *p<0.001* | -0.056 |
| Cognitive_factor | ~ | temporalsuperiorlongitudinalfasiculus_md | -0.032 | 0.009 | *p<0.001* | -0.045 |
| Cognitive_factor | ~ | parietalsuperiorlongitudinalfasiculus_md | -0.044 | 0.009 | *p<0.001* | -0.062 |
| Cognitive_factor | ~ | superiorcorticostriate_md | -0.024 | 0.008 | *0.004* | -0.033 |
| Cognitive_factor | ~ | superiorcorticostriatefrontalcortex_md | -0.015 | 0.008 | 0.062 | -0.022 |
| Cognitive_factor | ~ | superiorcorticostriateparietalcortex_md | -0.030 | 0.008 | *p<0.001* | -0.042 |
| Cognitive_factor | ~ | striatalinferiorfrontalcortex_md | 0.003 | 0.009 | 0.760 | 0.003 |
| Cognitive_factor | ~ | inferiorfrontalsuperiorfrontalcortex_md | -0.017 | 0.009 | *0.044* | -0.024 |
| Cognitive_factor | ~ | fornix_exfimbria_md | -0.007 | 0.010 | 0.481 | -0.010 |

### White Matter Volume

*Supplementary Table 8. Regression estimates for the model estimating how white matter volume of one region predict the cognitive factor*

| Path | | | Estimate | SE | *p* | Standardized Estimate |
|---|---|---|---|---|---|---|
| Cognitive_factor | ~ | fornix_wmv | 0.219 | 0.010 | *p<0.001* | 0.303 |
| Cognitive_factor | ~ | cingulatecingulum_wmv | 0.124 | 0.009 | *p<0.001* | 0.169 |
| Cognitive_factor | ~ | parahippocampalcingulum_wmv | 0.155 | 0.010 | *p<0.001* | 0.209 |
| Cognitive_factor | ~ | corticospinalpyramidal_wmv | 0.198 | 0.010 | *p<0.001* | 0.274 |
| Cognitive_factor | ~ | anteriorthalamicradiations_wmv | 0.153 | 0.009 | *p<0.001* | 0.215 |
| Cognitive_factor | ~ | uncinate_wmv | 0.136 | 0.009 | *p<0.001* | 0.187 |

| | | | | | | |
|---|---|---|---|---|---|---|
| Cognitive_factor | ~ | inferiorlongitudinalfasiculus_wmv | 0.166 | 0.010 | *p<0.001* | 0.226 |
| Cognitive_factor | ~ | inferiorfrontooccipitalfasiculus_wmv | 0.174 | 0.010 | *p<0.001* | 0.240 |
| Cognitive_factor | ~ | forcepsmajor_wmv | 0.161 | 0.009 | *p<0.001* | 0.230 |
| Cognitive_factor | ~ | forcepsminor_wmv | 0.146 | 0.009 | *p<0.001* | 0.209 |
| Cognitive_factor | ~ | corpuscallosum_wmv | 0.179 | 0.009 | *p<0.001* | 0.256 |
| Cognitive_factor | ~ | superiorlongitudinalfasiculus_wmv | 0.191 | 0.010 | *p<0.001* | 0.261 |
| Cognitive_factor | ~ | temporalsuperiorlongitudinalfasiculus_wmv | 0.197 | 0.010 | *p<0.001* | 0.266 |
| Cognitive_factor | ~ | parietalsuperiorlongitudinalfasiculus_wmv | 0.183 | 0.010 | *p<0.001* | 0.251 |
| Cognitive_factor | ~ | superiorcorticostriate_wmv | 0.172 | 0.009 | *p<0.001* | 0.237 |
| Cognitive_factor | ~ | superiorcorticostriatefrontalcortex_wmv | 0.169 | 0.009 | *p<0.001* | 0.234 |
| Cognitive_factor | ~ | superiorcorticostriateparietalcortex_wmv | 0.172 | 0.009 | *p<0.001* | 0.233 |
| Cognitive_factor | ~ | striatalinferiorfrontalcortex_wmv | 0.149 | 0.009 | *p<0.001* | 0.203 |
| Cognitive_factor | ~ | inferiorfrontalsuperiorfrontalcortex_wmv | 0.181 | 0.009 | *p<0.001* | 0.247 |
| Cognitive_factor | ~ | fornix_exfimbria_wmv | 0.201 | 0.010 | *p<0.001* | 0.268 |

### Models per metric estimating cognitive factor from several regions in one metric

For every metric, we reported the table for the models with all the regions included as predictors and the models with only the regions that survived the regularization (the one that we used for the analyses).

#### Cortical Thickness (model with all the regions)

*Supplementary Table 9. Regression estimates for the model estimating how cortical thickness of all the regions predict the cognitive factor*

| Path | | | Estimate | SE | *p* | Standardized Estimate |
|---|---|---|---|---|---|---|
| Cognitive_factor | ~ | bankssts_ct | 0.018 | 0.013 | 0.164 | 0.021 |
| Cognitive_factor | ~ | caudalanteriorcingulate_ct | -0.049 | 0.012 | *p<0.001* | -0.058 |
| Cognitive_factor | ~ | caudalmiddlefrontal_ct | 0.025 | 0.016 | 0.124 | 0.032 |
| Cognitive_factor | ~ | cuneus_ct | 0.021 | 0.014 | 0.120 | 0.028 |
| Cognitive_factor | ~ | entorhinal_ct | 0.006 | 0.010 | 0.560 | 0.008 |
| Cognitive_factor | ~ | fusiform_ct | 0.007 | 0.015 | 0.641 | 0.009 |
| Cognitive_factor | ~ | inferiorparietal_ct | -0.053 | 0.018 | *0.003* | -0.069 |
| Cognitive_factor | ~ | inferiortemporal_ct | 0.023 | 0.014 | 0.090 | 0.030 |
| Cognitive_factor | ~ | isthmuscingulate_ct | -0.008 | 0.010 | 0.436 | -0.010 |
| Cognitive_factor | ~ | lateraloccipital_ct | 0.091 | 0.015 | *p<0.001* | 0.123 |
| Cognitive_factor | ~ | lateralorbitofrontal_ct | 0.017 | 0.013 | 0.184 | 0.022 |
| Cognitive_factor | ~ | lingual_ct | 0.062 | 0.013 | *p<0.001* | 0.082 |
| Cognitive_factor | ~ | medialorbitofrontal_ct | -0.036 | 0.012 | *0.004* | -0.044 |
| Cognitive_factor | ~ | middletemporal_ct | 0.030 | 0.015 | 0.046 | 0.039 |
| Cognitive_factor | ~ | parahippocampal_ct | 0.076 | 0.010 | *p<0.001* | 0.098 |
| Cognitive_factor | ~ | paracentral_ct | 0.002 | 0.015 | 0.882 | 0.003 |
| Cognitive_factor | ~ | parsopercularis_ct | -0.087 | 0.014 | *p<0.001* | -0.107 |
| Cognitive_factor | ~ | parsorbitalis_ct | -0.005 | 0.013 | 0.670 | -0.007 |
| Cognitive_factor | ~ | parstriangularis_ct | -0.045 | 0.014 | *0.002* | -0.055 |
| Cognitive_factor | ~ | pericalcarine_ct | -0.007 | 0.012 | 0.562 | -0.009 |
| Cognitive_factor | ~ | postcentral_ct | 0.055 | 0.013 | *p<0.001* | 0.072 |
| Cognitive_factor | ~ | posteriorcingulate_ct | 0.010 | 0.013 | 0.429 | 0.012 |
| Cognitive_factor | ~ | precentral_ct | 0.116 | 0.016 | *p<0.001* | 0.152 |
| Cognitive_factor | ~ | precuneus_ct | -0.024 | 0.015 | 0.112 | -0.032 |
| Cognitive_factor | ~ | rostralanteriorcingulate_ct | -0.035 | 0.012 | *0.004* | -0.042 |
| Cognitive_factor | ~ | rostralmiddlefrontal_ct | 0.036 | 0.017 | 0.033 | 0.047 |
| Cognitive_factor | ~ | superiorfrontal_ct | -0.074 | 0.017 | *p<0.001* | -0.101 |
| Cognitive_factor | ~ | superiorparietal_ct | -0.087 | 0.018 | *p<0.001* | -0.117 |
| Cognitive_factor | ~ | superiortemporal_ct | -0.015 | 0.015 | 0.321 | -0.020 |
| Cognitive_factor | ~ | supramarginal_ct | -0.009 | 0.017 | 0.608 | -0.011 |
| Cognitive_factor | ~ | frontalpole_ct | -0.029 | 0.012 | *0.013* | -0.034 |
| Cognitive_factor | ~ | temporalpole_ct | 0.022 | 0.012 | 0.064 | 0.027 |
| Cognitive_factor | ~ | transversetemporal_ct | 0.020 | 0.012 | 0.085 | 0.025 |
| Cognitive_factor | ~ | insula_ct | 0.014 | 0.012 | 0.219 | 0.018 |

*Supplementary Figure 5. Standardized parameter estimates of how cortical thickness of each region of interest together predict the cognitive factor*

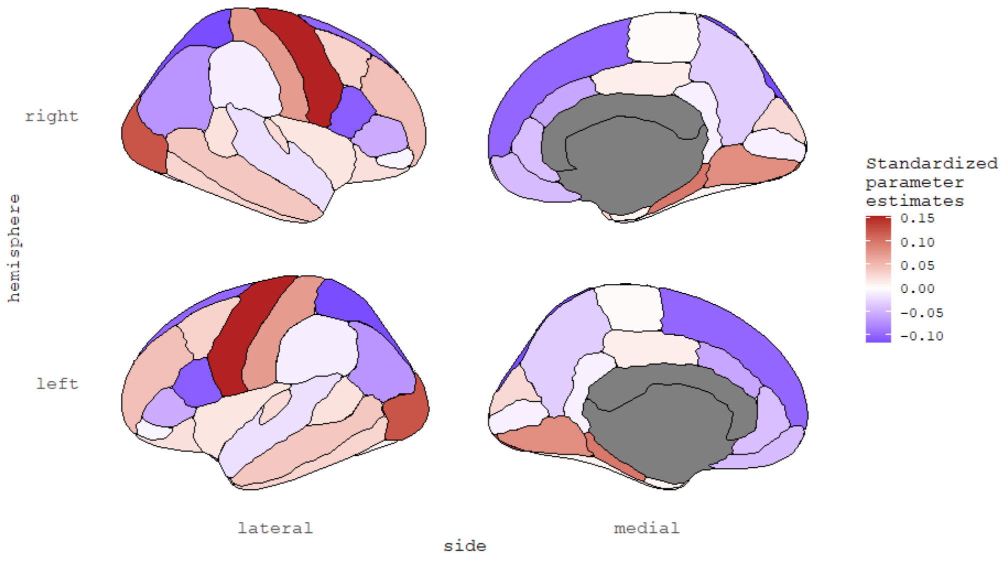


*Supplementary Table 10. Comparison of the model with free parameters and the model with parameters constrained to a same value for the cortical thickness model with all the regions. AIC/BIC are two information criterions. Pr(>Chisq): p-value of the chi-square ratio test.*

| | Degree of freedom | AIC | BIC | Difference in degree of freedom | Pr (>Chisq) |
|---|---|---|---|---|---|
| Free model | 141 | 130758.9 | 131112.2 | NA | NA |
| Constrained model | 174 | 131314.0 | 131429.4 | 33 | *p<0.001* |

### Cortical Thickness (model with the regularized regions)

*Supplementary Table 11. Regression estimates for the model estimating how cortical thickness of the regularized regions predict the cognitive factor*

| Path | | | Estimate | SE | *p* | Standardized Estimate |
|---|---|---|---|---|---|---|
| Cognitive_factor | ~ | bankssts_ct | 0.010 | 0.013 | 0.439 | 0.011 |
| Cognitive_factor | ~ | caudalanteriorcingulate_ct | -0.046 | 0.011 | *p<0.001* | -0.054 |
| Cognitive_factor | ~ | entorhinal_ct | 0.013 | 0.010 | 0.210 | 0.016 |
| Cognitive_factor | ~ | inferiortemporal_ct | 0.018 | 0.013 | 0.164 | 0.024 |
| Cognitive_factor | ~ | isthmuscingulate_ct | -0.007 | 0.010 | 0.488 | -0.009 |
| Cognitive_factor | ~ | lateraloccipital_ct | 0.090 | 0.014 | *p<0.001* | 0.121 |
| Cognitive_factor | ~ | lingual_ct | 0.071 | 0.012 | *p<0.001* | 0.094 |
| Cognitive_factor | ~ | middletemporal_ct | 0.016 | 0.014 | 0.242 | 0.021 |
| Cognitive_factor | ~ | parahippocampal_ct | 0.074 | 0.010 | *p<0.001* | 0.095 |

| | | | | | | |
|---|---|---|---|---|---|---|
| Cognitive_factor | ~ | paracentral_ct | 0.009 | 0.014 | 0.553 | 0.011 |
| Cognitive_factor | ~ | parsopercularis_ct | -0.085 | 0.013 | *p<0.001* | -0.105 |
| Cognitive_factor | ~ | parstriangularis_ct | -0.051 | 0.014 | *p<0.001* | -0.063 |
| Cognitive_factor | ~ | pericalcarine_ct | 0.002 | 0.012 | 0.852 | 0.003 |
| Cognitive_factor | ~ | postcentral_ct | 0.057 | 0.012 | *p<0.001* | 0.075 |
| Cognitive_factor | ~ | precentral_ct | 0.128 | 0.015 | *p<0.001* | 0.167 |
| Cognitive_factor | ~ | precuneus_ct | -0.032 | 0.015 | *0.031* | -0.042 |
| Cognitive_factor | ~ | rostralanteriorcingulate_ct | -0.042 | 0.011 | *p<0.001* | -0.049 |
| Cognitive_factor | ~ | rostralmiddlefrontal_ct | 0.023 | 0.016 | 0.139 | 0.031 |
| Cognitive_factor | ~ | superiorfrontal_ct | -0.074 | 0.016 | *p<0.001* | -0.099 |
| Cognitive_factor | ~ | superiorparietal_ct | -0.107 | 0.016 | *p<0.001* | -0.144 |
| Cognitive_factor | ~ | temporalpole_ct | 0.021 | 0.011 | 0.060 | 0.027 |
| Cognitive_factor | ~ | transversetemporal_ct | 0.016 | 0.011 | 0.146 | 0.020 |

*Supplementary Figure 6. Standardized parameter estimates of how the cortical thickness of each regularized region of interest together predict the cognitive factor*

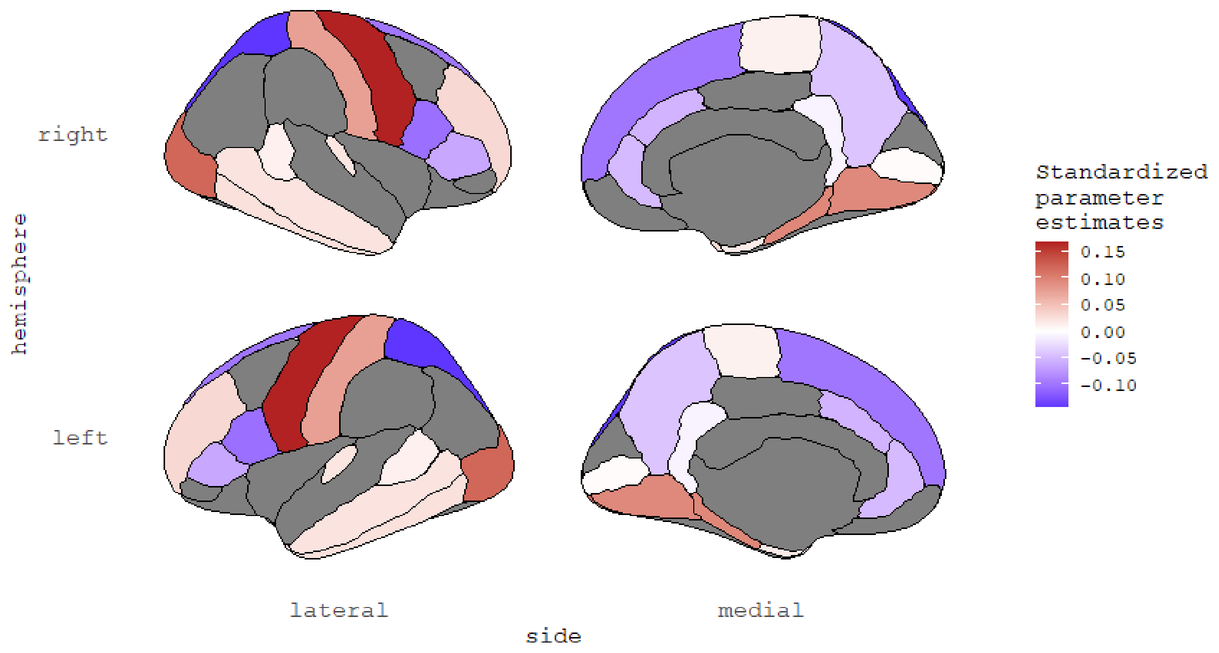


*Supplementary Table 12. Comparison of the model with free parameters and the model with parameters constrained to a same value for the cortical thickness model with the regularized regions. AIC/BIC are two information criterions. Pr(>Chisq): p-value of the chi-square ratio test.*

| | **Degree of freedom** | **AIC** | **BIC** | **Difference in degree of freedom** | **Pr (>Chisq)** |
|---|---|---|---|---|---|
| Free model | 93 | 130773.4 | 131040.1 | NA | NA |
| Constrained model | 114 | 131299.2 | 131414.5 | 21 | *p<0.001* |

### Surface Area (model with all the regions)

*Supplementary Table 13. Regression estimates for the model estimating how surface area of all the regions predict the cognitive factor*

| Path | | | Estimate | SE | *p* | Standardized Estimate |
|---|---|---|---|---|---|---|
| Cognitive_factor | ~ | bankssts_sa | -0.038 | 0.016 | *0.014* | -0.047 |
| Cognitive_factor | ~ | caudalanteriorcingulate_sa | 0.041 | 0.017 | *0.018* | 0.046 |
| Cognitive_factor | ~ | caudalmiddlefrontal_sa | 0.032 | 0.015 | *0.036* | 0.041 |
| Cognitive_factor | ~ | cuneus_sa | -0.025 | 0.018 | 0.151 | -0.033 |
| Cognitive_factor | ~ | entorhinal_sa | 0.071 | 0.012 | *p<0.001* | 0.089 |
| Cognitive_factor | ~ | fusiform_sa | 0.054 | 0.017 | *0.001* | 0.071 |
| Cognitive_factor | ~ | inferiorparietal_sa | -0.028 | 0.015 | 0.070 | -0.035 |
| Cognitive_factor | ~ | inferiortemporal_sa | 0.090 | 0.020 | *p<0.001* | 0.118 |
| Cognitive_factor | ~ | isthmuscingulate_sa | -0.040 | 0.015 | *0.006* | -0.051 |
| Cognitive_factor | ~ | lateraloccipital_sa | 0.010 | 0.015 | 0.514 | 0.013 |
| Cognitive_factor | ~ | lateralorbitofrontal_sa | 0.059 | 0.017 | *0.001* | 0.079 |
| Cognitive_factor | ~ | lingual_sa | 0.030 | 0.017 | 0.077 | 0.039 |
| Cognitive_factor | ~ | medialorbitofrontal_sa | 0.001 | 0.016 | 0.967 | 0.001 |
| Cognitive_factor | ~ | middletemporal_sa | 0.013 | 0.023 | 0.568 | 0.018 |
| Cognitive_factor | ~ | parahippocampal_sa | -0.001 | 0.012 | 0.955 | -0.001 |
| Cognitive_factor | ~ | paracentral_sa | -0.040 | 0.014 | *0.004* | -0.051 |
| Cognitive_factor | ~ | parsopercularis_sa | 0.045 | 0.014 | *0.002* | 0.056 |
| Cognitive_factor | ~ | parsorbitalis_sa | 0.035 | 0.016 | *0.032* | 0.045 |
| Cognitive_factor | ~ | parstriangularis_sa | -0.062 | 0.016 | *p<0.001* | -0.078 |
| Cognitive_factor | ~ | pericalcarine_sa | -0.004 | 0.018 | 0.828 | -0.005 |
| Cognitive_factor | ~ | postcentral_sa | -0.043 | 0.017 | *0.011* | -0.057 |
| Cognitive_factor | ~ | posteriorcingulate_sa | 0.005 | 0.017 | 0.743 | 0.007 |
| Cognitive_factor | ~ | precentral_sa | 0.029 | 0.017 | 0.084 | 0.039 |
| Cognitive_factor | ~ | precuneus_sa | 0.020 | 0.016 | 0.202 | 0.027 |
| Cognitive_factor | ~ | rostralanteriorcingulate_sa | 0.008 | 0.016 | 0.634 | 0.010 |
| Cognitive_factor | ~ | rostralmiddlefrontal_sa | -0.013 | 0.016 | 0.421 | -0.017 |
| Cognitive_factor | ~ | superiorfrontal_sa | 0.024 | 0.018 | 0.195 | 0.032 |
| Cognitive_factor | ~ | superiorparietal_sa | 0.021 | 0.015 | 0.160 | 0.028 |
| Cognitive_factor | ~ | superiortemporal_sa | 0.016 | 0.021 | 0.442 | 0.021 |
| Cognitive_factor | ~ | supramarginal_sa | -0.008 | 0.015 | 0.620 | -0.010 |
| Cognitive_factor | ~ | frontalpole_sa | 0.050 | 0.014 | *p<0.001* | 0.062 |
| Cognitive_factor | ~ | temporalpole_sa | -0.096 | 0.014 | *p<0.001* | -0.121 |
| Cognitive_factor | ~ | transversetemporal_sa | -0.033 | 0.015 | *0.029* | -0.041 |
| Cognitive_factor | ~ | insula_sa | 0.039 | 0.015 | *0.007* | 0.052 |

*Supplementary Figure 7. Standardized parameter estimates of how the surface area of each region of interest together predict the cognitive factor*

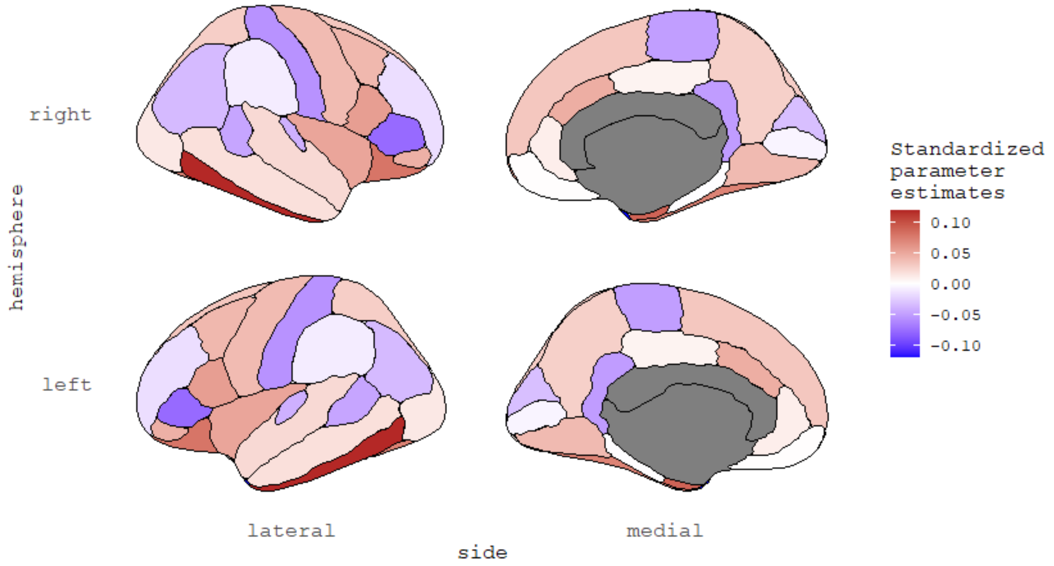


*Supplementary Table 14. Comparison of the model with free parameters and the model with parameters constrained to a same value for the surface area model with all the regions. AIC/BIC are two information criterions. Pr(>Chisq): p-value of the chi-square ratio test.*

| | Degree of freedom | AIC | BIC | Difference in degree of freedom | Pr (>Chisq) |
|---|---|---|---|---|---|
| Free model | 141 | 130573.0 | 130926.3 | NA | NA |
| Constrained model | 174 | 130779.8 | 130895.1 | 33 | *p<0.001* |

### Surface Area (model with the regularized regions)

*Supplementary Table 15. Regression estimates for the model estimating how surface area of the regularized regions predict the cognitive factor*

| Path | | | Estimate | SE | *p* | Standardized Estimate |
|---|---|---|---|---|---|---|
| Cognitive_factor | ~ | bankssts_sa | -0.049 | 0.014 | *0.001* | -0.060 |
| Cognitive_factor | ~ | caudalmiddlefrontal_sa | 0.035 | 0.014 | *0.015* | 0.044 |
| Cognitive_factor | ~ | cuneus_sa | -0.014 | 0.016 | 0.366 | -0.019 |
| Cognitive_factor | ~ | entorhinal_sa | 0.071 | 0.012 | *p<0.001* | 0.089 |
| Cognitive_factor | ~ | fusiform_sa | 0.068 | 0.016 | *p<0.001* | 0.089 |
| Cognitive_factor | ~ | inferiortemporal_sa | 0.088 | 0.020 | *p<0.001* | 0.116 |
| Cognitive_factor | ~ | isthmuscingulate_sa | -0.035 | 0.014 | *0.011* | -0.044 |
| Cognitive_factor | ~ | lateralorbitofrontal_sa | 0.074 | 0.014 | *p<0.001* | 0.099 |
| Cognitive_factor | ~ | middletemporal_sa | 0.020 | 0.021 | 0.337 | 0.027 |
| Cognitive_factor | ~ | parahippocampal_sa | 0.003 | 0.012 | 0.828 | 0.003 |
| Cognitive_factor | ~ | paracentral_sa | -0.034 | 0.013 | *0.010* | -0.043 |
| Cognitive_factor | ~ | parsopercularis_sa | 0.045 | 0.014 | *0.001* | 0.055 |
| Cognitive_factor | ~ | parstriangularis_sa | -0.045 | 0.014 | *0.001* | -0.056 |

| | | | | | | |
|---|---|---|---|---|---|---|
| Cognitive_factor | ~ | pericalcarine_sa | 0.019 | 0.014 | 0.181 | 0.026 |
| Cognitive_factor | ~ | posteriorcingulate_sa | 0.016 | 0.015 | 0.292 | 0.020 |
| Cognitive_factor | ~ | precentral_sa | 0.023 | 0.015 | 0.135 | 0.030 |
| Cognitive_factor | ~ | rostralanteriorcingulate_sa | 0.036 | 0.014 | *0.011* | 0.044 |
| Cognitive_factor | ~ | temporalpole_sa | -0.088 | 0.013 | *p<0.001* | -0.110 |
| Cognitive_factor | ~ | insula_sa | 0.031 | 0.014 | *0.022* | 0.041 |

*Supplementary Figure 8. Standardized parameter estimates of how the surface area of each regularized region of interest together predict the cognitive factor*

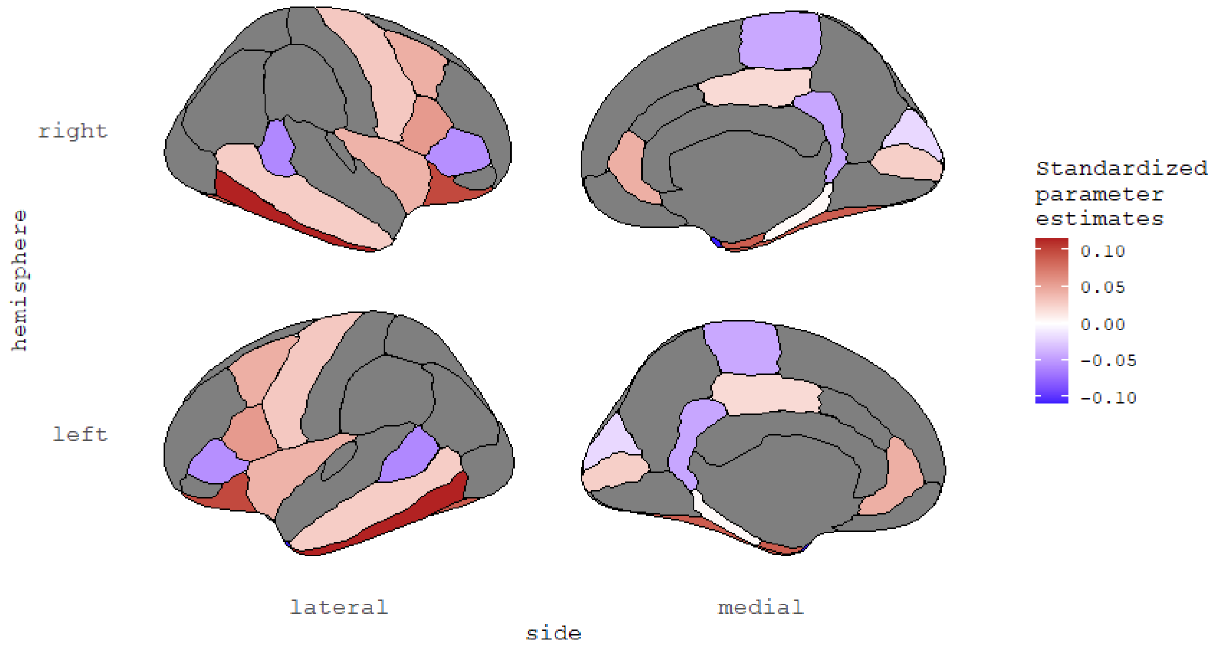


*Supplementary Table 16. Comparison of the model with free parameters and the model with parameters constrained to a same value for the surface area model with the regularized regions. AIC/BIC are two information criterions. Pr(>Chisq): p-value of the chi-square ratio test.*

| | **Degree of freedom** | **AIC** | **BIC** | **Difference in degree of freedom** | **Pr (>Chisq)** |
|---|---|---|---|---|---|
| Free model | 81 | 130604.7 | 130849.9 | NA | NA |
| Constrained model | 99 | 130780.3 | 130895.6 | 18 | *p<0.001* |

### Grey Matter Volume (model with all the regions)

*Supplementary Table 17. Regression estimates for the model estimating how grey matter volume of all the regions predict the cognitive factor*

| Path | | | Estimate | SE | *p* | Standardized Estimate |
|---|---|---|---|---|---|---|
| Cognitive_factor | ~ | bankssts_gmv | -0.038 | 0.014 | *0.008* | -0.045 |
| Cognitive_factor | ~ | caudalanteriorcingulate_gmv | 0.029 | 0.015 | 0.063 | 0.031 |
| Cognitive_factor | ~ | caudalmiddlefrontal_gmv | 0.019 | 0.013 | 0.152 | 0.024 |
| Cognitive_factor | ~ | cuneus_gmv | -0.012 | 0.016 | 0.445 | -0.016 |
| Cognitive_factor | ~ | entorhinal_gmv | 0.047 | 0.012 | *p<0.001* | 0.058 |
| Cognitive_factor | ~ | fusiform_gmv | 0.054 | 0.014 | *p<0.001* | 0.070 |
| Cognitive_factor | ~ | inferiorparietal_gmv | -0.028 | 0.014 | *0.050* | -0.036 |
| Cognitive_factor | ~ | inferiortemporal_gmv | 0.035 | 0.015 | *0.020* | 0.045 |
| Cognitive_factor | ~ | isthmuscingulate_gmv | -0.064 | 0.014 | *p<0.001* | -0.080 |
| Cognitive_factor | ~ | lateraloccipital_gmv | 0.047 | 0.014 | *0.001* | 0.061 |
| Cognitive_factor | ~ | lateralorbitofrontal_gmv | 0.064 | 0.017 | *p<0.001* | 0.086 |
| Cognitive_factor | ~ | lingual_gmv | 0.010 | 0.014 | 0.473 | 0.014 |
| Cognitive_factor | ~ | medialorbitofrontal_gmv | -0.074 | 0.015 | *p<0.001* | -0.095 |
| Cognitive_factor | ~ | middletemporal_gmv | 0.134 | 0.016 | *p<0.001* | 0.178 |
| Cognitive_factor | ~ | parahippocampal_gmv | 0.063 | 0.011 | *p<0.001* | 0.077 |
| Cognitive_factor | ~ | paracentral_gmv | -0.030 | 0.013 | *0.025* | -0.037 |
| Cognitive_factor | ~ | parsopercularis_gmv | 0.013 | 0.013 | 0.299 | 0.016 |
| Cognitive_factor | ~ | parsorbitalis_gmv | 0.014 | 0.014 | 0.324 | 0.017 |
| Cognitive_factor | ~ | parstriangularis_gmv | -0.035 | 0.014 | *0.010* | -0.043 |
| Cognitive_factor | ~ | pericalcarine_gmv | 0.009 | 0.015 | 0.555 | 0.012 |
| Cognitive_factor | ~ | postcentral_gmv | -0.022 | 0.014 | 0.106 | -0.029 |
| Cognitive_factor | ~ | posteriorcingulate_gmv | -0.024 | 0.014 | 0.095 | -0.029 |
| Cognitive_factor | ~ | precentral_gmv | 0.082 | 0.015 | *p<0.001* | 0.107 |
| Cognitive_factor | ~ | precuneus_gmv | 0.005 | 0.016 | 0.769 | 0.006 |
| Cognitive_factor | ~ | rostralanteriorcingulate_gmv | -0.006 | 0.015 | 0.691 | -0.007 |
| Cognitive_factor | ~ | rostralmiddlefrontal_gmv | -0.013 | 0.014 | 0.363 | -0.017 |
| Cognitive_factor | ~ | superiorfrontal_gmv | 0.011 | 0.016 | 0.502 | 0.015 |
| Cognitive_factor | ~ | superiorparietal_gmv | 0.015 | 0.014 | 0.294 | 0.020 |
| Cognitive_factor | ~ | superiortemporal_gmv | -0.010 | 0.016 | 0.546 | -0.013 |
| Cognitive_factor | ~ | supramarginal_gmv | -0.006 | 0.014 | 0.668 | -0.008 |
| Cognitive_factor | ~ | frontalpole_gmv | -0.008 | 0.012 | 0.499 | -0.010 |
| Cognitive_factor | ~ | temporalpole_gmv | 0.002 | 0.012 | 0.860 | 0.003 |
| Cognitive_factor | ~ | transversetemporal_gmv | 0.025 | 0.013 | 0.063 | 0.031 |
| Cognitive_factor | ~ | insula_gmv | 0.005 | 0.014 | 0.729 | 0.007 |

*Supplementary Figure 9. Standardized parameter estimates of how the grey matter volume of each region of interest together predict the cognitive factor*

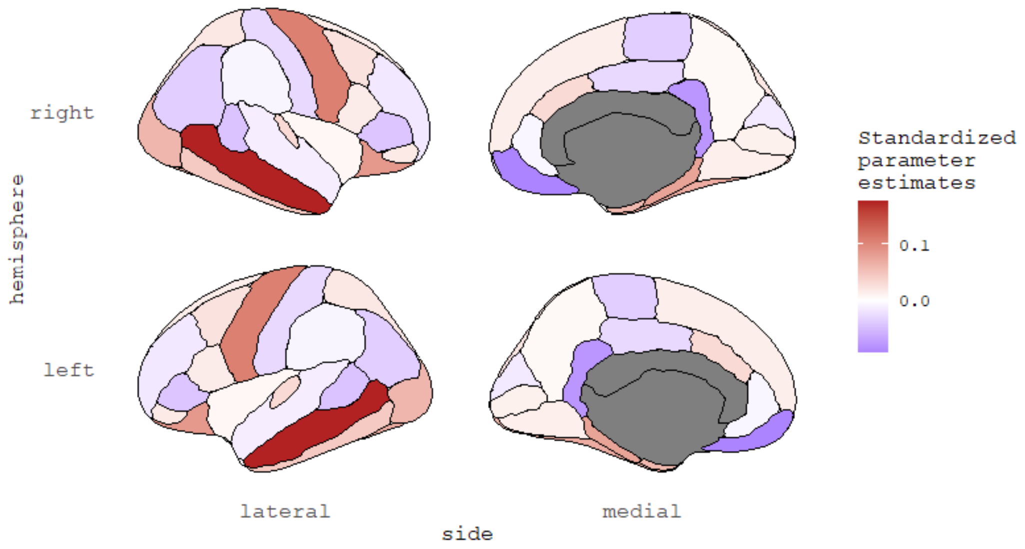


*Supplementary Table 18. Comparison of the model with free parameters and the model with parameters constrained to a same value for the grey matter volume model with all the regions. AIC/BIC are two information criterions. Pr(>Chisq): p-value of the chi-square ratio test.*

| | Degree of freedom | AIC | BIC | Difference in degree of freedom | Pr (>Chisq) |
|---|---|---|---|---|---|
| Free model | 141 | 130425.7 | 130779.0 | NA | NA |
| Constrained model | 174 | 130689.0 | 130804.4 | 33 | *p<0.001* |

### Grey Matter Volume (model with the regularized regions)

*Supplementary Table 19. Regression estimates for the model estimating how grey matter volume of the regularized regions predict the cognitive factor*

| Path | | | Estimate | SE | *p* | Standardized Estimate |
|---|---|---|---|---|---|---|
| Cognitive_factor | ~ | cuneus_gmv | 0.023 | 0.010 | *0.021* | 0.030 |
| Cognitive_factor | ~ | entorhinal_gmv | 0.056 | 0.011 | *p<0.001* | 0.069 |
| Cognitive_factor | ~ | fusiform_gmv | 0.060 | 0.013 | *p<0.001* | 0.077 |
| Cognitive_factor | ~ | isthmuscingulate_gmv | -0.054 | 0.012 | *p<0.001* | -0.068 |
| Cognitive_factor | ~ | middletemporal_gmv | 0.133 | 0.013 | *p<0.001* | 0.177 |
| Cognitive_factor | ~ | parahippocampal_gmv | 0.065 | 0.011 | *p<0.001* | 0.080 |
| Cognitive_factor | ~ | parstriangularis_gmv | -0.016 | 0.011 | 0.144 | -0.019 |
| Cognitive_factor | ~ | posteriorcingulate_gmv | -0.024 | 0.013 | 0.057 | -0.029 |
| Cognitive_factor | ~ | precentral_gmv | 0.091 | 0.012 | *p<0.001* | 0.118 |

| | | | | | | |
|---|---|---|---|---|---|---|
| Cognitive_factor | ~ | superiortemporal_gmv | -0.031 | 0.015 | *0.034* | -0.041 |
| Cognitive_factor | ~ | transversetemporal_gmv | 0.035 | 0.013 | *0.005* | 0.044 |

*Supplementary Figure 10. Standardized parameter estimates of how the grey matter volume of each regularized region of interest together predict the cognitive factor*

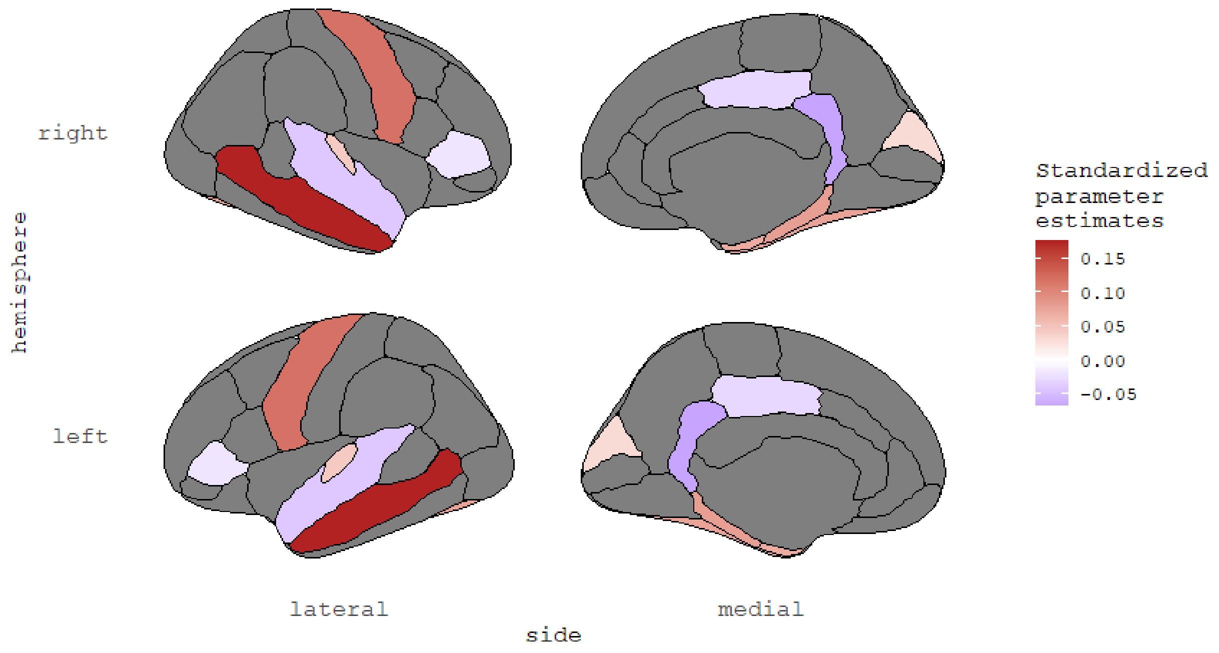


*Supplementary Table 20. Comparison of the model with free parameters and the model with parameters constrained to a same value for the grey matter volume model with the regularized regions. AIC/BIC are two information criterions. Pr(>Chisq): p-value of the chi-square ratio test.*

| | **Degree of freedom** | **AIC** | **BIC** | **Difference in degree of freedom** | **Pr (>Chisq)** |
|---|---|---|---|---|---|
| Free model | 49 | 130476.4 | 130663.8 | NA | NA |
| Constrained model | 59 | 130666.9 | 130782.3 | 10 | *p<0.001* |

### Fractional Anisotropy (model with all the regions)

*Supplementary Table 21. Regression estimates for the model estimating how fractional anisotropy of all the regions predict the cognitive factor*

| Path | | | Estimate | SE | *p* | Standardized Estimate |
|---|---|---|---|---|---|---|
| Cognitive_factor | ~ | fornix_fa | 0.092 | 0.016 | *p<0.001* | 0.120 |
| Cognitive_factor | ~ | cingulatecingulum_fa | -0.091 | 0.012 | *p<0.001* | -0.119 |
| Cognitive_factor | ~ | parahippocampalcingulum_fa | 0.046 | 0.012 | *p<0.001* | 0.061 |
| Cognitive_factor | ~ | corticospinalpyramidal_fa | 0.084 | 0.017 | *p<0.001* | 0.113 |
| Cognitive_factor | ~ | anteriorthalamicradiations_fa | 0.008 | 0.013 | 0.545 | 0.011 |
| Cognitive_factor | ~ | uncinate_fa | -0.047 | 0.017 | *0.007* | -0.062 |
| Cognitive_factor | ~ | inferiorlongitudinalfasiculus_fa | 0.021 | 0.016 | 0.181 | 0.027 |
| Cognitive_factor | ~ | inferiorfrontooccipitalfasiculus_fa | 0.053 | 0.018 | *0.003* | 0.071 |
| Cognitive_factor | ~ | forcepsmajor_fa | 0.038 | 0.018 | *0.035* | 0.054 |
| Cognitive_factor | ~ | forcepsminor_fa | -0.107 | 0.020 | *p<0.001* | -0.153 |
| Cognitive_factor | ~ | corpuscallosum_fa | 0.012 | 0.031 | 0.696 | 0.017 |
| Cognitive_factor | ~ | superiorlongitudinalfasiculus_fa | 0.316 | 0.136 | *0.020* | 0.424 |
| Cognitive_factor | ~ | temporalsuperiorlongitudinalfasiculus_fa | -0.195 | 0.073 | *0.007* | -0.260 |
| Cognitive_factor | ~ | parietalsuperiorlongitudinalfasiculus_fa | 0.035 | 0.079 | 0.659 | 0.046 |
| Cognitive_factor | ~ | superiorcorticostriate_fa | 0.879 | 0.116 | *p<0.001* | 1.162 |
| Cognitive_factor | ~ | superiorcorticostriatefrontalcortex_fa | -0.243 | 0.051 | *p<0.001* | -0.323 |
| Cognitive_factor | ~ | superiorcorticostriateparietalcortex_fa | -0.732 | 0.081 | *p<0.001* | -0.958 |
| Cognitive_factor | ~ | striatalinferiorfrontalcortex_fa | -0.080 | 0.016 | *p<0.001* | -0.104 |
| Cognitive_factor | ~ | inferiorfrontalsuperiorfrontalcortex_fa | 0.013 | 0.019 | 0.486 | 0.017 |
| Cognitive_factor | ~ | fornix_exfimbria_fa | -0.071 | 0.016 | *p<0.001* | -0.090 |

*Supplementary Table 22. Comparison of the model with free parameters and the model with parameters constrained to a same value for the fractional anisotropy model with all the regions. AIC/BIC are two information criterions. Pr(>Chisq): p-value of the chi-square ratio test.*

| | Degree of freedom | AIC | BIC | Difference in degree of freedom | Pr (>Chisq) |
|---|---|---|---|---|---|
| **Free model** | 85 | 123512.8 | 123763.3 | NA | NA |
| **Constrained model** | 104 | 123949.8 | 124064.3 | 19 | *p<0.001* |

### Fractional Anisotropy (model with the regularized regions)

*Supplementary Table 23. Regression estimates for the model estimating how fractional anisotropy of the regularized regions predict the cognitive factor*

| Path | | | Estimate | SE | *p* | Standardized Estimate |
|---|---|---|---|---|---|---|
| Cognitive_factor | ~ | forcepsmajor_fa | 0.039 | 0.012 | *0.001* | 0.056 |
| Cognitive_factor | ~ | forcepsminor_fa | -0.144 | 0.012 | *p<0.001* | -0.206 |
| Cognitive_factor | ~ | fornix_fa | 0.022 | 0.011 | *0.041* | 0.029 |
| Cognitive_factor | ~ | parahippocampalcingulum_fa | 0.042 | 0.011 | *p<0.001* | 0.057 |
| Cognitive_factor | ~ | corticospinalpyramidal_fa | 0.018 | 0.012 | 0.153 | 0.024 |
| Cognitive_factor | ~ | inferiorlongitudinalfasiculus_fa | 0.018 | 0.013 | 0.189 | 0.023 |
| Cognitive_factor | ~ | superiorlongitudinalfasiculus_fa | 0.128 | 0.013 | *p<0.001* | 0.173 |

*Supplementary Table 24. Comparison of the model with free parameters and the model with parameters constrained to a same value for the fractional anisotropy model with the regularized regions. AIC/BIC are two information criterions. Pr(>Chisq): p-value of the chi-square ratio test.*

| | Degree of freedom | AIC | BIC | Difference in degree of freedom | Pr (>Chisq) |
|---|---|---|---|---|---|
| **Free model** | 33 | 123715.0 | 123872.4 | NA | NA |
| **Constrained model** | 39 | 123924.8 | 124039.2 | 6 | *p<0.001* |

### Mean Diffusivity (model with all the regions)

*Supplementary Table 25. Regression estimates for the model estimating how mean diffusivity of all the regions predict the cognitive factor*

| Path | | | Estimate | SE | *p* | Standardized Estimate |
|---|---|---|---|---|---|---|
| Cognitive_factor | ~ | fornix_md | -0.070 | 0.023 | 0.002 | -0.095 |
| Cognitive_factor | ~ | cingulatecingulum_md | 0.022 | 0.016 | 0.175 | 0.025 |
| Cognitive_factor | ~ | parahippocampalcingulum_md | -0.029 | 0.015 | *0.049* | -0.039 |
| Cognitive_factor | ~ | corticospinalpyramidal_md | 0.009 | 0.016 | 0.569 | 0.012 |
| Cognitive_factor | ~ | anteriorthalamicradiations_md | -0.003 | 0.018 | 0.864 | -0.004 |
| Cognitive_factor | ~ | uncinate_md | -0.018 | 0.019 | 0.361 | -0.024 |
| Cognitive_factor | ~ | inferiorlongitudinalfasiculus_md | -0.061 | 0.021 | *0.003* | -0.079 |
| Cognitive_factor | ~ | inferiorfrontooccipitalfasiculus_md | 0.119 | 0.024 | *p<0.001* | 0.153 |
| Cognitive_factor | ~ | forcepsmajor_md | -0.013 | 0.018 | 0.456 | -0.019 |
| Cognitive_factor | ~ | forcepsminor_md | 0.085 | 0.018 | *p<0.001* | 0.123 |
| Cognitive_factor | ~ | corpuscallosum_md | -0.060 | 0.026 | *0.019* | -0.084 |
| Cognitive_factor | ~ | superiorlongitudinalfasiculus_md | -0.647 | 0.195 | *0.001* | -0.865 |

| | | | | | | |
|---|---|---|---|---|---|---|
| Cognitive_factor | ~ | temporalsuperiorlongitudinalfasiculus_md | 0.386 | 0.103 | *p<0.001* | 0.517 |
| Cognitive_factor | ~ | parietalsuperiorlongitudinalfasiculus_md | 0.179 | 0.109 | 0.102 | 0.240 |
| Cognitive_factor | ~ | superiorcorticostriate_md | 0.286 | 0.149 | 0.055 | 0.374 |
| Cognitive_factor | ~ | superiorcorticostriatefrontalcortex_md | 0.008 | 0.070 | 0.907 | 0.011 |
| Cognitive_factor | ~ | superiorcorticostriateparietalcortex_md | -0.306 | 0.103 | *0.003* | -0.400 |
| Cognitive_factor | ~ | striatalinferiorfrontalcortex_md | -0.044 | 0.025 | 0.078 | -0.055 |
| Cognitive_factor | ~ | inferiorfrontalsuperiorfrontalcortex_md | 0.063 | 0.031 | *0.038* | 0.084 |
| Cognitive_factor | ~ | fornix_exfimbria_md | 0.075 | 0.021 | *p<0.001* | 0.101 |

*Supplementary Table 26. Comparison of the model with free parameters and the model with parameters constrained to a same value for the mean diffusivity model with all the regions. AIC/BIC are two information criterions. Pr(>Chisq): p-value of the chi-square ratio test.*

| | Degree of freedom | AIC | BIC | Difference in degree of freedom | Pr (>Chisq) |
|---|---|---|---|---|---|
| **Free model** | 85 | 123885.5 | 124135.9 | NA | NA |
| **Constrained model** | 104 | 124018.4 | 124132.8 | 19 | *p<0.001* |

### Mean Diffusivity (model with the regularized regions)

*Supplementary Table 27. Regression estimates for the model estimating how mean diffusivity of the regularized regions predict the cognitive factor*

| Path | | | Estimate | SE | *p* | Standardized Estimate |
|---|---|---|---|---|---|---|
| Cognitive_factor | ~ | forcepsmajor_md | -0.030 | 0.013 | *0.025* | -0.043 |
| Cognitive_factor | ~ | forcepsminor_md | 0.055 | 0.012 | *p<0.001* | 0.080 |
| Cognitive_factor | ~ | cingulatecingulum_md | 0.024 | 0.015 | 0.118 | 0.029 |
| Cognitive_factor | ~ | corticospinalpyramidal_md | -0.010 | 0.014 | 0.456 | -0.014 |
| Cognitive_factor | ~ | parietalsuperiorlongitudinalfasiculus_md | -0.067 | 0.016 | *p<0.001* | -0.093 |
| Cognitive_factor | ~ | fornix_exfimbria_md | 0.002 | 0.010 | 0.827 | 0.003 |

*Supplementary Table 28. Comparison of the model with free parameters and the model with parameters constrained to a same value for the mean diffusivity model with the regularized regions. AIC/BIC are two information criterions. Pr(>Chisq): p-value of the chi-square ratio test.*

| | Degree of freedom | AIC | BIC | Difference in degree of freedom | Pr (>Chisq) |
|---|---|---|---|---|---|
| **Free model** | 29 | 123990.0 | 124140.3 | NA | NA |
| **Constrained model** | 34 | 124040.9 | 124155.4 | 5 | *p<0.001* |

### White Matter Volume (model with all the regions)

*Supplementary Table 29. Regression estimates for the model estimating how white matter volume of all the regions predict the cognitive factor*

| Path | | | Estimate | SE | *p* | Standardized Estimate |
|---|---|---|---|---|---|---|
| Cognitive_factor | ~ | fornix_wmv | 0.174 | 0.028 | *p<0.001* | 0.240 |
| Cognitive_factor | ~ | cingulatecingulum_wmv | -0.084 | 0.016 | *p<0.001* | -0.112 |
| Cognitive_factor | ~ | parahippocampalcingulum_wmv | 0.002 | 0.013 | 0.879 | 0.003 |
| Cognitive_factor | ~ | corticospinalpyramidal_wmv | 0.210 | 0.024 | *p<0.001* | 0.287 |
| Cognitive_factor | ~ | anteriorthalamicradiations_wmv | 0.074 | 0.024 | *0.002* | 0.103 |
| Cognitive_factor | ~ | uncinate_wmv | -0.029 | 0.016 | 0.073 | -0.039 |
| Cognitive_factor | ~ | inferiorlongitudinalfasiculus_wmv | 0.020 | 0.017 | 0.245 | 0.026 |
| Cognitive_factor | ~ | inferiorfrontooccipitalfasiculus_wmv | 0.015 | 0.027 | 0.569 | 0.021 |
| Cognitive_factor | ~ | forcepsmajor_wmv | -0.027 | 0.019 | 0.158 | -0.038 |
| Cognitive_factor | ~ | forcepsminor_wmv | -0.079 | 0.026 | *0.002* | -0.112 |
| Cognitive_factor | ~ | corpuscallosum_wmv | 0.095 | 0.040 | *0.017* | 0.134 |
| Cognitive_factor | ~ | superiorlongitudinalfasiculus_wmv | -0.020 | 0.081 | 0.809 | -0.027 |
| Cognitive_factor | ~ | temporalsuperiorlongitudinalfasiculus_wmv | 0.113 | 0.054 | *0.037* | 0.151 |
| Cognitive_factor | ~ | parietalsuperiorlongitudinalfasiculus_wmv | -0.017 | 0.043 | 0.691 | -0.023 |
| Cognitive_factor | ~ | superiorcorticostriate_wmv | -0.082 | 0.087 | 0.348 | -0.111 |
| Cognitive_factor | ~ | superiorcorticostriatefrontalcortex_wmv | -0.153 | 0.051 | *0.003* | -0.209 |
| Cognitive_factor | ~ | superiorcorticostriateparietalcortex_wmv | -0.009 | 0.056 | 0.870 | -0.012 |
| Cognitive_factor | ~ | striatalinferiorfrontalcortex_wmv | -0.050 | 0.019 | *0.008* | -0.067 |
| Cognitive_factor | ~ | inferiorfrontalsuperiorfrontalcortex_wmv | 0.101 | 0.023 | *p<0.001* | 0.136 |
| Cognitive_factor | ~ | fornix_exfimbria_wmv | -0.034 | 0.027 | 0.204 | -0.045 |

*Supplementary Table 30. Comparison of the model with free parameters and the model with parameters constrained to a same value for the white matter volume model with all the regions. AIC/BIC are two information criterions. Pr(>Chisq): p-value of the chi-square ratio test.*

| | Degree of freedom | AIC | BIC | Difference in degree of freedom | Pr (>Chisq) |
|---|---|---|---|---|---|
| **Free model** | 85 | 123223.1 | 123473.5 | NA | NA |
| **Constrained model** | 104 | 123498.9 | 123613.4 | 19 | *p<0.001* |

### White Matter Volume (model with the regularized regions)

*Supplementary Table 31. Regression estimates for the model estimating how white matter volume of the regularized regions predict the cognitive factor*

| Path | | | Estimate | SE | *p* | Standardized Estimate |
|---|---|---|---|---|---|---|
| Cognitive_factor | ~ | forcepsmajor_wmv | 0.008 | 0.013 | 0.550 | 0.011 |
| Cognitive_factor | ~ | fornix_wmv | 0.150 | 0.016 | *p<0.001* | 0.207 |
| Cognitive_factor | ~ | cingulatecingulum_wmv | -0.089 | 0.015 | *p<0.001* | -0.118 |
| Cognitive_factor | ~ | parahippocampalcingulum_wmv | 0.004 | 0.012 | 0.719 | 0.006 |
| Cognitive_factor | ~ | corticospinalpyramidal_wmv | 0.117 | 0.017 | *p<0.001* | 0.160 |
| Cognitive_factor | ~ | anteriorthalamicradiations_wmv | -0.007 | 0.018 | 0.715 | -0.009 |
| Cognitive_factor | ~ | uncinate_wmv | -0.029 | 0.015 | 0.057 | -0.039 |
| Cognitive_factor | ~ | inferiorlongitudinalfasiculus_wmv | 0.028 | 0.016 | 0.080 | 0.038 |
| Cognitive_factor | ~ | inferiorfrontooccipitalfasiculus_wmv | -0.018 | 0.025 | 0.477 | -0.024 |
| Cognitive_factor | ~ | temporalsuperiorlongitudinalfasiculus_wmv | 0.067 | 0.017 | *p<0.001* | 0.091 |

*Supplementary Table 32. Comparison of the model with free parameters and the model with parameters constrained to a same value for the white matter volume model with the regularized regions. AIC/BIC are two information criterions. Pr(>Chisq): p-value of the chi-square ratio test.*

| | Degree of freedom | AIC | BIC | Difference in degree of freedom | Pr (>Chisq) |
|---|---|---|---|---|---|
| **Free model** | 45 | 123280.8 | 123459.7 | NA | NA |
| **Constrained model** | 54 | 123486.0 | 123600.5 | 9 | *p<0.001* |

### Models per tissue estimating cognitive factor from several regions in three metrics

For grey and white matter, we reported the tables for the models with all the regions included as predictors and the models with only the regions that survived the regularization (the one that we used for the analyses).

#### Grey matter metrics (model with all the regions)

*Supplementary Table 33. Regression estimates for the model estimating how cortical thickness, surface area and grey matter volume of all the regions predict the cognitive factor*

| Path | | | Estimate | SE | *p* | Standardized Estimate |
|---|---|---|---|---|---|---|
| Cognitive_factor | ~ | bankssts_ct | -0.002 | 0.018 | 0.911 | -0.002 |
| Cognitive_factor | ~ | caudalanteriorcingulate_ct | -0.045 | 0.019 | *0.015* | -0.052 |
| Cognitive_factor | ~ | caudalmiddlefrontal_ct | 0.067 | 0.029 | *0.022* | 0.085 |
| Cognitive_factor | ~ | cuneus_ct | 0.010 | 0.030 | 0.753 | 0.012 |
| Cognitive_factor | ~ | entorhinal_ct | 0.021 | 0.019 | 0.273 | 0.025 |
| Cognitive_factor | ~ | fusiform_ct | 0.009 | 0.025 | 0.723 | 0.011 |
| Cognitive_factor | ~ | inferiorparietal_ct | -0.037 | 0.032 | 0.249 | -0.047 |
| Cognitive_factor | ~ | inferiortemporal_ct | 0.060 | 0.029 | *0.039* | 0.076 |
| Cognitive_factor | ~ | isthmuscingulate_ct | 0.016 | 0.019 | 0.404 | 0.019 |
| Cognitive_factor | ~ | lateraloccipital_ct | 0.025 | 0.037 | 0.500 | 0.033 |
| Cognitive_factor | ~ | lateralorbitofrontal_ct | -0.079 | 0.022 | *p<0.001* | -0.098 |
| Cognitive_factor | ~ | lingual_ct | 0.063 | 0.032 | *0.046* | 0.081 |
| Cognitive_factor | ~ | medialorbitofrontal_ct | 0.054 | 0.023 | *0.020* | 0.066 |
| Cognitive_factor | ~ | middletemporal_ct | -0.059 | 0.034 | 0.080 | -0.076 |
| Cognitive_factor | ~ | parahippocampal_ct | 0.037 | 0.028 | 0.184 | 0.047 |
| Cognitive_factor | ~ | paracentral_ct | 0.011 | 0.027 | 0.697 | 0.013 |
| Cognitive_factor | ~ | parsopercularis_ct | -0.044 | 0.027 | 0.100 | -0.053 |
| Cognitive_factor | ~ | parsorbitalis_ct | 0.014 | 0.023 | 0.544 | 0.017 |
| Cognitive_factor | ~ | parstriangularis_ct | -0.043 | 0.027 | 0.117 | -0.051 |
| Cognitive_factor | ~ | pericalcarine_ct | 0.100 | 0.029 | *0.001* | 0.128 |
| Cognitive_factor | ~ | postcentral_ct | 0.048 | 0.036 | 0.181 | 0.063 |
| Cognitive_factor | ~ | posteriorcingulate_ct | -0.048 | 0.022 | *0.028* | -0.057 |
| Cognitive_factor | ~ | precentral_ct | 0.173 | 0.038 | *p<0.001* | 0.223 |
| Cognitive_factor | ~ | precuneus_ct | -0.064 | 0.026 | *0.013* | -0.083 |
| Cognitive_factor | ~ | rostralanteriorcingulate_ct | -0.053 | 0.020 | *0.006* | -0.061 |
| Cognitive_factor | ~ | rostralmiddlefrontal_ct | 0.114 | 0.030 | *p<0.001* | 0.146 |
| Cognitive_factor | ~ | superiorfrontal_ct | -0.004 | 0.032 | 0.908 | -0.005 |
| Cognitive_factor | ~ | superiorparietal_ct | -0.098 | 0.034 | *0.004* | -0.130 |
| Cognitive_factor | ~ | superiortemporal_ct | -0.074 | 0.035 | *0.035* | -0.095 |
| Cognitive_factor | ~ | supramarginal_ct | 0.051 | 0.032 | 0.106 | 0.065 |
| Cognitive_factor | ~ | frontalpole_ct | -0.001 | 0.022 | 0.979 | -0.001 |
| Cognitive_factor | ~ | temporalpole_ct | -0.042 | 0.021 | *0.048* | -0.051 |
| Cognitive_factor | ~ | transversetemporal_ct | -0.056 | 0.019 | *0.003* | -0.067 |

| | | | | | | |
|---|---|---|---|---|---|---|
| Cognitive_factor | ~ | insula_ct | -0.047 | 0.027 | 0.079 | -0.058 |
| Cognitive_factor | ~ | bankssts_sa | -0.060 | 0.046 | 0.189 | -0.072 |
| Cognitive_factor | ~ | caudalanteriorcingulate_sa | -0.025 | 0.054 | 0.642 | -0.028 |
| Cognitive_factor | ~ | caudalmiddlefrontal_sa | 0.219 | 0.070 | *0.002* | 0.275 |
| Cognitive_factor | ~ | cuneus_sa | -0.034 | 0.048 | 0.475 | -0.045 |
| Cognitive_factor | ~ | entorhinal_sa | 0.022 | 0.032 | 0.493 | 0.027 |
| Cognitive_factor | ~ | fusiform_sa | 0.101 | 0.056 | 0.069 | 0.131 |
| Cognitive_factor | ~ | inferiorparietal_sa | 0.002 | 0.069 | 0.973 | 0.003 |
| Cognitive_factor | ~ | inferiortemporal_sa | 0.289 | 0.065 | *p<0.001* | 0.376 |
| Cognitive_factor | ~ | isthmuscingulate_sa | 0.007 | 0.050 | 0.895 | 0.008 |
| Cognitive_factor | ~ | lateraloccipital_sa | -0.097 | 0.066 | 0.144 | -0.126 |
| Cognitive_factor | ~ | lateralorbitofrontal_sa | -0.137 | 0.054 | *0.012* | -0.180 |
| Cognitive_factor | ~ | lingual_sa | 0.115 | 0.058 | *0.048* | 0.152 |
| Cognitive_factor | ~ | medialorbitofrontal_sa | 0.221 | 0.049 | *p<0.001* | 0.283 |
| Cognitive_factor | ~ | middletemporal_sa | -0.225 | 0.062 | *p<0.001* | -0.298 |
| Cognitive_factor | ~ | parahippocampal_sa | -0.062 | 0.038 | 0.106 | -0.077 |
| Cognitive_factor | ~ | paracentral_sa | 0.072 | 0.053 | 0.176 | 0.090 |
| Cognitive_factor | ~ | parsopercularis_sa | 0.150 | 0.071 | *0.034* | 0.183 |
| Cognitive_factor | ~ | parsorbitalis_sa | -0.020 | 0.045 | 0.652 | -0.026 |
| Cognitive_factor | ~ | parstriangularis_sa | -0.037 | 0.065 | 0.567 | -0.046 |
| Cognitive_factor | ~ | pericalcarine_sa | 0.215 | 0.050 | *p<0.001* | 0.288 |
| Cognitive_factor | ~ | postcentral_sa | -0.032 | 0.068 | 0.639 | -0.041 |
| Cognitive_factor | ~ | posteriorcingulate_sa | -0.112 | 0.062 | 0.072 | -0.136 |
| Cognitive_factor | ~ | precentral_sa | 0.143 | 0.068 | *0.035* | 0.185 |
| Cognitive_factor | ~ | precuneus_sa | -0.078 | 0.066 | 0.233 | -0.104 |
| Cognitive_factor | ~ | rostralanteriorcingulate_sa | -0.129 | 0.049 | *0.008* | -0.158 |
| Cognitive_factor | ~ | rostralmiddlefrontal_sa | 0.307 | 0.065 | *p<0.001* | 0.401 |
| Cognitive_factor | ~ | superiorfrontal_sa | -0.043 | 0.077 | 0.577 | -0.057 |
| Cognitive_factor | ~ | superiorparietal_sa | -0.027 | 0.067 | 0.685 | -0.035 |
| Cognitive_factor | ~ | superiortemporal_sa | -0.107 | 0.065 | 0.098 | -0.141 |
| Cognitive_factor | ~ | supramarginal_sa | 0.028 | 0.072 | 0.694 | 0.036 |
| Cognitive_factor | ~ | frontalpole_sa | 0.031 | 0.027 | 0.263 | 0.037 |
| Cognitive_factor | ~ | temporalpole_sa | -0.128 | 0.027 | *p<0.001* | -0.159 |
| Cognitive_factor | ~ | transversetemporal_sa | -0.155 | 0.034 | *p<0.001* | -0.193 |
| Cognitive_factor | ~ | insula_sa | -0.090 | 0.054 | 0.096 | -0.117 |
| Cognitive_factor | ~ | bankssts_gmv | 0.028 | 0.047 | 0.551 | 0.033 |
| Cognitive_factor | ~ | caudalanteriorcingulate_gmv | 0.068 | 0.055 | 0.218 | 0.072 |
| Cognitive_factor | ~ | caudalmiddlefrontal_gmv | -0.194 | 0.067 | *0.004* | -0.240 |
| Cognitive_factor | ~ | cuneus_gmv | 0.016 | 0.057 | 0.775 | 0.021 |
| Cognitive_factor | ~ | entorhinal_gmv | 0.014 | 0.029 | 0.637 | 0.017 |
| Cognitive_factor | ~ | fusiform_gmv | -0.047 | 0.054 | 0.383 | -0.060 |
| Cognitive_factor | ~ | inferiorparietal_gmv | -0.004 | 0.067 | 0.957 | -0.005 |
| Cognitive_factor | ~ | inferiortemporal_gmv | -0.187 | 0.062 | *0.003* | -0.240 |
| Cognitive_factor | ~ | isthmuscingulate_gmv | -0.057 | 0.049 | 0.246 | -0.071 |

| | | | | | | |
|---|---|---|---|---|---|---|
| Cognitive_factor | ~ | lateraloccipital_gmv | 0.116 | 0.071 | 0.102 | 0.151 |
| Cognitive_factor | ~ | lateralorbitofrontal_gmv | 0.158 | 0.051 | *0.002* | 0.210 |
| Cognitive_factor | ~ | lingual_gmv | -0.131 | 0.065 | *0.042* | -0.171 |
| Cognitive_factor | ~ | medialorbitofrontal_gmv | -0.243 | 0.050 | *p<0.001* | -0.306 |
| Cognitive_factor | ~ | middletemporal_gmv | 0.271 | 0.056 | *p<0.001* | 0.356 |
| Cognitive_factor | ~ | parahippocampal_gmv | 0.087 | 0.042 | *0.038* | 0.107 |
| Cognitive_factor | ~ | paracentral_gmv | -0.093 | 0.054 | 0.084 | -0.114 |
| Cognitive_factor | ~ | parsopercularis_gmv | -0.117 | 0.067 | 0.080 | -0.142 |
| Cognitive_factor | ~ | parsorbitalis_gmv | 0.032 | 0.042 | 0.445 | 0.040 |
| Cognitive_factor | ~ | parstriangularis_gmv | 0.004 | 0.062 | 0.945 | 0.005 |
| Cognitive_factor | ~ | pericalcarine_gmv | -0.243 | 0.056 | *p<0.001* | -0.324 |
| Cognitive_factor | ~ | postcentral_gmv | -0.033 | 0.069 | 0.635 | -0.042 |
| Cognitive_factor | ~ | posteriorcingulate_gmv | 0.084 | 0.060 | 0.162 | 0.100 |
| Cognitive_factor | ~ | precentral_gmv | -0.095 | 0.066 | 0.150 | -0.123 |
| Cognitive_factor | ~ | precuneus_gmv | 0.064 | 0.064 | 0.319 | 0.085 |
| Cognitive_factor | ~ | rostralanteriorcingulate_gmv | 0.119 | 0.048 | *0.013* | 0.140 |
| Cognitive_factor | ~ | rostralmiddlefrontal_gmv | -0.290 | 0.063 | *p<0.001* | -0.376 |
| Cognitive_factor | ~ | superiorfrontal_gmv | 0.081 | 0.068 | 0.234 | 0.106 |
| Cognitive_factor | ~ | superiorparietal_gmv | 0.062 | 0.067 | 0.355 | 0.080 |
| Cognitive_factor | ~ | superiortemporal_gmv | 0.155 | 0.061 | *0.012* | 0.201 |
| Cognitive_factor | ~ | supramarginal_gmv | -0.046 | 0.068 | 0.493 | -0.058 |
| Cognitive_factor | ~ | frontalpole_gmv | -0.001 | 0.029 | 0.970 | -0.001 |
| Cognitive_factor | ~ | temporalpole_gmv | 0.105 | 0.029 | *p<0.001* | 0.128 |
| Cognitive_factor | ~ | transversetemporal_gmv | 0.143 | 0.032 | *p<0.001* | 0.177 |
| Cognitive_factor | ~ | insula_gmv | 0.106 | 0.054 | *0.048* | 0.142 |

*Supplementary Figure 11. P-value of the models estimating how the grey matter metrics of each region of interest together predict the cognitive factor*

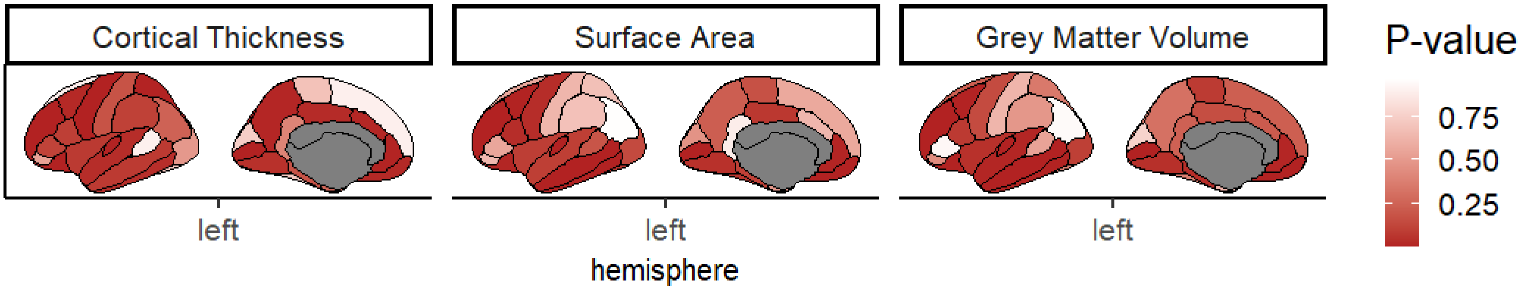


*Supplementary Figure 12. Standardized parameter estimates of how the grey matter metrics of each region of interest together predict the cognitive factor*

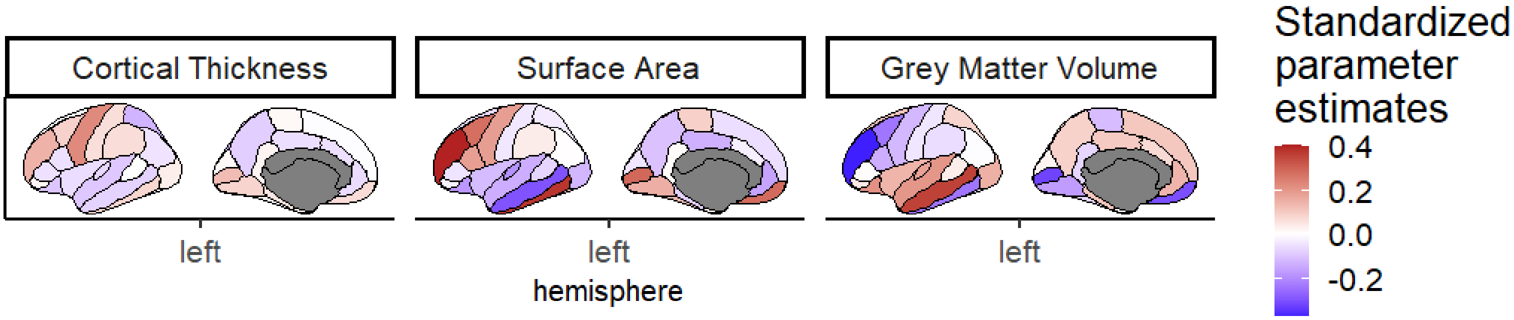


*Supplementary Table 34. Comparison of the model with free parameters and the model with parameters constrained to a same value for the grey matter model with all the regions. AIC/BIC are two information criterions. Pr(>Chisq): p-value of the chi-square ratio test.*

| | Degree of freedom | AIC | BIC | Difference in degree of freedom | Pr (>Chisq) |
|---|---|---|---|---|---|
| **Free model** | 413 | 129973.5 | 130817.1 | NA | NA |
| **Constrained model** | 514 | 130717.2 | 130832.5 | 101 | *p<0.001* |

### Grey matter metrics (model with the regularized regions)

*Supplementary Table 35. Regression estimates for the model estimating how cortical thickness, surface area and grey matter volume of the regularized regions predict the cognitive factor*

| Path | | | Estimate | SE | *p* | Standardized Estimate |
|---|---|---|---|---|---|---|
| Cognitive_factor | ~ | bankssts_ct | -0.017 | 0.013 | 0.182 | -0.019 |
| Cognitive_factor | ~ | caudalanteriorcingulate_ct | -0.053 | 0.011 | *p<0.001* | -0.061 |
| Cognitive_factor | ~ | cuneus_ct | 0.001 | 0.012 | 0.944 | 0.001 |
| Cognitive_factor | ~ | entorhinal_ct | 0.016 | 0.010 | 0.119 | 0.019 |
| Cognitive_factor | ~ | fusiform_ct | -0.008 | 0.014 | 0.574 | -0.010 |
| Cognitive_factor | ~ | isthmuscingulate_ct | -0.021 | 0.011 | *0.050* | -0.025 |
| Cognitive_factor | ~ | lateraloccipital_ct | 0.061 | 0.014 | *p<0.001* | 0.082 |
| Cognitive_factor | ~ | lateralorbitofrontal_ct | -0.019 | 0.013 | 0.140 | -0.023 |
| Cognitive_factor | ~ | medialorbitofrontal_ct | -0.052 | 0.012 | *p<0.001* | -0.063 |
| Cognitive_factor | ~ | middletemporal_ct | 0.055 | 0.017 | *0.001* | 0.070 |
| Cognitive_factor | ~ | paracentral_ct | -0.019 | 0.013 | 0.151 | -0.024 |
| Cognitive_factor | ~ | parstriangularis_ct | -0.055 | 0.013 | *p<0.001* | -0.067 |
| Cognitive_factor | ~ | rostralmiddlefrontal_ct | 0.131 | 0.023 | *p<0.001* | 0.168 |
| Cognitive_factor | ~ | superiorfrontal_ct | 0.013 | 0.017 | 0.457 | 0.017 |
| Cognitive_factor | ~ | supramarginal_ct | 0.009 | 0.015 | 0.549 | 0.011 |
| Cognitive_factor | ~ | frontalpole_ct | -0.024 | 0.014 | 0.082 | -0.028 |
| Cognitive_factor | ~ | transversetemporal_ct | 0.014 | 0.011 | 0.205 | 0.017 |
| Cognitive_factor | ~ | caudalanteriorcingulate_sa | 0.041 | 0.017 | *0.015* | 0.046 |
| Cognitive_factor | ~ | cuneus_sa | -0.012 | 0.016 | 0.463 | -0.015 |
| Cognitive_factor | ~ | fusiform_sa | 0.068 | 0.015 | *p<0.001* | 0.089 |
| Cognitive_factor | ~ | inferiortemporal_sa | 0.088 | 0.019 | *p<0.001* | 0.115 |
| Cognitive_factor | ~ | isthmus`ingulate_sa | -0.051 | 0.014 | *p<0.001* | -0.065 |
| Cognitive_factor | ~ | lingual_sa | 0.010 | 0.016 | 0.517 | 0.014 |
| Cognitive_factor | ~ | medialorbitofrontal_sa | -0.006 | 0.014 | 0.690 | -0.007 |
| Cognitive_factor | ~ | parahippocampal_sa | -0.150 | 0.017 | *p<0.001* | -0.187 |
| Cognitive_factor | ~ | pericalcarine_sa | 0.010 | 0.018 | 0.572 | 0.014 |
| Cognitive_factor | ~ | posteriorcingulate_sa | -0.044 | 0.016 | *0.005* | -0.054 |
| Cognitive_factor | ~ | precentral_sa | -0.064 | 0.028 | *0.021* | -0.083 |
| Cognitive_factor | ~ | rostralmiddlefrontal_sa | 0.349 | 0.046 | *p<0.001* | 0.459 |
| Cognitive_factor | ~ | superiorfrontal_sa | 0.029 | 0.018 | 0.108 | 0.039 |
| Cognitive_factor | ~ | superiorparietal_sa | 0.029 | 0.012 | *0.017* | 0.038 |
| Cognitive_factor | ~ | temporalpole_sa | -0.045 | 0.013 | *0.001* | -0.056 |
| Cognitive_factor | ~ | transversetemporal_sa | -0.015 | 0.012 | 0.212 | -0.019 |
| Cognitive_factor | ~ | insula_sa | 0.001 | 0.023 | 0.979 | 0.001 |
| Cognitive_factor | ~ | caudalmiddlefrontal_gmv | 0.024 | 0.013 | 0.073 | 0.030 |
| Cognitive_factor | ~ | middletemporal_gmv | 0.053 | 0.017 | *0.002* | 0.070 |
| Cognitive_factor | ~ | parahippocampal_gmv | 0.176 | 0.015 | *p<0.001* | 0.217 |
| Cognitive_factor | ~ | parsorbitalis_gmv | 0.020 | 0.012 | 0.103 | 0.025 |

| | | | | | | |
|---|---|---|---|---|---|---|
| Cognitive_factor | ~ | precentral_gmv | 0.117 | 0.026 | *p<0.001* | 0.152 |
| Cognitive_factor | ~ | rostralanteriorcingulate_gmv | -0.011 | 0.015 | 0.463 | -0.013 |
| Cognitive_factor | ~ | rostralmiddlefrontal_gmv | -0.365 | 0.046 | *p<0.001* | -0.475 |
| Cognitive_factor | ~ | frontalpole_gmv | 0.038 | 0.015 | *0.012* | 0.046 |
| Cognitive_factor | ~ | insula_gmv | 0.022 | 0.023 | 0.350 | 0.029 |

*Supplementary Figure 13. P-value of the models estimating how the grey matter metrics of each regularized region of interest together predict the cognitive factor*

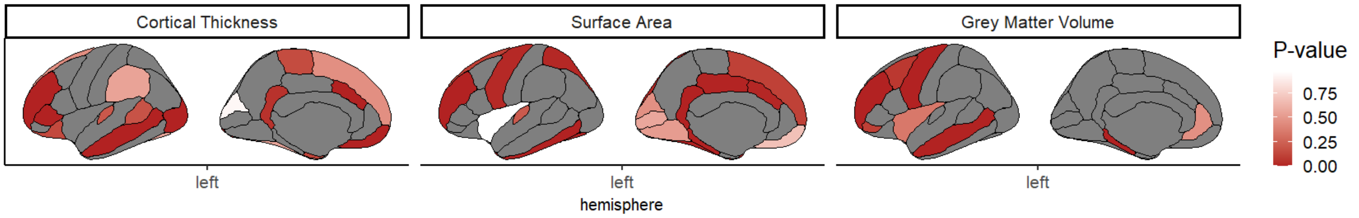


*Supplementary Figure 14. Standardized parameter estimates of how the grey matter metrics of each regularized region of interest together predict the cognitive factor*

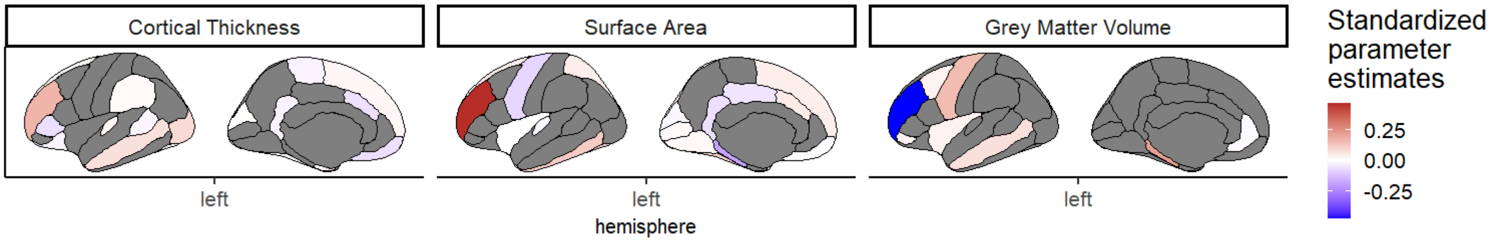


*Supplementary Table 36. Comparison of the model with free parameters and the model with parameters constrained to a same value for the grey matter model with the regularized regions. AIC/BIC are two information criterions. Pr(>Chisq): p-value of the chi-square ratio test.*

| | Degree of freedom | AIC | BIC | Difference in degree of freedom | Pr (>Chisq) |
|---|---|---|---|---|---|
| **Free model** | 177 | 130220.1 | 130638.2 | NA | NA |
| **Constrained model** | 219 | 130757.3 | 130872.6 | 42 | *p<0.001* |

### White matter metrics (model with all the regions)

*Supplementary Table 37. Regression estimates for the model estimating how fractional anisotropy, mean diffusivity and white matter volume of all the regions predict the cognitive factor*

| Path | | | Estimate | SE | *p* | Standardized Estimate |
|---|---|---|---|---|---|---|
| Cognitive_factor | ~ | fornix_fa | 0.023 | 0.019 | 0.216 | 0.030 |
| Cognitive_factor | ~ | cingulatecingulum_fa | -0.095 | 0.015 | *p<0.001* | -0.121 |
| Cognitive_factor | ~ | parahippocampalcingulum_fa | 0.013 | 0.013 | 0.312 | 0.017 |
| Cognitive_factor | ~ | corticospinalpyramidal_fa | 0.061 | 0.019 | *0.002* | 0.080 |
| Cognitive_factor | ~ | anteriorthalamicradiations_fa | 0.038 | 0.017 | *0.021* | 0.050 |
| Cognitive_factor | ~ | uncinate_fa | -0.071 | 0.020 | *p<0.001* | -0.092 |
| Cognitive_factor | ~ | inferiorlongitudinalfasiculus_fa | 0.018 | 0.019 | 0.332 | 0.023 |
| Cognitive_factor | ~ | inferiorfrontooccipitalfasiculus_fa | 0.079 | 0.023 | *0.001* | 0.103 |
| Cognitive_factor | ~ | forcepsmajor_fa | 0.004 | 0.022 | 0.858 | 0.006 |
| Cognitive_factor | ~ | forcepsminor_fa | -0.056 | 0.025 | *0.023* | -0.078 |
| Cognitive_factor | ~ | corpuscallosum_fa | -0.044 | 0.038 | 0.246 | -0.061 |
| Cognitive_factor | ~ | superiorlongitudinalfasiculus_fa | -0.171 | 0.143 | 0.230 | -0.225 |
| Cognitive_factor | ~ | temporalsuperiorlongitudinalfasiculus_fa | 0.079 | 0.077 | 0.299 | 0.104 |
| Cognitive_factor | ~ | parietalsuperiorlongitudinalfasiculus_fa | 0.230 | 0.085 | *0.007* | 0.301 |
| Cognitive_factor | ~ | superiorcorticostriate_fa | 0.542 | 0.139 | *p<0.001* | 0.700 |
| Cognitive_factor | ~ | superiorcorticostriatefrontalcortex_fa | -0.146 | 0.064 | *0.023* | -0.189 |
| Cognitive_factor | ~ | superiorcorticostriateparietalcortex_fa | -0.448 | 0.093 | *p<0.001* | -0.573 |
| Cognitive_factor | ~ | striatalinferiorfrontalcortex_fa | -0.054 | 0.018 | *0.002* | -0.068 |
| Cognitive_factor | ~ | inferiorfrontalsuperiorfrontalcortex_fa | 0.064 | 0.022 | *0.004* | 0.083 |
| Cognitive_factor | ~ | fornix_exfimbria_fa | -0.047 | 0.018 | *0.009* | -0.059 |
| Cognitive_factor | ~ | fornix_md | -0.005 | 0.026 | 0.844 | -0.007 |
| Cognitive_factor | ~ | cingulatecingulum_md | 0.015 | 0.017 | 0.354 | 0.018 |
| Cognitive_factor | ~ | parahippocampalcingulum_md | -0.012 | 0.015 | 0.445 | -0.015 |
| Cognitive_factor | ~ | corticospinalpyramidal_md | 0.111 | 0.019 | *p<0.001* | 0.145 |
| Cognitive_factor | ~ | anteriorthalamicradiations_md | 0.065 | 0.017 | *p<0.001* | 0.085 |
| Cognitive_factor | ~ | uncinate_md | -0.068 | 0.022 | *0.002* | -0.089 |
| Cognitive_factor | ~ | inferiorlongitudinalfasiculus_md | -0.004 | 0.023 | 0.852 | -0.005 |
| Cognitive_factor | ~ | inferiorfrontooccipitalfasiculus_md | 0.065 | 0.027 | *0.018* | 0.081 |
| Cognitive_factor | ~ | forcepsmajor_md | 0.023 | 0.023 | 0.328 | 0.032 |
| Cognitive_factor | ~ | forcepsminor_md | 0.060 | 0.023 | *0.008* | 0.084 |
| Cognitive_factor | ~ | corpuscallosum_md | -0.078 | 0.032 | *0.013* | -0.107 |
| Cognitive_factor | ~ | superiorlongitudinalfasiculus_md | -0.116 | 0.202 | 0.564 | -0.151 |
| Cognitive_factor | ~ | temporalsuperiorlongitudinalfasiculus_md | 0.132 | 0.106 | 0.212 | 0.171 |
| Cognitive_factor | ~ | parietalsuperiorlongitudinalfasiculus_md | -0.062 | 0.116 | 0.592 | -0.080 |
| Cognitive_factor | ~ | superiorcorticostriate_md | -0.009 | 0.164 | 0.957 | -0.011 |
| Cognitive_factor | ~ | superiorcorticostriatefrontalcortex_md | -0.040 | 0.074 | 0.583 | -0.051 |
| Cognitive_factor | ~ | superiorcorticostriateparietalcortex_md | -0.106 | 0.114 | 0.353 | -0.134 |
| Cognitive_factor | ~ | striatalinferiorfrontalcortex_md | -0.061 | 0.027 | *0.023* | -0.074 |
| Cognitive_factor | ~ | inferiorfrontalsuperiorfrontalcortex_md | 0.101 | 0.032 | *0.002* | 0.131 |
| Cognitive_factor | ~ | fornix_exfimbria_md | 0.022 | 0.025 | 0.376 | 0.029 |
| Cognitive_factor | ~ | fornix_wmv | 0.237 | 0.032 | *p<0.001* | 0.322 |

| | | | | | | |
|---|---|---|---|---|---|---|
| Cognitive_factor | ~ | cingulatecingulum_wmv | -0.021 | 0.019 | 0.271 | -0.028 |
| Cognitive_factor | ~ | parahippocampalcingulum_wmv | 0.039 | 0.013 | *0.004* | 0.051 |
| Cognitive_factor | ~ | corticospinalpyramidal_wmv | 0.198 | 0.024 | *p<0.001* | 0.266 |
| Cognitive_factor | ~ | anteriorthalamicradiations_wmv | 0.018 | 0.027 | 0.493 | 0.025 |
| Cognitive_factor | ~ | uncinate_wmv | -0.007 | 0.018 | 0.698 | -0.009 |
| Cognitive_factor | ~ | inferiorlongitudinalfasiculus_wmv | -0.003 | 0.021 | 0.873 | -0.004 |
| Cognitive_factor | ~ | inferiorfrontooccipitalfasiculus_wmv | -0.054 | 0.032 | 0.089 | -0.073 |
| Cognitive_factor | ~ | forcepsmajor_wmv | 0.000 | 0.021 | 0.996 | 0.000 |
| Cognitive_factor | ~ | forcepsminor_wmv | -0.046 | 0.026 | 0.084 | -0.064 |
| Cognitive_factor | ~ | corpuscallosum_wmv | 0.082 | 0.044 | 0.060 | 0.114 |
| Cognitive_factor | ~ | superiorlongitudinalfasiculus_wmv | -0.111 | 0.084 | 0.187 | -0.147 |
| Cognitive_factor | ~ | temporalsuperiorlongitudinalfasiculus_wmv | 0.147 | 0.056 | *0.009* | 0.193 |
| Cognitive_factor | ~ | parietalsuperiorlongitudinalfasiculus_wmv | 0.009 | 0.047 | 0.849 | 0.012 |
| Cognitive_factor | ~ | superiorcorticostriate_wmv | -0.085 | 0.087 | 0.331 | -0.114 |
| Cognitive_factor | ~ | superiorcorticostriatefrontalcortex_wmv | -0.158 | 0.053 | *0.003* | -0.212 |
| Cognitive_factor | ~ | superiorcorticostriateparietalcortex_wmv | -0.008 | 0.057 | 0.885 | -0.011 |
| Cognitive_factor | ~ | striatalinferiorfrontalcortex_wmv | 0.030 | 0.021 | 0.162 | 0.039 |
| Cognitive_factor | ~ | inferiorfrontalsuperiorfrontalcortex_wmv | 0.076 | 0.024 | *0.002* | 0.101 |
| Cognitive_factor | ~ | fornix_exfimbria_wmv | -0.065 | 0.031 | *0.036* | -0.084 |

*Supplementary Table 38. Comparison of the model with free parameters and the model with parameters constrained to a same value for the white matter model with all the regions. AIC/BIC are two information criterions. Pr(>Chisq): p-value of the chi-square ratio test.*

| | **Degree of freedom** | **AIC** | **BIC** | **Difference in degree of freedom** | **Pr (>Chisq)** |
|---|---|---|---|---|---|
| Free model | 245 | 122708.0 | 123244.7 | NA | NA |
| Constrained model | 304 | 123602.8 | 123717.3 | 59 | *p<0.001* |

### White matter metrics (model with the regularized regions)

*Supplementary Table 39. Regression estimates for the model estimating how fractional anisotropy, mean diffusivity and white matter volume of the regularized regions predict the cognitive factor*

| Path | | | Estimate | SE | *p* | Standardized Estimate |
|---|---|---|---|---|---|---|
| Cognitive_factor | ~ | fornix_fa | 0.026 | 0.012 | *0.028* | 0.035 |
| Cognitive_factor | ~ | cingulatecingulum_fa | -0.119 | 0.012 | *p<0.001* | -0.155 |
| Cognitive_factor | ~ | anteriorthalamicradiations_fa | 0.009 | 0.014 | 0.536 | 0.012 |
| Cognitive_factor | ~ | inferiorlongitudinalfasiculus_fa | 0.003 | 0.015 | 0.850 | 0.004 |
| Cognitive_factor | ~ | superiorlongitudinalfasiculus_fa | 0.135 | 0.015 | *p<0.001* | 0.182 |
| Cognitive_factor | ~ | striatalinferiorfrontalcortex_fa | -0.065 | 0.013 | *p<0.001* | -0.086 |
| Cognitive_factor | ~ | inferiorfrontalsuperiorfrontalcortex_fa | 0.049 | 0.016 | *0.002* | 0.066 |
| Cognitive_factor | ~ | anteriorthalamicradiations_md | 0.068 | 0.016 | *p<0.001* | 0.094 |
| Cognitive_factor | ~ | inferiorlongitudinalfasiculus_md | -0.021 | 0.021 | 0.325 | -0.028 |
| Cognitive_factor | ~ | inferiorfrontooccipitalfasiculus_md | 0.013 | 0.024 | 0.589 | 0.017 |
| Cognitive_factor | ~ | forcepsmajor_md | 0.021 | 0.014 | 0.135 | 0.029 |
| Cognitive_factor | ~ | superiorcorticostriateparietalcortex_md | -0.063 | 0.017 | *p<0.001* | -0.086 |
| Cognitive_factor | ~ | fornix_exfimbria_md | -0.016 | 0.012 | 0.184 | -0.021 |
| Cognitive_factor | ~ | corticospinalpyramidal_wmv | 0.144 | 0.018 | *p<0.001* | 0.197 |
| Cognitive_factor | ~ | anteriorthalamicradiations_wmv | -0.008 | 0.023 | 0.721 | -0.012 |
| Cognitive_factor | ~ | uncinate_wmv | -0.017 | 0.016 | 0.296 | -0.023 |
| Cognitive_factor | ~ | inferiorlongitudinalfasiculus_wmv | 0.010 | 0.019 | 0.595 | 0.013 |
| Cognitive_factor | ~ | inferiorfrontooccipitalfasiculus_wmv | -0.002 | 0.028 | 0.942 | -0.003 |
| Cognitive_factor | ~ | forcepsmajor_wmv | 0.049 | 0.019 | *0.008* | 0.069 |
| Cognitive_factor | ~ | forcepsminor_wmv | -0.035 | 0.023 | 0.128 | -0.050 |
| Cognitive_factor | ~ | corpuscallosum_wmv | 0.042 | 0.032 | 0.190 | 0.059 |
| Cognitive_factor | ~ | temporalsuperiorlongitudinalfasiculus_wmv | 0.073 | 0.017 | *p<0.001* | 0.097 |
| Cognitive_factor | ~ | striatalinferiorfrontalcortex_wmv | -0.005 | 0.019 | 0.788 | -0.007 |

*Supplementary Table 40. Standardized parameter estimates of how the white matter metrics of each regularized region of interest together predict the cognitive factor*

| Path | | | FA | MD | WMV |
|---|---|---|---|---|---|
| Cognitive_factor | ~ | corpuscallosum | - | - | 0.059 |
| Cognitive_factor | ~ | forcepsmajor | - | 0.029 | 0.069 |
| Cognitive_factor | ~ | forcepsminor | - | - | -0.050 |
| Cognitive_factor | ~ | anteriorthalamicradiations | 0.012 | 0.094 | -0.012 |
| Cognitive_factor | ~ | cingulatecingulum | -0.155 | - | - |
| Cognitive_factor | ~ | corticospinalpyramidal | - | - | 0.197 |
| Cognitive_factor | ~ | fornix | 0.035 | - | - |
| Cognitive_factor | ~ | fornix_exfimbria | - | -0.021 | - |
| Cognitive_factor | ~ | inferiorfrontalsuperiorfrontalcortex | 0.066 | - | - |
| Cognitive_factor | ~ | inferiorfrontooccipitalfasiculus | - | 0.017 | -0.003 |
| Cognitive_factor | ~ | inferiorlongitudinalfasiculus | 0.004 | -0.028 | 0.013 |
| Cognitive_factor | ~ | striatalinferiorfrontalcortex | -0.086 | - | -0.007 |
| Cognitive_factor | ~ | superiorcorticostriateparietalcortex | - | -0.09 | - |
| Cognitive_factor | ~ | superiorlongitudinalfasiculus | 0.182 | - | - |
| Cognitive_factor | ~ | temporalsuperiorlongitudinalfasiculus | - | - | 0.097 |
| Cognitive_factor | ~ | uncinate | - | - | -0.023 |

*Supplementary Table 41. Comparison of the model with free parameters and the model with parameters constrained to a same value for the white matter model with the regularized regions. AIC/BIC are two information criterions. Pr(>Chisq): p-value of the chi-square ratio test.*

| | Degree of freedom | AIC | BIC | Difference in degree of freedom | Pr (>Chisq) |
|---|---|---|---|---|---|
| Free model | 97 | 123117.4 | 123389.4 | NA | NA |
| Constrained model | 119 | 123601.0 | 123715.5 | 22 | *p<0.001* |

### Models with grey and white matter metrics estimating cognitive factor from several regions in the six metrics

For the grey and white matter model, we reported the table for the model with all the regions and the model with only the regions that survived the regularization included as predictors (the one that we used for the analyses).

#### Grey and white matter metrics (model with all the regions)

*Supplementary Table 42. Regression estimates for the model estimating how cortical thickness, surface area, grey matter volume, fractional anisotropy, mean diffusivity and white matter volume of all the regions predict the cognitive factor*

| Path | | | Estimate | SE | *p* | Standardized Estimate |
|---|---|---|---|---|---|---|
| Cognitive_factor | ~ | bankssts_ct | 0.002 | 0.018 | 0.916 | 0.002 |
| Cognitive_factor | ~ | caudalanteriorcingulate_ct | -0.027 | 0.018 | 0.136 | -0.032 |
| Cognitive_factor | ~ | caudalmiddlefrontal_ct | 0.015 | 0.030 | 0.622 | 0.018 |
| Cognitive_factor | ~ | cuneus_ct | 0.007 | 0.031 | 0.812 | 0.009 |
| Cognitive_factor | ~ | entorhinal_ct | 0.014 | 0.020 | 0.485 | 0.016 |
| Cognitive_factor | ~ | fusiform_ct | -0.003 | 0.026 | 0.920 | -0.003 |
| Cognitive_factor | ~ | inferiorparietal_ct | -0.022 | 0.032 | 0.486 | -0.028 |
| Cognitive_factor | ~ | inferiortemporal_ct | 0.042 | 0.030 | 0.152 | 0.052 |
| Cognitive_factor | ~ | isthmuscingulate_ct | 0.022 | 0.020 | 0.283 | 0.026 |
| Cognitive_factor | ~ | lateraloccipital_ct | -0.007 | 0.038 | 0.845 | -0.010 |
| Cognitive_factor | ~ | lateralorbitofrontal_ct | -0.076 | 0.023 | *0.001* | -0.092 |
| Cognitive_factor | ~ | lingual_ct | 0.061 | 0.032 | 0.055 | 0.078 |
| Cognitive_factor | ~ | medialorbitofrontal_ct | 0.040 | 0.024 | 0.097 | 0.047 |
| Cognitive_factor | ~ | middletemporal_ct | -0.033 | 0.034 | 0.335 | -0.041 |
| Cognitive_factor | ~ | parahippocampal_ct | 0.028 | 0.029 | 0.331 | 0.035 |
| Cognitive_factor | ~ | paracentral_ct | 0.015 | 0.028 | 0.581 | 0.019 |
| Cognitive_factor | ~ | parsopercularis_ct | -0.038 | 0.027 | 0.163 | -0.045 |
| Cognitive_factor | ~ | parsorbitalis_ct | 0.006 | 0.024 | 0.806 | 0.007 |
| Cognitive_factor | ~ | parstriangularis_ct | -0.050 | 0.027 | 0.068 | -0.059 |
| Cognitive_factor | ~ | pericalcarine_ct | 0.049 | 0.030 | 0.096 | 0.063 |
| Cognitive_factor | ~ | postcentral_ct | 0.013 | 0.037 | 0.735 | 0.016 |
| Cognitive_factor | ~ | posteriorcingulate_ct | -0.029 | 0.023 | 0.206 | -0.034 |
| Cognitive_factor | ~ | precentral_ct | 0.166 | 0.039 | *p<0.001* | 0.208 |
| Cognitive_factor | ~ | precuneus_ct | -0.053 | 0.026 | *0.045* | -0.067 |
| Cognitive_factor | ~ | rostralanteriorcingulate_ct | -0.038 | 0.020 | 0.059 | -0.043 |
| Cognitive_factor | ~ | rostralmiddlefrontal_ct | 0.115 | 0.031 | *p<0.001* | 0.144 |
| Cognitive_factor | ~ | superiorfrontal_ct | -0.007 | 0.033 | 0.827 | -0.009 |
| Cognitive_factor | ~ | superiorparietal_ct | -0.061 | 0.036 | 0.084 | -0.080 |
| Cognitive_factor | ~ | superiortemporal_ct | -0.072 | 0.036 | *0.043* | -0.091 |
| Cognitive_factor | ~ | supramarginal_ct | 0.052 | 0.032 | 0.101 | 0.065 |
| Cognitive_factor | ~ | frontalpole_ct | 0.004 | 0.022 | 0.868 | 0.004 |

| | | | | | | |
|---|---|---|---|---|---|---|
| Cognitive_factor | ~ | temporalpole_ct | -0.038 | 0.022 | 0.077 | -0.045 |
| Cognitive_factor | ~ | transversetemporal_ct | -0.043 | 0.020 | *0.027* | -0.052 |
| Cognitive_factor | ~ | insula_ct | -0.023 | 0.028 | 0.399 | -0.028 |
| Cognitive_factor | ~ | bankssts_sa | -0.059 | 0.046 | 0.202 | -0.070 |
| Cognitive_factor | ~ | caudalanteriorcingulate_sa | 0.018 | 0.048 | 0.703 | 0.020 |
| Cognitive_factor | ~ | caudalmiddlefrontal_sa | 0.069 | 0.073 | 0.343 | 0.086 |
| Cognitive_factor | ~ | cuneus_sa | -0.035 | 0.049 | 0.472 | -0.046 |
| Cognitive_factor | ~ | entorhinal_sa | 0.009 | 0.033 | 0.784 | 0.011 |
| Cognitive_factor | ~ | fusiform_sa | 0.061 | 0.057 | 0.289 | 0.079 |
| Cognitive_factor | ~ | inferiorparietal_sa | 0.056 | 0.070 | 0.421 | 0.072 |
| Cognitive_factor | ~ | inferiortemporal_sa | 0.231 | 0.065 | *p<0.001* | 0.299 |
| Cognitive_factor | ~ | isthmuscingulate_sa | -0.001 | 0.053 | 0.978 | -0.002 |
| Cognitive_factor | ~ | lateraloccipital_sa | -0.143 | 0.068 | *0.034* | -0.186 |
| Cognitive_factor | ~ | lateralorbitofrontal_sa | -0.108 | 0.055 | *0.048* | -0.142 |
| Cognitive_factor | ~ | lingual_sa | 0.113 | 0.060 | 0.057 | 0.149 |
| Cognitive_factor | ~ | medialorbitofrontal_sa | 0.169 | 0.051 | *0.001* | 0.215 |
| Cognitive_factor | ~ | middletemporal_sa | -0.168 | 0.063 | *0.008* | -0.222 |
| Cognitive_factor | ~ | parahippocampal_sa | -0.060 | 0.039 | 0.129 | -0.074 |
| Cognitive_factor | ~ | paracentral_sa | 0.070 | 0.055 | 0.205 | 0.086 |
| Cognitive_factor | ~ | parsopercularis_sa | 0.151 | 0.071 | *0.032* | 0.184 |
| Cognitive_factor | ~ | parsorbitalis_sa | -0.019 | 0.046 | 0.683 | -0.024 |
| Cognitive_factor | ~ | parstriangularis_sa | -0.084 | 0.067 | 0.212 | -0.103 |
| Cognitive_factor | ~ | pericalcarine_sa | 0.123 | 0.052 | *0.018* | 0.164 |
| Cognitive_factor | ~ | postcentral_sa | -0.042 | 0.070 | 0.551 | -0.054 |
| Cognitive_factor | ~ | posteriorcingulate_sa | -0.097 | 0.073 | 0.184 | -0.116 |
| Cognitive_factor | ~ | precentral_sa | 0.149 | 0.070 | *0.032* | 0.192 |
| Cognitive_factor | ~ | precuneus_sa | -0.023 | 0.066 | 0.727 | -0.031 |
| Cognitive_factor | ~ | rostralanteriorcingulate_sa | -0.118 | 0.050 | *0.018* | -0.144 |
| Cognitive_factor | ~ | rostralmiddlefrontal_sa | 0.261 | 0.068 | *p<0.001* | 0.339 |
| Cognitive_factor | ~ | superiorfrontal_sa | -0.007 | 0.078 | 0.932 | -0.009 |
| Cognitive_factor | ~ | superiorparietal_sa | 0.012 | 0.069 | 0.862 | 0.016 |
| Cognitive_factor | ~ | superiortemporal_sa | -0.095 | 0.066 | 0.150 | -0.124 |
| Cognitive_factor | ~ | supramarginal_sa | 0.050 | 0.074 | 0.497 | 0.063 |
| Cognitive_factor | ~ | frontalpole_sa | 0.022 | 0.028 | 0.432 | 0.026 |
| Cognitive_factor | ~ | temporalpole_sa | -0.120 | 0.027 | *p<0.001* | -0.148 |
| Cognitive_factor | ~ | transversetemporal_sa | -0.122 | 0.034 | *p<0.001* | -0.151 |
| Cognitive_factor | ~ | insula_sa | -0.057 | 0.056 | 0.303 | -0.074 |
| Cognitive_factor | ~ | bankssts_gmv | 0.033 | 0.047 | 0.485 | 0.039 |
| Cognitive_factor | ~ | caudalanteriorcingulate_gmv | 0.014 | 0.050 | 0.785 | 0.014 |
| Cognitive_factor | ~ | caudalmiddlefrontal_gmv | -0.041 | 0.070 | 0.561 | -0.050 |
| Cognitive_factor | ~ | cuneus_gmv | 0.021 | 0.058 | 0.721 | 0.027 |
| Cognitive_factor | ~ | entorhinal_gmv | 0.009 | 0.030 | 0.755 | 0.012 |
| Cognitive_factor | ~ | fusiform_gmv | -0.030 | 0.056 | 0.590 | -0.038 |
| Cognitive_factor | ~ | inferiorparietal_gmv | -0.067 | 0.068 | 0.321 | -0.085 |

| | | | | | | |
|---|---|---|---|---|---|---|
| Cognitive_factor | ~ | inferiortemporal_gmv | -0.160 | 0.063 | *0.011* | -0.204 |
| Cognitive_factor | ~ | isthmuscingulate_gmv | -0.055 | 0.052 | 0.290 | -0.068 |
| Cognitive_factor | ~ | lateraloccipital_gmv | 0.161 | 0.072 | *0.026* | 0.209 |
| Cognitive_factor | ~ | lateralorbitofrontal_gmv | 0.150 | 0.052 | *0.004* | 0.198 |
| Cognitive_factor | ~ | lingual_gmv | -0.135 | 0.066 | *0.041* | -0.175 |
| Cognitive_factor | ~ | medialorbitofrontal_gmv | -0.161 | 0.052 | *0.002* | -0.201 |
| Cognitive_factor | ~ | middletemporal_gmv | 0.196 | 0.057 | *0.001* | 0.257 |
| Cognitive_factor | ~ | parahippocampal_gmv | 0.058 | 0.043 | 0.176 | 0.071 |
| Cognitive_factor | ~ | paracentral_gmv | -0.077 | 0.055 | 0.164 | -0.093 |
| Cognitive_factor | ~ | parsopercularis_gmv | -0.114 | 0.067 | 0.090 | -0.137 |
| Cognitive_factor | ~ | parsorbitalis_gmv | 0.038 | 0.043 | 0.372 | 0.047 |
| Cognitive_factor | ~ | parstriangularis_gmv | 0.048 | 0.064 | 0.449 | 0.058 |
| Cognitive_factor | ~ | pericalcarine_gmv | -0.150 | 0.058 | *0.010* | -0.199 |
| Cognitive_factor | ~ | postcentral_gmv | -0.017 | 0.071 | 0.809 | -0.022 |
| Cognitive_factor | ~ | posteriorcingulate_gmv | 0.065 | 0.069 | 0.351 | 0.077 |
| Cognitive_factor | ~ | precentral_gmv | -0.109 | 0.068 | 0.107 | -0.140 |
| Cognitive_factor | ~ | precuneus_gmv | 0.014 | 0.065 | 0.830 | 0.018 |
| Cognitive_factor | ~ | rostralanteriorcingulate_gmv | 0.119 | 0.048 | *0.014* | 0.139 |
| Cognitive_factor | ~ | rostralmiddlefrontal_gmv | -0.245 | 0.065 | *p<0.001* | -0.315 |
| Cognitive_factor | ~ | superiorfrontal_gmv | 0.056 | 0.069 | 0.418 | 0.073 |
| Cognitive_factor | ~ | superiorparietal_gmv | 0.011 | 0.069 | 0.873 | 0.014 |
| Cognitive_factor | ~ | superiortemporal_gmv | 0.147 | 0.063 | *0.019* | 0.190 |
| Cognitive_factor | ~ | supramarginal_gmv | -0.070 | 0.069 | 0.311 | -0.087 |
| Cognitive_factor | ~ | frontalpole_gmv | -0.007 | 0.029 | 0.820 | -0.008 |
| Cognitive_factor | ~ | temporalpole_gmv | 0.097 | 0.030 | *0.001* | 0.116 |
| Cognitive_factor | ~ | transversetemporal_gmv | 0.110 | 0.033 | *0.001* | 0.136 |
| Cognitive_factor | ~ | insula_gmv | 0.084 | 0.055 | 0.127 | 0.111 |
| Cognitive_factor | ~ | fornix_fa | 0.019 | 0.019 | 0.332 | 0.024 |
| Cognitive_factor | ~ | cingulatecingulum_fa | -0.044 | 0.016 | *0.006* | -0.056 |
| Cognitive_factor | ~ | parahippocampalcingulum_fa | -0.001 | 0.013 | 0.955 | -0.001 |
| Cognitive_factor | ~ | corticospinalpyramidal_fa | 0.048 | 0.020 | *0.015* | 0.062 |
| Cognitive_factor | ~ | anteriorthalamicradiations_fa | 0.034 | 0.017 | *0.048* | 0.043 |
| Cognitive_factor | ~ | uncinate_fa | -0.056 | 0.020 | *0.005* | -0.071 |
| Cognitive_factor | ~ | inferiorlongitudinalfasiculus_fa | 0.015 | 0.019 | 0.413 | 0.019 |
| Cognitive_factor | ~ | inferiorfrontooccipitalfasiculus_fa | 0.071 | 0.024 | *0.003* | 0.092 |
| Cognitive_factor | ~ | forcepsmajor_fa | 0.020 | 0.023 | 0.375 | 0.028 |
| Cognitive_factor | ~ | forcepsminor_fa | -0.026 | 0.024 | 0.289 | -0.036 |
| Cognitive_factor | ~ | corpuscallosum_fa | -0.086 | 0.040 | *0.030* | -0.118 |
| Cognitive_factor | ~ | superiorlongitudinalfasiculus_fa | -0.076 | 0.142 | 0.594 | -0.099 |
| Cognitive_factor | ~ | temporalsuperiorlongitudinalfasiculus_fa | 0.040 | 0.077 | 0.602 | 0.052 |
| Cognitive_factor | ~ | parietalsuperiorlongitudinalfasiculus_fa | 0.124 | 0.084 | 0.142 | 0.161 |
| Cognitive_factor | ~ | superiorcorticostriate_fa | 0.365 | 0.132 | *0.006* | 0.469 |
| Cognitive_factor | ~ | superiorcorticostriatefrontalcortex_fa | -0.088 | 0.059 | 0.132 | -0.114 |
| Cognitive_factor | ~ | superiorcorticostriateparietalcortex_fa | -0.320 | 0.091 | *p<0.001* | -0.407 |

| | | | | | | |
|---|---|---|---|---|---|---|
| Cognitive_factor | ~ | striatalinferiorfrontalcortex_fa | -0.047 | 0.018 | *0.008* | -0.059 |
| Cognitive_factor | ~ | inferiorfrontalsuperiorfrontalcortex_fa | 0.055 | 0.022 | *0.013* | 0.071 |
| Cognitive_factor | ~ | fornix_exfimbria_fa | -0.050 | 0.018 | *0.005* | -0.062 |
| Cognitive_factor | ~ | fornix_md | -0.016 | 0.024 | 0.497 | -0.021 |
| Cognitive_factor | ~ | cingulatecingulum_md | 0.007 | 0.016 | 0.654 | 0.008 |
| Cognitive_factor | ~ | parahippocampalcingulum_md | -0.026 | 0.015 | 0.085 | -0.034 |
| Cognitive_factor | ~ | corticospinalpyramidal_md | 0.096 | 0.019 | *p<0.001* | 0.124 |
| Cognitive_factor | ~ | anteriorthalamicradiations_md | 0.041 | 0.017 | *0.016* | 0.054 |
| Cognitive_factor | ~ | uncinate_md | -0.042 | 0.021 | *0.043* | -0.055 |
| Cognitive_factor | ~ | inferiorlongitudinalfasiculus_md | -0.006 | 0.023 | 0.801 | -0.007 |
| Cognitive_factor | ~ | inferiorfrontooccipitalfasiculus_md | 0.031 | 0.027 | 0.247 | 0.039 |
| Cognitive_factor | ~ | forcepsmajor_md | 0.060 | 0.021 | *0.004* | 0.083 |
| Cognitive_factor | ~ | forcepsminor_md | 0.067 | 0.023 | *0.003* | 0.093 |
| Cognitive_factor | ~ | corpuscallosum_md | -0.120 | 0.032 | *p<0.001* | -0.163 |
| Cognitive_factor | ~ | superiorlongitudinalfasiculus_md | -0.166 | 0.204 | 0.415 | -0.214 |
| Cognitive_factor | ~ | temporalsuperiorlongitudinalfasiculus_md | 0.149 | 0.107 | 0.163 | 0.192 |
| Cognitive_factor | ~ | parietalsuperiorlongitudinalfasiculus_md | -0.023 | 0.116 | 0.844 | -0.029 |
| Cognitive_factor | ~ | superiorcorticostriate_md | 0.119 | 0.160 | 0.457 | 0.149 |
| Cognitive_factor | ~ | superiorcorticostriatefrontalcortex_md | -0.147 | 0.073 | *0.044* | -0.185 |
| Cognitive_factor | ~ | superiorcorticostriateparietalcortex_md | -0.123 | 0.111 | 0.266 | -0.155 |
| Cognitive_factor | ~ | striatalinferiorfrontalcortex_md | -0.035 | 0.026 | 0.171 | -0.043 |
| Cognitive_factor | ~ | inferiorfrontalsuperiorfrontalcortex_md | 0.134 | 0.032 | *p<0.001* | 0.172 |
| Cognitive_factor | ~ | fornix_exfimbria_md | 0.022 | 0.024 | 0.365 | 0.029 |
| Cognitive_factor | ~ | fornix_wmv | 0.134 | 0.034 | *p<0.001* | 0.182 |
| Cognitive_factor | ~ | cingulatecingulum_wmv | -0.006 | 0.021 | 0.776 | -0.008 |
| Cognitive_factor | ~ | parahippocampalcingulum_wmv | 0.041 | 0.014 | *0.004* | 0.053 |
| Cognitive_factor | ~ | corticospinalpyramidal_wmv | 0.194 | 0.024 | *p<0.001* | 0.260 |
| Cognitive_factor | ~ | anteriorthalamicradiations_wmv | 0.028 | 0.028 | 0.306 | 0.038 |
| Cognitive_factor | ~ | uncinate_wmv | -0.001 | 0.019 | 0.972 | -0.001 |
| Cognitive_factor | ~ | inferiorlongitudinalfasiculus_wmv | -0.010 | 0.022 | 0.662 | -0.013 |
| Cognitive_factor | ~ | inferiorfrontooccipitalfasiculus_wmv | -0.073 | 0.032 | *0.025* | -0.097 |
| Cognitive_factor | ~ | forcepsmajor_wmv | 0.011 | 0.022 | 0.601 | 0.016 |
| Cognitive_factor | ~ | forcepsminor_wmv | -0.108 | 0.028 | *p<0.001* | -0.150 |
| Cognitive_factor | ~ | corpuscallosum_wmv | 0.094 | 0.046 | *0.038* | 0.131 |
| Cognitive_factor | ~ | superiorlongitudinalfasiculus_wmv | -0.110 | 0.084 | 0.188 | -0.146 |
| Cognitive_factor | ~ | temporalsuperiorlongitudinalfasiculus_wmv | 0.125 | 0.056 | *0.025* | 0.164 |
| Cognitive_factor | ~ | parietalsuperiorlongitudinalfasiculus_wmv | 0.026 | 0.048 | 0.588 | 0.034 |
| Cognitive_factor | ~ | superiorcorticostriate_wmv | 0.052 | 0.087 | 0.551 | 0.069 |
| Cognitive_factor | ~ | superiorcorticostriatefrontalcortex_wmv | -0.219 | 0.053 | *p<0.001* | -0.292 |
| Cognitive_factor | ~ | superiorcorticostriateparietalcortex_wmv | -0.070 | 0.057 | 0.218 | -0.092 |
| Cognitive_factor | ~ | striatalinferiorfrontalcortex_wmv | 0.006 | 0.022 | 0.804 | 0.007 |
| Cognitive_factor | ~ | inferiorfrontalsuperiorfrontalcortex_wmv | 0.005 | 0.026 | 0.851 | 0.007 |
| Cognitive_factor | ~ | fornix_exfimbria_wmv | -0.005 | 0.032 | 0.880 | -0.006 |

*Supplementary Table 43. Comparison of the model with free parameters and the model with parameters constrained to a same value for the grey & white matter model with all the regions. AIC/BIC are two information criterions. Pr(>Chisq): p-value of the chi-square ratio test.*

| | **Degree of freedom** | **AIC** | **BIC** | **Difference in degree of freedom** | **Pr (>Chisq)** |
|---|---|---|---|---|---|
| Free model | 653 | 122276 | 123543 | NA | NA |
| Constrained model | 814 | 123378 | 123492 | 161 | *p<0.001* |

### Grey and white matter metrics (model with the regularized regions)

*Supplementary Table 44. Regression estimates for the model estimating how cortical thickness, surface area, grey matter volume, fractional anisotropy, mean diffusivity and white matter volume of the regularized regions predict the cognitive factor*

| **Path** | | | **Estimate** | **SE** | ***p*** | **Standardized Estimate** |
|---|---|---|---|---|---|---|
| Cognitive_factor | ~ | caudalanteriorcingulate_ct | -0.032 | 0.012 | *0.008* | -0.037 |
| Cognitive_factor | ~ | entorhinal_ct | 0.013 | 0.010 | 0.208 | 0.016 |
| Cognitive_factor | ~ | lateraloccipital_ct | 0.067 | 0.015 | *p<0.001* | 0.089 |
| Cognitive_factor | ~ | lingual_ct | 0.001 | 0.011 | 0.911 | 0.002 |
| Cognitive_factor | ~ | medialorbitofrontal_ct | -0.031 | 0.012 | *0.008* | -0.038 |
| Cognitive_factor | ~ | middletemporal_ct | 0.013 | 0.014 | 0.365 | 0.016 |
| Cognitive_factor | ~ | parahippocampal_ct | 0.094 | 0.012 | *p<0.001* | 0.120 |
| Cognitive_factor | ~ | parsopercularis_ct | -0.077 | 0.014 | *p<0.001* | -0.092 |
| Cognitive_factor | ~ | parstriangularis_ct | -0.030 | 0.015 | *0.047* | -0.036 |
| Cognitive_factor | ~ | precentral_ct | 0.086 | 0.015 | *p<0.001* | 0.110 |
| Cognitive_factor | ~ | precuneus_ct | -0.033 | 0.015 | *0.024* | -0.043 |
| Cognitive_factor | ~ | rostralanteriorcingulate_ct | -0.018 | 0.013 | 0.163 | -0.021 |
| Cognitive_factor | ~ | rostralmiddlefrontal_ct | 0.019 | 0.014 | 0.186 | 0.024 |
| Cognitive_factor | ~ | superiorparietal_ct | -0.065 | 0.016 | *p<0.001* | -0.085 |
| Cognitive_factor | ~ | transversetemporal_ct | 0.026 | 0.011 | *0.022* | 0.031 |
| Cognitive_factor | ~ | insula_ct | -0.007 | 0.012 | 0.558 | -0.009 |
| Cognitive_factor | ~ | caudalmiddlefrontal_sa | 0.037 | 0.014 | *0.009* | 0.047 |
| Cognitive_factor | ~ | cuneus_sa | -0.011 | 0.016 | 0.513 | -0.014 |
| Cognitive_factor | ~ | isthmuscingulate_sa | -0.067 | 0.014 | *p<0.001* | -0.085 |
| Cognitive_factor | ~ | parsopercularis_sa | 0.042 | 0.014 | *0.002* | 0.051 |
| Cognitive_factor | ~ | pericalcarine_sa | 0.006 | 0.015 | 0.691 | 0.008 |
| Cognitive_factor | ~ | posteriorcingulate_sa | 0.049 | 0.052 | 0.348 | 0.059 |
| Cognitive_factor | ~ | superiorparietal_sa | 0.014 | 0.012 | 0.263 | 0.018 |
| Cognitive_factor | ~ | entorhinal_gmv | 0.023 | 0.011 | *0.040* | 0.029 |

| | | | | | | |
|---|---|---|---|---|---|---|
| Cognitive_factor | ~ | middletemporal_gmv | 0.049 | 0.013 | *p<0.001* | 0.064 |
| Cognitive_factor | ~ | parahippocampal_gmv | 0.099 | 0.014 | *p<0.001* | 0.132 |
| Cognitive_factor | ~ | parstriangularis_gmv | 0.003 | 0.013 | 0.842 | 0.003 |
| Cognitive_factor | ~ | posteriorcingulate_gmv | -0.007 | 0.013 | 0.597 | -0.009 |
| Cognitive_factor | ~ | precentral_gmv | -0.076 | 0.046 | 0.097 | -0.092 |
| Cognitive_factor | ~ | rostralanteriorcingulate_gmv | 0.016 | 0.015 | 0.287 | 0.021 |
| Cognitive_factor | ~ | superiortemporal_gmv | 0.040 | 0.015 | *0.007* | 0.047 |
| Cognitive_factor | ~ | fornix_fa | 0.011 | 0.014 | 0.408 | 0.015 |
| Cognitive_factor | ~ | cingulatecingulum_fa | 0.003 | 0.012 | 0.783 | 0.004 |
| Cognitive_factor | ~ | anteriorthalamicradiations_fa | -0.065 | 0.013 | *p<0.001* | -0.084 |
| Cognitive_factor | ~ | inferiorlongitudinalfasiculus_fa | -0.017 | 0.013 | 0.198 | -0.023 |
| Cognitive_factor | ~ | superiorlongitudinalfasiculus_fa | -0.002 | 0.014 | 0.883 | -0.003 |
| Cognitive_factor | ~ | superiorcorticostriatefrontalcortex_fa | 0.083 | 0.015 | *p<0.001* | 0.112 |
| Cognitive_factor | ~ | inferiorfrontalsuperiorfrontalcortex_fa | 0.024 | 0.014 | 0.093 | 0.032 |
| Cognitive_factor | ~ | cingulatecingulum_md | 0.003 | 0.017 | 0.848 | 0.004 |
| Cognitive_factor | ~ | uncinate_md | 0.014 | 0.015 | 0.361 | 0.016 |
| Cognitive_factor | ~ | inferiorfrontooccipitalfasiculus_md | 0.015 | 0.019 | 0.445 | 0.020 |
| Cognitive_factor | ~ | superiorlongitudinalfasiculus_md | 0.038 | 0.019 | *0.046* | 0.051 |
| Cognitive_factor | ~ | superiorcorticostriateparietalcortex_md | -0.030 | 0.022 | 0.168 | -0.041 |
| Cognitive_factor | ~ | corticospinalpyramidal_wmv | -0.042 | 0.020 | *0.033* | -0.057 |
| Cognitive_factor | ~ | uncinate_wmv | 0.233 | 0.023 | *p<0.001* | 0.318 |
| Cognitive_factor | ~ | forcepsminor_wmv | -0.023 | 0.015 | 0.134 | -0.031 |
| Cognitive_factor | ~ | corpuscallosum_wmv | -0.111 | 0.021 | *p<0.001* | -0.157 |
| Cognitive_factor | ~ | superiorcorticostriatefrontalcortex_wmv | 0.162 | 0.025 | *p<0.001* | 0.229 |
| Cognitive_factor | ~ | striatalinferiorfrontalcortex_wmv | -0.204 | 0.028 | *p<0.001* | -0.278 |

*Supplementary Table 45. Standardized parameter estimates of how the grey & white matter metrics of each regularized region of interest together predict the cognitive factor*

| Path | | | CT | SA | GMV | FA | MD | WMV |
|---|---|---|---|---|---|---|---|---|
| Cognitive_factor | ~ | caudalanteriorcingulate | -0.037 | - | - | | | |
| Cognitive_factor | ~ | caudalmiddlefrontal | - | 0.047 | - | | | |
| Cognitive_factor | ~ | cuneus | - | -0.014 | - | | | |
| Cognitive_factor | ~ | entorhinal | 0.016 | - | 0.029 | | | |
| Cognitive_factor | ~ | fusiform | - | - | 0.064 | | | |
| Cognitive_factor | ~ | insula | -0.009 | - | - | | | |
| Cognitive_factor | ~ | isthmuscingulate | - | -0.085 | - | | | |
| Cognitive_factor | ~ | lateraloccipital | 0.089 | - | - | | | |
| Cognitive_factor | ~ | lingual | 0.002 | - | - | | | |
| Cognitive_factor | ~ | medialorbitofrontal | -0.038 | - | - | | | |
| Cognitive_factor | ~ | middletemporal | 0.016 | - | 0.132 | | | |
| Cognitive_factor | ~ | parahippocampal | 0.120 | - | 0.003 | | | |
| Cognitive_factor | ~ | parsopercularis | -0.092 | 0.051 | - | | | |

| | | | | | | | | |
|---|---|---|---|---|---|---|---|---|
| Cognitive_factor | ~ | parstriangularis | -0.036 | - | -0.009 | | | |
| Cognitive_factor | ~ | pericalcarine | - | 0.008 | - | | | |
| Cognitive_factor | ~ | posteriorcingulate | - | 0.059 | -0.092 | | | |
| Cognitive_factor | ~ | precentral | 0.110 | - | 0.021 | | | |
| Cognitive_factor | ~ | precuneus | -0.043 | - | - | | | |
| Cognitive_factor | ~ | rostralanteriorcingulate | -0.021 | - | 0.047 | | | |
| Cognitive_factor | ~ | rostralmiddlefrontal | 0.024 | - | - | | | |
| Cognitive_factor | ~ | superiorparietal | -0.085 | 0.018 | - | | | |
| Cognitive_factor | ~ | superiortemporal | - | - | 0.015 | | | |
| Cognitive_factor | ~ | transversetemporal | 0.031 | - | - | | | |
| Cognitive_factor | ~ | corpuscallosum | | | | - | - | 0.229 |
| Cognitive_factor | ~ | forcepsminor | | | | - | - | -0.157 |
| Cognitive_factor | ~ | anteriorthalamicradiations | | | | -0.023 | - | - |
| Cognitive_factor | ~ | cingulatecingulum | | | | -0.084 | 0.016 | - |
| Cognitive_factor | ~ | corticospinalpyramidal | | | | - | - | 0.318 |
| Cognitive_factor | ~ | fornix | | | | 0.004 | - | - |
| Cognitive_factor | ~ | inferiorfrontalsuperiorfrontalcortex | | | | 0.004 | - | - |
| Cognitive_factor | ~ | inferiorfrontooccipitalfasiculus | | | | - | 0.051 | - |
| Cognitive_factor | ~ | inferiorlongitudinalfasiculus | | | | -0.003 | - | - |
| Cognitive_factor | ~ | striatalinferiorfrontalcortex | | | | - | - | 0.008 |
| Cognitive_factor | ~ | superiorcorticostriatefrontalcortex | | | | 0.032 | - | -0.278 |
| Cognitive_factor | ~ | superiorcorticostriateparietalcortex | | | | - | -0.057 | - |
| Cognitive_factor | ~ | superiorlongitudinalfasiculus | | | | 0.112 | -0.041 | - |
| Cognitive_factor | ~ | superiorlongitudinalfasiculus | | | | - | 0.020 | -0.031 |
| Cognitive_factor | ~ | uncinate | | | | - | - | 0.229 |

*Supplementary Table 46. Comparison of the model with free parameters and the model with parameters constrained to a same value for the grey & white matter model with the regularized regions. AIC/BIC are two information criterions. Pr(>Chisq): p-value of the chi-square ratio test.*

| | **Degree of freedom** | **AIC** | **BIC** | **Difference in degree of freedom** | **Pr (>Chisq)** |
|---|---|---|---|---|---|
| Free model | 205 | 122675.3 | 123140.5 | NA | NA |
| Constrained model | 254 | 123490.5 | 123605.0 | 49 | *p<0.001* |

### Comparison between models with grey & white matter metrics and with grey & white matter metrics & TIV

We compare the parameter estimates of each region/tract between a model with grey and white matter metrics and a model with grey and white matter metrics and TIV.

*Supplementary Figure 15. Standardized parameter estimates comparison of how the grey matter metrics (top) and the white matter metrics (bottom) predict the cognitive factor in a model with and without TIV*

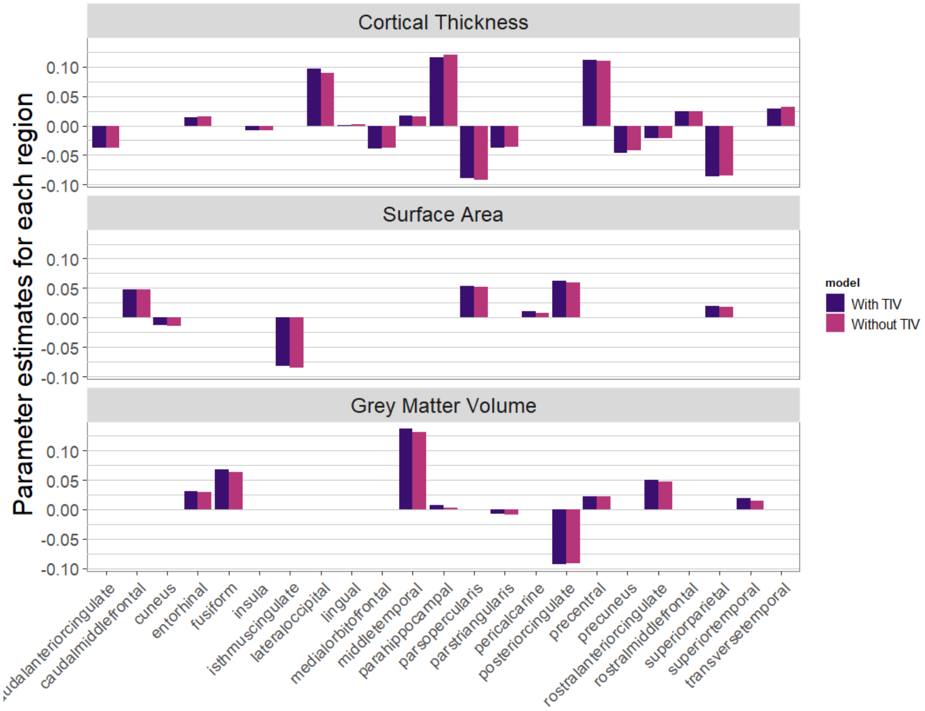


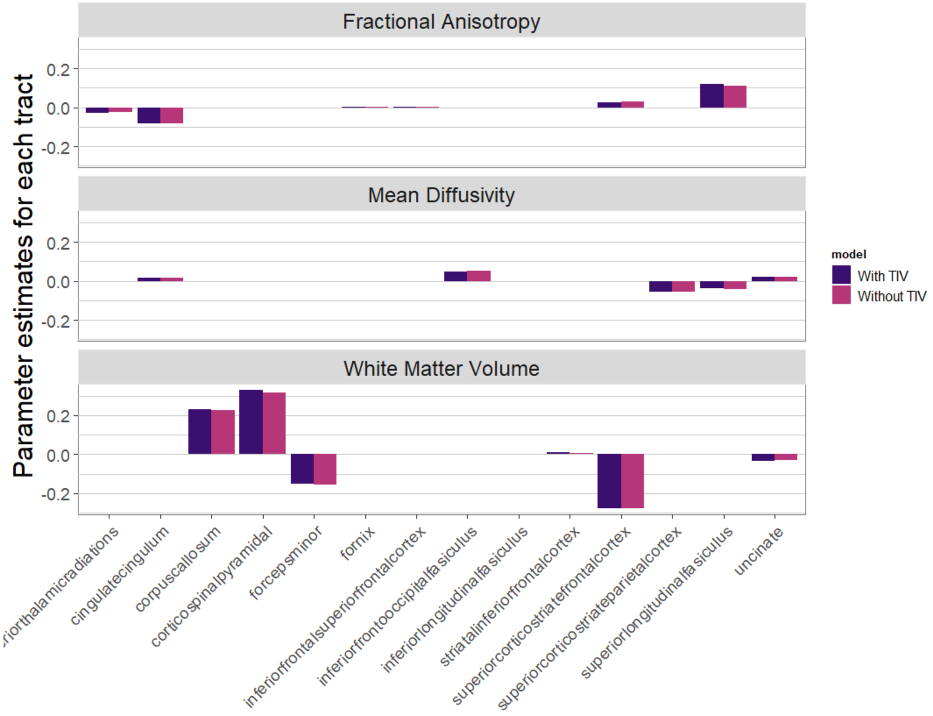
